# Genomic evidence for hybridization and introgression between blue peafowl and green peafowl and selection for white plumage

**DOI:** 10.1101/2023.12.27.573425

**Authors:** Gang Wang, Liping Ban, Xinye Zhang, Xiurong Zhao, Xufang Ren, Anqi Chen, Li Zhang, Yan Lu, Zhihua Jiang, Xiaoyu Zhao, Junhui Wen, Yalan Zhang, Xue Cheng, Huie Wang, Wenting Dai, Yong Liu, Zhonghua Ning, Lujiang Qu

**Affiliations:** National Engineering Laboratory for Animal Breeding and Key Laboratory of Animal Genetics, Breeding and Reproduction, Ministry of Agriculture and Rural Affairs, College of Animal Science and Technology, China Agricultural University, Beijing, China; College of Grassland Science and Technology, China Agricultural University, Beijing, China; Beijing Key Laboratory of Captive Wildlife Technologies, Beijing Zoo, Beijing, China; Institute of Animal Husbandry and Veterinary Medicine, Beijing Academy of Agricultural and Forestry Sciences, Beijing, China Beijing; Department of Animal Sciences, Washington State University, Pullman, USA; School of Animal Science and technology, Tarim University, Xinjiang, China; Nongxiao Breeding Poultry Breeding Co., Ltd. Beijing, China; Xingrui Technology Co., Ltd. Hebei, China

**Keywords:** Peafowl, Hybridization, Introgression, Conservation, White plumage

## Abstract

The blue peafowl (*Pavo cristatus*) and the green peafowl (*Pavo muticus*) have significant public affection due to their stunning appearance, although the green peafowl is currently endangered. Some studies have suggested introgression between these the two species, although evidence is mixed. In this study, we successfully assembled a high-quality chromosome-level reference genome of the blue peafowl, including the autosomes, Z and W sex chromosomes as well as a complete mitochondria DNA sequence. Data from 77 peafowl whole genomes, 76 peafowl mitochondrial genomes and 33 peahen W chromosomes genomes provide the first substantial genetic evidence for recent hybridization between green and blue peafowl. We found three hybrid green peafowls in zoo samples rather than in the wild samples, with blue peafowl genomic content of 16-34%. Maternal genetic analysis showed that two of the hybrid female green peafowls contained complete blue peafowl mitochondrial genomes and W chromosomes. Hybridization of endangered species with its relatives is extremely detrimental to conservation. Some animal protection agencies release captive green peafowls in order to maintain the wild population of green peafowls. Therefore, in order to better protect the endangered green peafowl, we suggest that purebred identification must be carried out before releasing green peafowls from zoos into the wild in order to preventing the hybrid green peafowl from contaminating the wild green peafowl. In addition, we also found that there were historical introgression events of green peafowl to blue peafowl in four Zoo blue peafowl individuals. The introgressed genomic regions contain IGFBP1 and IGFBP2 genes that could affect blue peafowl body size. Finally, we identified that the nonsense mutation (g.4:12583552G>A) in the EDNRB2 gene is the genetic causative mutation for white feather color of blue peafowl (also called white peafowl), which prevents melanocytes from being transported into feathers, such that melanin cannot be deposited.

## 1. Introduction

The peafowls (Aves, Galliformes, Phasianidae, *Pavo*), which includes the blue peafowl, *Pavo cristatus*, and green peafowl, *Pavo muticus*, are widely regarded as symbols of beauty, nobility, auspiciousness, and good luck in Asian culture^1–3^. They are extensively used in art, religion, literature, and decoration^3–5^. Their irreplaceable symbolic significance has led to extensive research in the fields of ecology^6^, archaeology, history^7^ and genomics^1,8–10^. Some studies have found the phylogenetic positions of the peafowl in Phasianidae, but the phylogenetic relationship of the peafowl to the chicken and turkey in the Phasianidae family is strongly controversial^1,9–11^.

The blue peafowl is distributed in several South Asian countries and widely bred all over the world. It was designated as National Bird of India in 1963 and granted the utmost protection^3,12^. The green peafowl is larger in size than the blue peafowl^4,13^. Unfortunately, as the only native peafowl in China, the green peafowl is classified as Endangered on the International Union for Conservation of Nature (IUCN) Red List and categorized as Critically Endangered on China’s Biodiversity Red List^13–15^. It was once widely and commonly distributed over subtropical and tropical forests in East and Southeast Asia^16,17^. However, the green peafowl suffered a sharp population decline over the past three decades and now the number of wild green peafowls is less than 500 in scattered habitats^13,17^. The endangered situation of the green peafowl is mainly attributed to habitat fragmentation due to climate change and agricultural activate, illegal poaching, etc^14–16,18,19^. However, a critical issue has been overlooked. The blue peafowl and green peafowl are closely related species, and although their wild habitats do not overlap, hybridization between them may still occur due to human breeding or other unknown factors^11,20,21^. The hybrid green peafowls are morphologically difficult to distinguish from purebred green peafowls. Hybridization with closely related species mixes gene pools and potentially losses genotypically distinct populations. These phenomena can be especially problematic for endangered species contacting more abundant ones and could contribute to the extinction of endangered species^22–24^, which has been demonstrated in red wolf, bison Bison bison and endangered Java warty pig^25–28^.

Introgression is the transfer of genetic material from one species into the gene pool of the other by the repeated backcrossing of an interspecific hybrid with one of its parent species^29^. Introgression is a long-term process, and it plays an extremely important role in species traits and environmental adaptation^30^. A typical finding of introgression is the historical introgression of Homo sapiens by Denisovans and Neanderthals to improve the immunity of Homo sapiens^31^. At present, also due to the lack of genomic data and related studies, we do not know whether there is introgression between blue peafowl and green peafowl and whether introgression has an effect on peafowl traits.

White peafowls are blue peafowl plumage colour mutants, which is characterized by the plumages are white but the eyes are black because they contain melanin^32^. Studies have confirmed that the white plumage trait of blue peafowls conforms to the Mendelian law of autosomal segregation^33^. In chickens and geese, this white plumage mutant individual, caused by leucism rather than albinism, is explained by mutations of the EDNRB2 gene^34,35^.

In this study, we firstly constructed the first chromosome-level genome of a blue peafowl including autosomes, sex chromosomes and mitochondrial DNA by using PacBio HiFi CCS and Hi-C sequencing. These genomic resources were utilized to analyze the position of the peafowl in the time-calibrated phylogenetic tree and determine the divergence time between blue peafowl and green peafowl. We also collected and reported datasets of genomic variation in wild and zoo green peafowls and blue peafowls. We aimed to systematically delineate the structure of Asian peafowl genetic diversity, and to detect recent hybridization and historical introgression events between blue peafowl and green peafowl. In addition, we investigated the genetic basis of the white plumage phenotype in peafowls and seeked to ascertain whether the same gene determining white feather color in peafowls are the same as those in other birds, and what are the similarities and differences in the molecular mechanisms of the same phenotype. Our research not only provide evidences for understanding the evolution of peafowls but new insights into saving the endangered green peafowl.

## 2. Methods

### 2.1 Sample collection

In this study, we collected one female green Peafowl (*P.cristatus*) from China Beijing Zoo for HiFi and Hi-C sequencing. We also collected two green Peafowls (*P.muticu*s) from China Beijing Zoo, 12 blue Peafowls (*P.cristatus*) from Hebei XingRui CO.LTD and the China Beijing Zoo, and 6 white Peafowls (*P.cristatus*) from Beijing Agricultural Vocational College for whole genome resequencing, while all resequenced peafowl individuals are artificially bred individuals. In addition, we collected samples of 54 green Peafowls (*P.muticus*) and 3 blue Peafowls (*P.cristatus*) from NCBI databases and CNSA databases (https://db.cngb.org/cnsa/). Among all the peafowl samples from the database, 13 green Peafowls (*P.muticus*) were artificially bred individuals, 44 green Peafowls (*P.muticus*) were wild individuals, and all blue Peafowls (*P.cristatus*) were artificially bred individuals. We obtained the feather follicle tissues of 6 blue peafowls and 6 white peafowls from the peafowl breeding garden in Mentougou District, Beijing for transcriptome sequencing, and combined them with the transcriptome data of 20 blue peafowls published on NCBI for subsequent genome annotation and Transcriptome analysis. In addition, in order to conduct population verification of genetic mutation of plumage color, we obtained blood samples from 11 white peafowls, 29 blue peafowls and 1 pied peafowl (blue white-flight plumage color) from Beijing Agricultural Vocational College and the Peafowl Breeding Park in Mentougou District, Beijing. The full sample information can be found in the Supplementary Table S1. All of the animals in this study were reviewed and approved by Ministry of Agriculture of China (Beijing, China), Animal Welfare Committee of China Agricultural University (Beijing, China).

### 2.2 Blue Peafowl HiFi sequencing and Hi-C sequencing

The genomic DNA extractions were performed on blood from a single female blue Peafowl (*P.cristatus*) individual, using the DNAeasy Blood & Tissue Kit (QIAGEN, Valencia, CA) following the manufacturer’s instructions. The DNA was quantified using the NanoDrop ND-2000 Spectrophotometer (Thermo Fisher Scientific, Waltham, MA) with its standard protocol. The extracted DNA was used to construct PacBio SMRTbell TM library prepared with the Sequel Sequencing Kit 3.0, according to the released protocol from the PacBio Company. The library was processed for PacBio HiFi CCS sequencing (https://github.com/PacificBiosciences/ccs) on the PacBio SequelⅡmachine by Biomarker Technologies company (Beijing, China). A total of 4,085,715,549 bp CCS data was generated. Average CCS length was over 14,503 bp and the longest CCS read length achieved 44,082 bp.

For Hi-C sequencing, the blood was fixed using formaldehyde for 15 min at a concentration of 1%. The chromatin was cross-linked and digested using the restriction enzyme HindIII, then blunt end-repaired, and tagged with biotin. The DNA was religated with the T4 DNA ligation enzyme. After ligation, formaldehyde crosslinks were reversed and the DNA purified from proteins. Biotin-containing DNA fragments were captured and used for the construction of the Hi-C library. The Hi-C library were sequenced on an Illumina HiSeq X Ten platform, producing ∼101.15 Gb Clean Data.

### 2.3 Genome assembly and Hi-C scaffolding

Jellyfish^36^ (v2.3.0) was used to obtain a frequency distribution of k-mer counting with the clean reads, producing kmer frequency distributions of 31-mers. Then, GenomeScope 2.0^37^ was used to evaluate the peafowl genome size. We used hifiasm to assemble the long reads into contigs by using default parameters^38^. Reads shorter than 8000 bp were discarded. We used the juicer (1.6) pipeline to map the Hi-C raw reads against the contigs^39^. Then, the yahs pipeline was used to join the contigs into chromosomes^40,41^. The Hi-C contact map based on the draft chromosomal assembly was then visualized in Juicebox which also allowed for manual adjustment of the orientations and order of contigs along the chromosomes (Supplementary Fig. S1a). Although the newly released genome of the green peafowl is relatively complete, it is poorly collinear with the chicken genome due to imperfect Hi-C scaffolding^8^. We downloaded the Hi-C sequencing data of the blue peafowl and used yash to improve the chromosome-level genome of green peafowl. The consistency and integrity of the assembled blue peafowl genome (WP-1) were separately assessed using BUSCO, based on single-copy orthologues from the AVES (odb10) database. And the Merqury^42^ (v.1.3) was also used to evaluate assembly. To obtain the white Peafowl (*P.cristatus*) mitochondrial genome, we assembled de novo using the NOVOPlasty^43^ (v.4.3.1) with default parameters. The mitochondrial sequence length is 16,694bp (Supplementary Fig. S6).

### 2.4 Blue peafowl genome annotation

To annotate the repeat content, we first used RepeatModeler2^44^ to predict and classify TEs throughout the genome. Tandem repeats were predicted with Tandem Repeats Finder^45^ and the raw results were filtered by the pyTanFinder pipeline^46^. The newly predicted families of TEs and tandem repeats were then combined with the Repbase library (http://www.repeatmasker.org/) (RepBase17.01) to annotate repeats using RepeatMasker (v4.0.7) (http://repeatmasker.org). In addition, we used LTR_finder^47^ to identify long terminal repeat (LTR) sequences.

To predict mRNA-encoding genes in the blue peafowl genome, we performed Ab initio gene prediction, transcriptome based gene prediction and homology-based predictions. For the homology-based predictions, we used protein data from six species (*Homo sapiens*, *Mus musculus*, *Pavo muticus*, *Meleagris gallopavo*, *Gallus gallus*, *Oxyura jamaicensis*) that were retrieved from the Ensembl (release 64) database and identified candidate coding regions with GenomeThreader^48^. For the transcriptome based gene prediction, We mapped the collected RNA-seq reads using HISAT2 (v.2.1.0)^49^, and assembled the transcriptomes using StringTie^50^.

TransDecoder (https://github.com/TransDecoder/TransDecoder) was used for identifying candidate coding regions within transcript sequences. To make de novo predictions, BRAKER2 (v.2.1.6)^51^ was run to use the homology protein as hints to generate predicted gene models from AUGUSTUS (v.3.4.0)^52^ and to train the hidden Markov model (HMM) of GeneMark-ET (v.3.67_lic)^53^. EVidenceModeler software^54^ was used to integrate the gene set and generate a non-redundant and more complete gene set via integration of the three respective annotation files that were assigned different weights (ab initio prediction was “1”, homology-based prediction was “5”, and transcriptome-based prediction was “10”). Finally, PASA^55^ was used to correct the annotation results of EVidenceModeler for the final gene set. Functional annotation of the protein-coding genes was accomplished using eggNOG-Mapper (v.2)^56^, a tool that enables rapid functional annotations of novel sequences on the basis of pre-computed orthology assignments, against the EggNOG (v.5.0) database^57^. And the protein database of SwissProt^58^, NR^59^, Pfam^60^ were also used to annotate Gene function. We also mapped the reference genes to the KEGG pathway database^61^ and identified the best match for each gene. tRNAscan-SE (v.2.0.11) software^62^ was used to search for the transfer RNA (tRNA) sequence of genome, with INFERNAL^63^ (v.1.1.3) from Rfam^64^ to predict microRNAs and snRNA of genome. And barrnap (v.0.9) (https://github.com/tseemann/barrnap) was used to annotated ribosomal RNA (rRNA) sequence of genome (Supplementary Fig. S1b).

For mitochondrial genome annotation, we submitted it to the AGORA web tool^65^, with the protein-coding and rRNA genes of the Gallus gallus mitochondrial genome (accession number: NC_053523.1) as a reference.

### 2.5 Genome synteny and collinearity analysis

We used the MUMmer^66^ (v.4.0.0rc1) tool nucmer to perform the pairwise whole genome alignment with the parameter “-b 400”. The alignments were filtered to keep the oneto-one best hits using delta-filt from the MUMmer package. Unanchored scaffolds were excluded from the alignments. We also performed collinearity analysis using MCscanx^67^ with default parameters. The NGenomeSyn^68^ (v.1.4.1) was used for synteny visualization.

### 2.6 Phylogenetic tree and divergence time

The protein sequences of 15 species were used to search the orthologues using OrthoFinder2^69^. The results showed that a total of 17,999 orthogroups were identified in 15 species, of which 2,323 single-copy orthologues were shared among these species. These single-copy orthologues were subsequently converted into coding sequence alignment by tracing the coding relationship using pal2nal.v14^70^. Then, we construct a phylogenetic tree using RAxML^71^ (v.8.0.0) with commands: “raxmlHPCAVX -N 100 -b 15738 -f j -m PROTGAMMALGX -s sample.phy-n all.tree”. Each of the 100 bootstrapped alignments was used to generate a guide tree with “GAMMA” model by RAxML and converted into a binary file using “parse-examl” command. Each pair of the guide tree and the binary file generated from 100 bootstrapped alignments were inputted for ML analysis with ExaML^72^ (v.3.0.15) and generated trees with branch lengths. Parallel method is also used to construct phylogenetic tree with ASTRAL^73^. Additionally, the divergence time of all species was estimated and calibrated through the divergence time between *Lagopus muta* and *Lagopus leucura*, *Tympanuchus pallidicinctus* and *Centrocercus urophasianus*, *Bambusicola Thoracicus* and *Gallus Gallus*, *Cygnus Atratus* and *Oxyura Jamaicensis*, *Columba Livia* and *Amazona Guildingii*, *Gallus Gallus* and *Bambusicola Thoracicus*, *Tympanuchus pallidicinctus* and *Centrocercus urophasianus*, *Lagopus muta* and *Lagopus leucura*, and *Cygnus Atratus* and *Oxyura Jamaicensis* from the TimeTree database^74^. We then used BASEML to estimate the overall mutation rate with the time calibration on the root node (94.39 MY). General reversible substitution model and discrete gamma rates were estimated by the maximum likelihood approach under strict clock. The divergence time was then estimated using MCMCtree^75^. In addition, we also collected the mitochondrial sequences of 11 species from the NCBI, combined with our assembled mitochondrial sequences, constructed a ultrametric tree of maternal inheritance.

We aligned the genomes of chickens, turkeys and two species of peafowls, with swans as the outgroup to form a five-species whole genome alignment data by using cactus (v.2.6) with default parameters^76^. For the phylogeny of local alignments of sliding windows among all compared genomes, we partitioned the data into nonoverlapping sliding windows with varying window sizes of 100kb to reconstruct phylogenetic trees. All windows with more than 75% gaps were removed. RAxML was used to construct maximum likelihood trees from each alignment, and the low consensus trees (bootstrap < 50%) were discarded^71^.

### 2.7 Estimates of the effective population size and divergence time

To characterize the historical demography of the wild green peafowl and blue peafowl, a strategy of combination of two complementary algorithms was employed for cross-validation and comparison. In general, the popular individual-genome-based PSMC approach was used to retrieve distant past history, while the SMC++ method based on various algorithms and data exploration were employed to better characterize more recent demography. For all of these analyses, the mutation rate was set to 1.33 * 10^-9^ per site per year estimated for the Indian peafowl^77^ and the generation time assumed to be four years to scale demographic events in calendar times, following a study on the blue peafowl^78^. The PSMC^79^ analysis, which utilized LD information, was conducted on autosomes of the high-coverage de novo assembly. We used bcftools to generate autosomal fasta files, respectively, according to recommendation in the PSMC documentation. The PSMC 100 bootstraps were performed with parameters optimized for birds^80^ (N30 –t5 –r5 –p 4+30*2+4+6+10) to determine variance in *Ne* estimates. And we plotted the PSMC by using python script (https://github.com/shengxinzhuan/easy_psmc_plot). A recently published approach with higher resolution in the recent past compared to PSMC accuracy, SMC++^81^, was used to predict the demographic history (or population sizes and divergence times) of green peafowl and blue peafowl based on multiple unphased individuals. The short generation time of wild green peafowl and blue peafowl makes possible the reliable and precise estimation of effective population sizes in the recent past using the method of SMC++^82^ (v.1.15.2).

We also used DADI^83,84^ to infer the divergence time between wild green peafowl and blue peafowl. We simulated four models with the same dataset under the two-population model in DADI independently. Model 1 (sym_mig): Instantanous size change followed by exponential growth with no population split; Model 2 (bottlegrowth_2d): Instantanous size change followed by exponential growth then split with migration; Model 3 (bottlegrowth_split_mig): Split into two populations of specifed size, with migration; Model 4 (split_asym_mig): Split into two populations of specifed size, with asymetric migration. To avoid the effect of selection on the demographic analysis, we focused on non-coding regions and extracted non-coding SNPs. To avoid the linkage of SNPs, we thinned the non-coding SNPs to 1% and obtained a dataset of 135,873 SNPs^85^. Owing to the unknown ancestral state of each SNP, we folded the frequency spectrum. The projection value was selected based on the strategy of maximizing the number of segregated sites. The starting parameter values for the first round were randomly assigned, and the best parameters obtained after completion of a round were used as the starting parameters for the subsequent rounds. After the convergence of the parameters, we retained the model with the highest log-likelihood as the final simulation result. The parameters of the optimal model were converted into absolute units using the average mutation rate per generation and generation interval. Confidence intervals for the parameters were generated using the Godambe information matrix (GIM) with 100 bootstraps^86^.

All blue peafowls and green peafowls populations used for SMC++ and DADI analyzes excluded hybrid and introgressed individuals found in subsequent studies.

### 2.8 Gene family construction and Branch-specific positive selection

To define gene families, we used coding sequences of all 15 species and extracted the longest protein for each gene. Gene family size expansion and contraction analysis was performed by CAFE5^87^ (v.5.0.0), and the results from OrthoFinder2 and a phylogenetic tree with divergence times were used as inputs for CAFE. These expanded gene families were annotated and classified through the analysis of GO ontology and KEGG pathways to further explore the impact of adaptive evolution on peafowl by using Kaobas ^88^ (v.3.0).

We obtained codon alignments for the single-copy orthologous groups by aligning Coding sequence by using MAFFT. Nonsynonymous and synonymous substitution rates (Ka/Ks) were calculated using codeml program in PAML (v.4.5) package^75^. We used the branch site model and two-ratio models to detect signatures of natural selection on coding genes of Neofelis. Statistical significance was determined using likelihood ratio tests. Functional annotation of the obtained gene dataset was also performed using Kobas^88^ (v.3.0).

### 2.9 Whole-genome resequencing and variant calling

We cleaned the Illumina NGS raw data to remove adaptors, trim low-quality bases and remove “N” sites with fastp^89^ (v.0.20.0). Subsequently, clean reads were mapped to our blue peafowl genome using BWA^90^ (v.0.7.10-r789) with default parameters. High-quality mapped reads (mapped, nonduplicated reads with mapping quality ≥ 20) were selected with SAMTools^91^ (v.1.3.1) and the following commands: “-view-F 4 -q 20” and “rmdup”73. For all samples, we used the “bamqc” module in Qualimap^92^ (v.2.2.1) to perform sequencing depth statistics. Only high-quality mapped reads were used for variant calling with GATK^93^ (v.4.2.6.1). BAM files were sorted and marked as PCR duplications with Picard (v.2.27.5, http://broadinstitute.github.io/picard/). There is no well-annotated SNP and short-indel database for blue peafowl, so it was not feasible to use the “Base Quality Score Recalibrator” (BQSR) and “IndelRealigner” options of GATK. To carry out variant calling, we used the command “HaplotypeCaller”, which calls SNPs and indels simultaneously via local de novo assembly of haplotypes in an active region. Applying the “hard filtering” method, we obtained an initial set of high-confidence SNPs and indels. The parameters of “hard filtering” were set by default, i.e., for SNPs we used QD < 2.0, FS > 200.0, SOR > 10.0, MQRankSum < −12.5, and ReadPosRankSum < −8.0, while for short indels, we considered QD < 2.0, FS > 200.0, and ReadPosRankSum < −8.0. After the initial filtering step, the number of SNPs and short Indels became 16,330,028 and 2,153,530, respectively. Notably, the ratios of Ts/Tv to 2.534 with filtering of the raw SNPs, which showcases the high quality of the SNP call sets. For many downstream analyses, the core set of SNPs and Indels were acquired by setting the MAF cut-off at 0.05. We performed SNP and Indel annotation according to the blue peafowl genome using the SnpEff^94^ (v.5.1). The intergenic region of the genome encompasses about 45.73% of SNPs and 70.0% of Indels. About 1.573% of SNPs are located in the coding sequence, and the nonsynonymous to synonymous SNP ratio is 0.508%. In comparison, only 1.081% of Indels are found in the coding sequence. In addition, we also used CNVcaller to detect copy number variations in *P.cristatus* for subsequent analysis of the feather color of white peafowl^95^.

### 2.10 Population genetic structure

We chose the core set of SNPs (MAF greater than 0.05) for additional pruning. PLINK^96^ (v1.90b6.12) was used to remove SNPs having high LD (r^2^≥0.5) within a continuous window of 50 SNPs (step size 5 SNPs), which yielded 3,147,759 SNPs for both analyses. The parameter used in this procedure was “--indep-pair-wise 50 5 0.2”. The results obtained from the above procedure was be used to perform principal component (PCA) analysis by using VCF2PCACluster (https://github.com/hewm2008/MingPCACluster). We used ADMIXTURE software^97^ to analyze the population structure of all peafowl samples with kinship (K) set from 2 to 5. Finally, we constructed the unrooted Neighbor-joining (NJ) tree by MEGA software^98^ (v.11). The FigTree software (v.1.4.4) was used for visualization. We also constructed the maximum likelihood (ML) tree for all individuals using treemix^99^ (v.1.13).

### 2.11 Population genetic diversity

As for calculation of the nucleotide diversity and the linkage disequilibrium decay of each peafowl population, we used two softwares, VCFtools^100^ (v.0.1.16) and PopLDdecay^101^ (v.3.42) with default parameters, respectively. In the study of nucleotide diversity and ROH, we used the same set of SNPs. Since LD analysis and nucleotide diversity analysis require the number of individuals in the peafowl population to be greater than 1, we removed populations with 1 individual and hybrid individual. Runs of homozygosity (ROH) of each individuals peafowl were identified using the homozyg option implemented in the PLINK, which slides a window of 50 SNPs (-homozyg-window-snp 50) across the genome estimating homozygosity. The following settings were performed for ROH identification: (1) required minimum density (-homozyg-density 50); (2) number of heterozygotes allowed in a window (-homozyg-window-het 3); (3) the number of missing calls allowed in window (-homozyg-window-missing 5). The ROH of each peafowl breed were counted and divided into four categories according to the length: 0.5-1Mb, 1-2Mb, 2-4Mb, >4Mb^102^.

### 2.12 Whole mitochondrial genome phylogeny

The BAM alignments were converted to fastq and subsequently used with Mapping Iterative Assembler (v.1.0) (https://github.com/mpieva/mapping-iterative-assembler) to assemble a mtDNA consensus sequence. We aligned our 76 mitochondrial genome sequences to a collection of 1 published *Gallus gallus* mitochondrial genome (accession number: NC_053523.1). The “TIM3+F+G4” model of nucleotide substitution was selected by comparing the Bayesian information criterion (BIC) scores in jModelTest^103^ (v.2.1.10). A phylogenetic tree was then inferred using ML methods. The ML analysis was performed with the program Raxml^71^ (v.8.2.12), and approximate likelihood-ratio tests were performed to establish statistical support of internal branches with the Gallus gallus as outgroup. The mitochondrial haplotypes were built by using DnaSP^104^ (v.6.12.03). And the PopART^105^ (v.1.4.4) was used for network haplotype visualization.

### 2.13 Hybridization analysis between green peafowls and blue peafowls

We used a chromosome painting approach with ancestry-informative sites to validate the delimitation of ancestry blocks detected by the HMM and to visualize patterns of introgression across the blue peafowl. This approach provides a lower level of resolution for ancestry block delimitation but with higher power to classify regions as derived from either parental genome. To identify introgressed intervals in Hybrid Individual of Pavo muticus, we used Loter (v.1.0.1)^106^ to infer the haplotype fragments of blue peafowl among all autosomes of Pavo muticus genomes. In addition, we identified alleles that were differentially fixed in green peafowls and blue peafowls parental populations and had no missing data using the script get_fixed_site_gts.rb. We thinned SNPs to be a minimum of 1 kb apart and mapped these ancestry-informative sites in the green peafowl samples using the scriptplot_fixed_site_gts.rb (https://github.com/mmatschiner/tutorials/blob/master/analysis_of_introgression_with_snp_data).

### 2.14 Introgression analysis

Treemix^107^ (v1.13) was used to confirm relationships between our focal populations and to visualize migration events between populations. We first built the maximum likelihood tree (zero migration events) in Treemix and then ran Treemix sequentially with one through ten migration events. We supplied this set of 5,671,362 biallelic SNPs to Treemix, rooted with Gallus gallus, and estimated the covariance matrix between populations using blocks of 500 SNPs. Three samples (qhd-01, qhd_02 and GF-f1A) were excluded from this analysis because ADMIXTURE indicated that they were likely hybrid individual. We calculated the variance explained by each model (zero through ten migration events) using the R script treemixVarianceExplained.R.

To investigate the relationship of the blue peafowl to green peafowl populations, we used AdmixTools^108^ (v.7.0.2) for the phylogenetic analysis. We also used three-population test estimates (f3 statistics) to test for admixtures across all peafowl populations. Three-population tests consider population triplets (C; A, B), where C is the test population and A and B are the reference populations. Significantly negative Z-scores (Z ≤ −3.80, after Bonferroni correction for multiple testing) indicate evidence of test population C containing an admixture of both reference populations of A and B.

The ABBA-BABA test, also known as the D-statistic, was used to infer the existence of gene flow between the populations. This analysis of D-statistic values was performed using Dsuite^109^ software, which can calculate D-statistics at the genome scale across all combinations of populations with VCF input files. According to the principle of ABBA-BABA test calculation, (P1, P2, P3, O) represent four different groups. And O is chicken (*Gallus gallus*) as the outgroup. Gene flow between domestic geese and their wild ancestors was inferred with the mallard as an outgroup. Dtrios was used to calculate the D and f4-ratio statistics for all trios of populations in the dataset, with a default value of 20,000 for the Jackknife block size. Then, using the Dinvestigate program to calculate the D-value for windows containing useable-size SNPs, the sliding window consisted of 2500 SNPs and a step of 500 SNPs. The locations of the windows with the top 5% D values were obtained and the genes in these windows were regarded as candidate genes for introgression. Functional annotation of the obtained gene dataset was performed using Kobas^88^ (v.3.0).

### 2.15 Histological characterization of the genetic mechanism of plumage color in white peafowls and elucidation of the molecular basis

In order to understand the reason of white peafowl (*P.cristatus*) plumages turning white, we used the peafowl individual resequencing SNP data set and used VCFtools to calculate *Fst*^100^. We calculated the genome-wide distribution of *Fst* between Blue peafowl (*P.cristatus*) and White peafowl (*P.cristatus*), using a sliding-window approach (1kb windows with 200bp increments). Absolute genetic divergence (Dxy) statistic was calculated to strengthen our results by using custom python scripts^110^. The bedtools software (v.2.30.0) was used to annotate the selection regions.

For RNA-seq data analysis, the paired-end reads were mapped to the peafowl reference genome (WP-1) using the HISAT2 (v2.6.1.0) software after quality control^49^. Transcripts were assembled and quantified with the StringTie (v2.1.1) software^50^. The GFFcompare (v0.12.6) was used to compare the alternative transcripts among individuals^111^. The differential expression analysis was performed using the DESeq2 (v1.4.5) package^112^. Ultimately, we enriched candidate genes by the online website KOBAS^88^(v3.0), so as to grasp the functions of selection genes and Differential Expression Genes (DEGs). To identify the causative mutation that causes blue peafowl feathers to turn white, we also examined the frequency of all (SNP, Indel and CNV) mutations within 10 kb before and after the candidate SNP.

DNA was extracted from blood samples with the DNAeasy Blood & Tissue Kit (QIAGEN, Valencia, CA). The EDNRB2 gene SNP mutation (g.4:12583552 G>A) was genotyped by Sanger sequencing on an ABI 3730xl DNA Analyzer (Applied Biosystems, USA) according to the manufacturer’s instructions. The primer sequences used were as follows: forward primer 5′-TGAAGAAGTGTAAGTCCCGCTG-3′ and reverse primer 5′-AGGTCTCGGTCCCAGTAGTT-3′.

Plumage samples stored in 4% paraformaldehyde were washed in phosphate buffered saline (PBS), dehydrated with a gradient of alcohol solutions (50% → 70% → 80% → 95% → 100% → 100%), cleared with xylene and infiltrated with paraffin wax. Samples were embedded in paraffin and sectioned into 4-micrometer-thick tissue sections. Both the plumage samples sections were stained with hematoxylin and eosin (H&E) and Masson-Fontana. We designed a fluorescent in situ hybridization (Fish) probe using the specific sequence of the CDS region of the EDNRB2 gene. Sections were incubated with 4′,6-diamidino-2-phenylindole (DAPI) to stain the nucleus and were imaged with a fluorescence microscope.

## 3. Result

### 3.1 Sequencing, assembly and annotation of the blue peafowl genome

The blue peafowl genome is comprised of 78 chromosomes, including at least 30 microchromosomes (76 autosomes + ZW sex chromosomes) according to the karyotype analysis^113^. In order to assemble a chromosome-level genome for blue peafowl, we generated 38 Gb of PacBio Circular Consensus Sequencing (CCS) HiFi reads and 101.15 Gb of chromatin conformation capture (Hi-C) reads. K-mer-based analyses of the Illumina paired-end sequencing reads (51.77 Gb, 52.71 × sequencing depth) estimated the size of the nuclear genome to be approximately 982.21 Mb (Supplementary Fig. S2 and Table S1).

The initial assembly was 1.13 Gb, consisting of 389 contigs with a N50 length of 30.6 Mb, indicating a high contiguity of the assembly. Contigs were then concatenated to the chromosome-level assembly by Hi-C reads. We ultimately obtained 36 pairs of autosomes (9 macrochromosomes, 27 microchromosomes) and ZW sex chromosomes with genome size (1.041Gb) (Supplementary Fig. S3 and Table S2). Our genome exhibits a 500-fold and 8-fold improvement, in the scaffold N50 (95 Mb), compared to those of the previously published peafowl genomes reported by Dhar et al.^9^ and Liu et al.^1^, respectively.

We next evaluated the quality of the genome assembly using Benchmarking Universal Single-Copy Orthologs (BUSCO)^114^, Merqury^42^ and Illumina short reads. The complete BUSCO of the blue peafowl genome assembly was 97.1%, indicating a high completeness of the gene space (Supplementary Fig. S4 and Table S3). Merqury compares k-mers from the assembly to those found in unassembled HiFi reads to estimate the completeness and accuracy. The completeness and quality value (QV) of the genome were 96.34% and 60.47 (>99.99% accuracy) respectively. These results attest to the high accuracy and completeness of our assembly (Table 1). According to our results, repetitive sequences accounted for 13.38% of the blue peafowl genome (Supplementary Table S4), including 10.44% tandem repeats and 7.35% transposable element proteins. Among tandem repeats, Long Interspersed Nuclear Elements (LINEs) constitute the majority, accounting for 7.41%. Additionally, short interspersed nuclear elements (SINEs) 0.05% and long terminal repeats (LTRs) accounts for 2.99%. We also identified 354 microRNAs, 308 tRNAs, 151 ribosomal RNAs, and 334 small nuclear RNAs (Supplementary Table S2 and S5). For annotation, the collected mRNA sequencing (RNA-seq) data were aligned to the reference genome, and 37,401 putative protein-coding gene models were predicted.

**Table 1.**
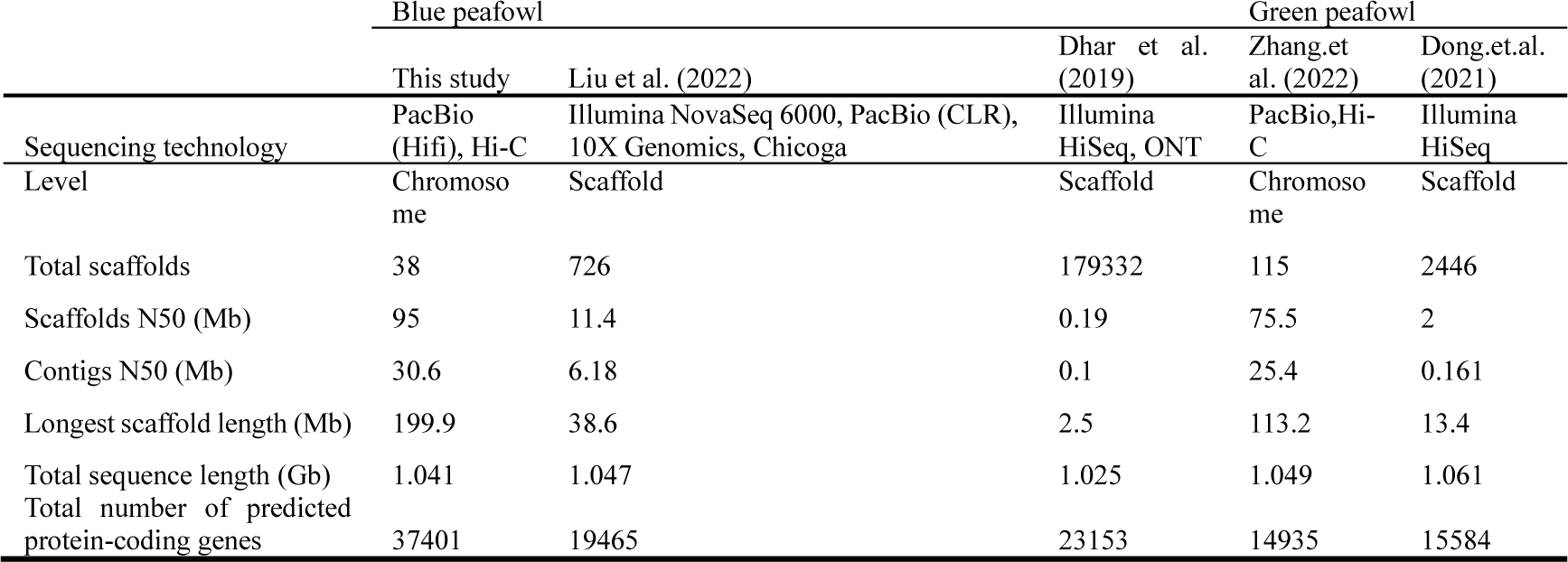
Quality metrics for the blue peafowl genome assembly generated in the current work and for other blue peafowl and green peafowl genome assemblies published in previous studies.

### 3.2 Peafowl genome rearrangement

Synteny and genome size have generally remained stable over the more than 100 million years’ of modern bird evolution^115–118^. Among the 12% of bird species with documented karyotypes, most have diploid numbers ranging from 76 to 82^115–117^. Saski et.al^113^ found that peafowl has a typical bird karyotype with 2n=78 chromosomes, which is consistent with that of chicken. In this study, we assembled 10 pairs of large chromosomes of 2n=74 in total, which slightly conflicts with the results of the karyotype analysis showing that peafowl have 8 large chromosomes and 30 pairs of microchromosomes of 78 in total (Fig. 1A). Newly improved green peafowl genome has the same number of chromosomes as the blue peafowl.

**Fig. 1.**
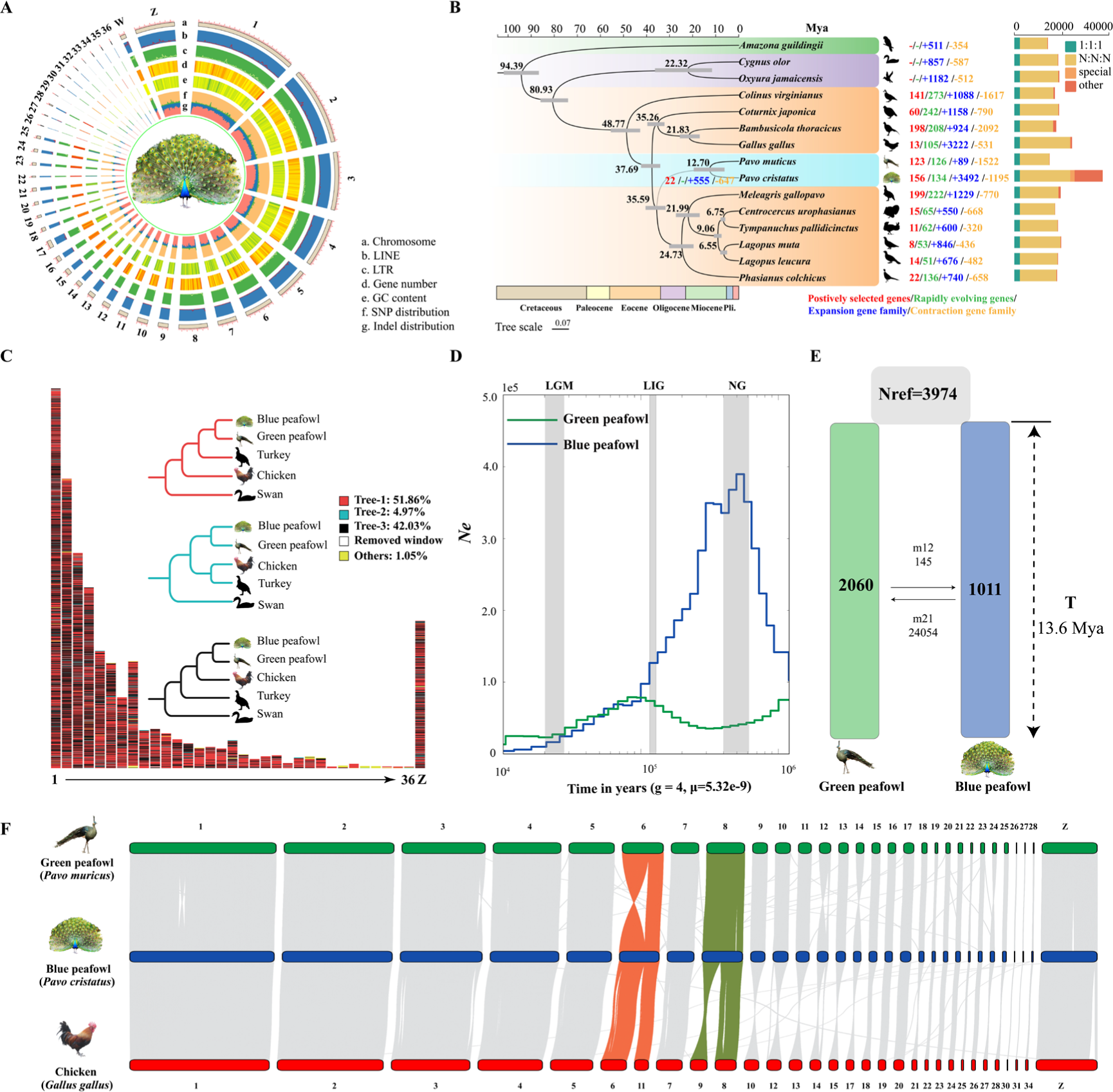
Assembly, composition, evolution and demographic history of the blue peafowl genome. (A) Blue peafowl genomic features. (B) Positively selected genes (PGS), rapidly evolving genes (RGs), Gene family, phylogenetic and molecular clock dating analysis of the blue peafowl genome with 15 other species, based on single-copy orthogroup data. The grey bars at the internodes represent 95% confidence intervals for divergence times. In the histogram, “1:1:1” indicates that the single-copy orthologs are shared by 15 species with 1 copy. “N:N:N” represents any other orthologous group (missing in 1 species). Species-specific shows the specific orthologs in each species. Other orthologs are unclustered into gene families. (C) Distribution of the genomic phylogenetic discordance across each chromosome of blue peafowl by 100-kb sliding windows. The different colors represent the variable phylogenetic topologies. (D) Population size history inference of blue peafowl (BP) and green peafowl (GP) respectively, acrossing the last glacial maximum (LGM, approx. 26–19 Ka), the last interglacial period (LIG, approx. 132–112 Ka) and the naynayxungla glaciation (NG, approx. 0.5–0.72 Ma). (E) Best-fit estimates of parameters by using DaDi showed the divergence time of blue peafowl and green peafowl. (F) Genome synteny and collinearity among the blue peafowl and chicken.

To understand the similarities between the common peafowl and chicken at the chromosome level, we compared our assembly with the chicken reference genome (*Gallus gallus* 7.0). Most of the large scaffolds had their counterparts to the macrochromosomes in chicken except scaffolds 6 and 8 (Fig. 1F). Among them, chicken chromosomes 6 and 11 were both aligned to blue peafowl scaffolds 6, whereas chromosomes 8 and 9 added up to scaffolds 8. We also compared the assembly with the turkey reference genome (Turkey 5.0). Correspondingly, the first nine scaffolds have their unique counterparts in the macrochromosomes of turkey (Supplementary Fig. S5).

Despite the overall high consistency of the common blue peafowl, chicken, and turkey genomes, some chromosomal rearrangements were observed (Fig. 1C). Because of the GC-richness, specific repeats, high microchromosome mutation rate in birds^119^, and lack of cytogenetic supports, determining whether these short scaffolds are bona fide chromosomal rearrangements or simply assembly errors is difficult, and further studies are required in this regard^120^.

### 3.3 Phylogenetic Trees Resolving the Divergence Time of blue peafowl and green peafowl

Peafowl is a general name for a type of bird of the order Galliformes and Phasianida, which is a consensus in past studies on both morphology and molecular biology^10^. However, local genomic phylogeny within extant Phasianidae, the phylogenetic relationship of peafowls with turkeys and chickens has always been controversial ^78^.

We obtained 17,999 gene families and 2,323 single-copy orthologous genes from blue peafowl, green peafowl and other 13 species. We generated a whole-genome species tree with the Psittaciformes and Anseriformes as outgroups by using 1,271,402 sites from the single-copy orthologous genes sets of 15 species, yielding a species tree with 100% bootstrap support for all nodes (Fig. 1B).

According to our results, the species tree clearly split into Psittaciformes, Galliformes and Anseriformes. Galliformes order is clustered, within which the Phasianidae family formed a group. Blue peafowls and green peafowls cluster together in the Phasianidae branch. To ensure that the phylogeny was robust, we used the coalescent-based phylogenetic method and the same gene set. As expected, the topology of the astral coalescent tree was identical to the species tree (Supplementary Fig. S7-S8).

Using the single-copy orthologous genes evidence tree and fossil calibration data, the divergence dating of the peafowl was estimated by means of the MCMCtree algorithm. We estimated that the most recent common ancestor of peafowl evolved between 31.71 and 40.47 million years ago (Mya). The blue peafowl and green peafowl diverged 12.69 Mya within the range of divergence (6.15–19.5 Mya). Their divergence coincided with the Miocene epoch, which entered a series of ice ages. At this point, new species replace the older types, rapid evolution of more species. Several species of Phasianidae, including *Meleagris gallopavo, Centrocercus urophasianus, Tympanuchus pallidicinctus* and *Lagopus*, evolved during this period. Notably, peafowl were found to be closer to turkey than chicken in the Phasianidae family. These findings were consistent with the findings of those reported by Liu et al.^32^. However, when we used mitochondrial sequences to construct phylogenetic tree, peafowls were shown to be more closely related to chickens than to turkeys (Supplementary Fig. S9). This opposite result attracts our attention.

To resolve these controversies, we constructed the topologies of locally aligned fragments across the whole genomes of two peafowls and chickens and turkeys with swan as outgroup. Our results found more than 40% phylogenetic discordance, which was evenly dispersed along the whole reference genome architecture (Fig. 1C).

As is well known, birds survived the Cretaceous–Paleogene extinction event (K-Pg) and underwent multiple species explosions ^121^. These species explosion events resulting in incomplete lineage sorting (ILS) have led to a much greater degree of confusion between branches of bird species compared to mammals.^122^. The speciation of peafowls coincided with the period of the pheasant family’s major explosion. Therefore, we speculate that the high proportion of confusion in the genome-wide phylogenetic trees of peafowls, chickens and turkey in this study is also caused by ILS.

### 3.4 Peafowls have different paternal and maternal demographic histories

To understand the demographic history in peafowls, we investigated each species with the pairwise sequentially Markovian coalescent (PSMC) approach and SMC++. PSMC model to infer the local time to the most recent common ancestor (TMRCA) as well as to assess changes in the historic effective population size (Ne) by taking into account whole-genome data from single deep-coverage (>25×) individuals. The SMC++ method was used to reconstruct the population history of the “core” groups of green and blue peafowls found in subsequent studies with a generation time of g = 4 and a mutation rate per generation of μg = 1.9 × 10-9.

In our results, the Ne changes of wild green peafowl exhibited by the two methods are somewhat different during the LGP period. The result of the wild green peafowl PSMC is consistent with the results of Dong et al, which showed early population decline from 800 to 210 ka, followed by a bounce to a peak Ne during early LGP (ca. 70 Ka), and a more marked decrease (seven-fold change) throughout the following part of LGP (ca.70–10 Ka) (Fig. 1D). SMC++ analyses showed that during the LGP period, the effective population size experienced a process of first decreasing (ca.110-30 ka) and then increasing (ca.10-20 ka), and suggested a peak Ne for the green peafowl after the LGP period (ca.10 ka). This may be due to differences in the individuals used in the analysis. Such a fluctuating population history broadly agrees with the two demographic patterns of most other threatened species, plausibly suggesting a common genetic consequence of late Pleistocene climatic oscillations.

However, blue peafowl showed an entirely different group history. Both PSMC and SMC++ population-based demographic analyses congruently suggested that the Ne of the blue peafowl experienced a dramatic expansion from 500 to 1000Ka and peaked around 500Ka before the LGP period, and then declined (with a small-scale recovery during the period). We speculate that the blue peafowl may not have adapted to the severe environmental changes during the last glacial period compared to the green peafowl.

We then used the simulation of diffusion approximations for demographic inference (DADI) to calculate the divergence times between the “core” groups of the blue peafowl and the green peafowl in this study. Among the four models definded, Model 4 (split_asym_mig) had the highest log-likelihood, indicating the best fit of model to the data (Fig. 1E, Supplementary Fig. S10, S11 and Table S6). The demographic paraments estimated using model 4 indicated that blue peafowl and the green peafowl diverged ∼13,624,718 years ago with a 95% confidence interval (CI) of ± 31180 years (95% CI ± 31180) which overlapped with the molecular divergence times of blue peafowl and the green peafowl by MCMCtree. The Ne values of blue peafowl and the green peafowl were 1011 (95%CI ± 10) and 2060 (95%CI ± 22). Model 4 also indicated that nonsymmetric gene flow existed between blue peafowl and the green peafowl after divergencing.

### 3.5 Genetic mechanisms underlying peafowls body size and immune system evolution

Like other birds groups^122^, extant Phasianidae species exhibit a large range of body sizes, from *Coturnix japonica* (∼14 cm) at one end of the spectrum to peafowls (>2.3 m in some individuals with 1.5 m tail plumages) at the other^8^. Thus, phasianidae body size has experienced significant divergence, particularly for the green peafowls with their substantial enlargement in body size.

We detected 123 positively selected genes (PGS) and 126 rapidly evolving genes in the green peafowls (Fig. 1B). Among these genes, IGFBP4 and IGF2, which are related to insulin-like growth factors, may have contributed to the evolution of this trait. Several studies on other vertebrate species corroborate the critical role of IGF-2 in body growth and adult size determination^123^. IGFBP4 is IGFs binding partner proteins are known to cause changed body size in mouse and human^124^. Blue peafowls are slightly smaller than green peafowls, but it is still among the largest species in the pheasant family^9^. We detected that MSTN has undergone rapid evolution in blue peafowls. MSTN is well-known as muscle growth inhibitory gene. Some cattle breeds (Belgian blue cattle) exhibit extremely exaggerated body size and muscle content due to mutations in this gene^125^. In birds, the rate of sequence divergence in immune-related genes is usually higher than in the other genes primarily because of the co-evolution of host–pathogen interactions^126^. We performed enrichment analysis on 555 expanded gene families in the common ancestors of the peafowls and found that eight genes (IFNW1, ACTG1, NUP98, IFNA3, KPNA2, ACTB, RNASEL, NXF1) were enriched in Influenza A KEGG pathway (gga05164) which related to immunity^127^ (Fig. 1B and Supplementary Fig. S12). In addition, we also found that the interleukin family gene IL7R and the CD8BP gene involved in the adaptive immune response were positively selected in the common ancestor of the peafowls (Supplementary Table. S7). Although rapid evolution of immune genes was found in a wide range of birds^122^, the number of PSGs and RGs genes involved in the Cytokine-cytokine receptor interaction pathway is highest in in blue peafowls (NODA, IL13RA1, BMP8A, GDF5) and green peafowls (CCR10, CD40LG, ACVR2B, CCR7) (Supplementary Fig. S13, Table. S8 and S9). It could suggest a key role of this pathway in the evolution of peafowl immune system. Among these genes, major histocompatibility complex (MHC) genes, interleukin family genes, and NF-KB signaling pathway genes play important roles in immune responses.

### 3.5 Genome resequencing

In this study, 18 blue peafowl and 2 green peafowl different geographical locations in China were selected for genome resequencing (Fig. 2A). For a more comprehensive analysis of peafowls, we also combined our data with available whole-genome resequencing data for 3 blue peafowl individuals and 54 green peafowl individuals from 7 regions in Asia, giving a total of 77 individuals (Supplementary Table S10 and S11). Whole-genome resequencing sequences were mapped to our assembled genome (WP-1). After quality control and filtering, 1399.13 Gb of high-quality sequences were obtained, with an average of 17.48 Gb per individual. Across all samples, a total of 12,453,511,109 mapped reads was obtained, with an average depth of 18.53 × and an average coverage of 92.07% per individual. After variant calling, a total of 27.78 million variants was obtained, including 16.33 million SNPs and 0.47 million indels. Among all variants, only 1.82% of SNPs and 1.08% of Indels are located in exons. Most mutations are located in intergenic regions (Supplementary Fig. S14). In addition, we also collected a rooster as an outgroup for subsequent analysis. Of these animals, 76 peafowl individuals were used for a mitochondrial sequence analysis and 38 female peafowl individuals used for a W-chromosome SNP analysis.

**Fig. 2.**
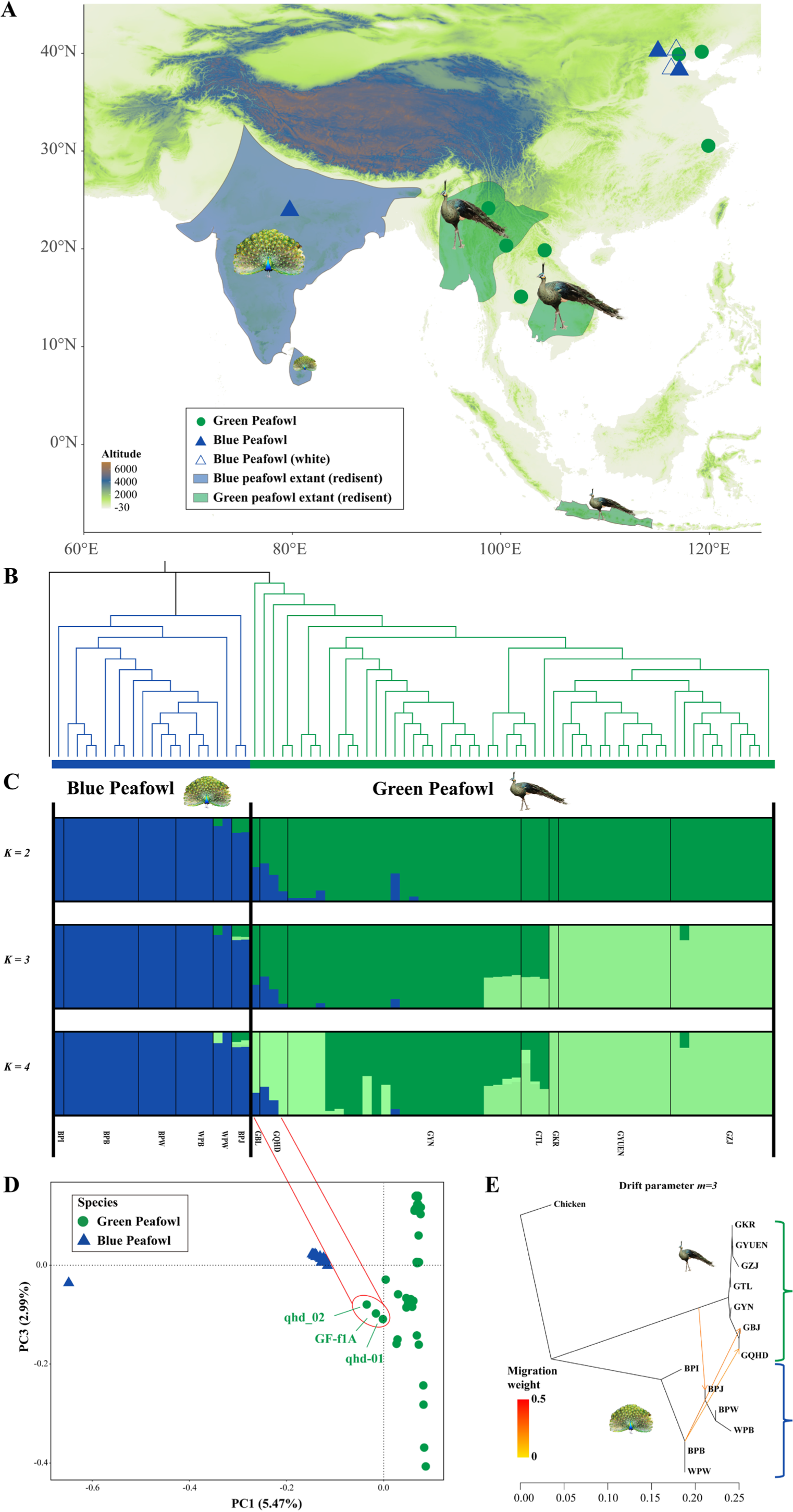
(A) Location of the samples used for this study. A total of 77 peafowls, including blue peafowl (n= 22, *P.cristatus*) and green peafowl (n= 55, *P. muticus*), were used. (B) A NJ tree of all peafowls estimated based on high-quality autosomal SNPs. (C) Population genetic structure of peafowl. The length of each colored segment represents the proportion of the individual genome from the ancestral populations (K=2–4), population names are at the bottom. (D) PCA plot of peafowl individuals. The three individuals in the red circle are the green peafowl that may have hybridized with the blue peafowl. (E) Genetic migration inferred using TreeMix with best draft parameter (m=3), while the migration weight (proportion of admixed population received from source) is indicated on the arrow by the number and color.

### 3.6 Population genetic structure of peafowl using autosomal variants

Genomic SNPs of blue peafowl and green peafowl were used to analyze the population structure of peafowl, as well as to analyze the introgression between blue peafowl and green peafowl.

The NJ tree of SNPs filtered and trimmed for linkage disequilibrium using chickens as an outgroup demonstrated a clear genetic structure with green peafowl and blue peafowl clustered into each clade (Fig. 3B). Admixture analysis (K from 2 to 9) of the genomic admixture with two population scenarios (K=2) had the best likelihood, and the specimens were divided into two groups corresponding to the morphology-based species identification. The blue peafowl (BPB, BPI, BPJ, BPW, WPB, WPW) formed one cluster, while green peafowl (GKR, GTL, GYN, GYUEN, GZJ) formed the second cluster (Fig. 3C and Supplementary Fig. S15). At K=3, three clusters were observed, some individuals in the green peafowl (GKR, GYUEN, GZJ) have new clusters. The character of green peafowl clusters has obvious geographical associations with their sampling locations. But there is no obvious difference at the national scale, which may be due to the fact that the green peafowl was sent to another place for conservation. Notably, the three green peafowl samples (GF-f1A, qhd-01, qhd-02) equally composed of the blue peafowl and green peafowl alleles were observed from k = 2 to 9. PCA analysis also recapitulated these findings. In the first principal component, the blue peafowl and green peafowl clusters were divided along the two eigenvectors without overlap. Three green peafowl individuals (GF-f1A, qhd-01, qhd-02) were positioned outside the discovered clusters, which deviated towards green peafowl in the first principal component (Fig. 3D).

**Fig. 3.**
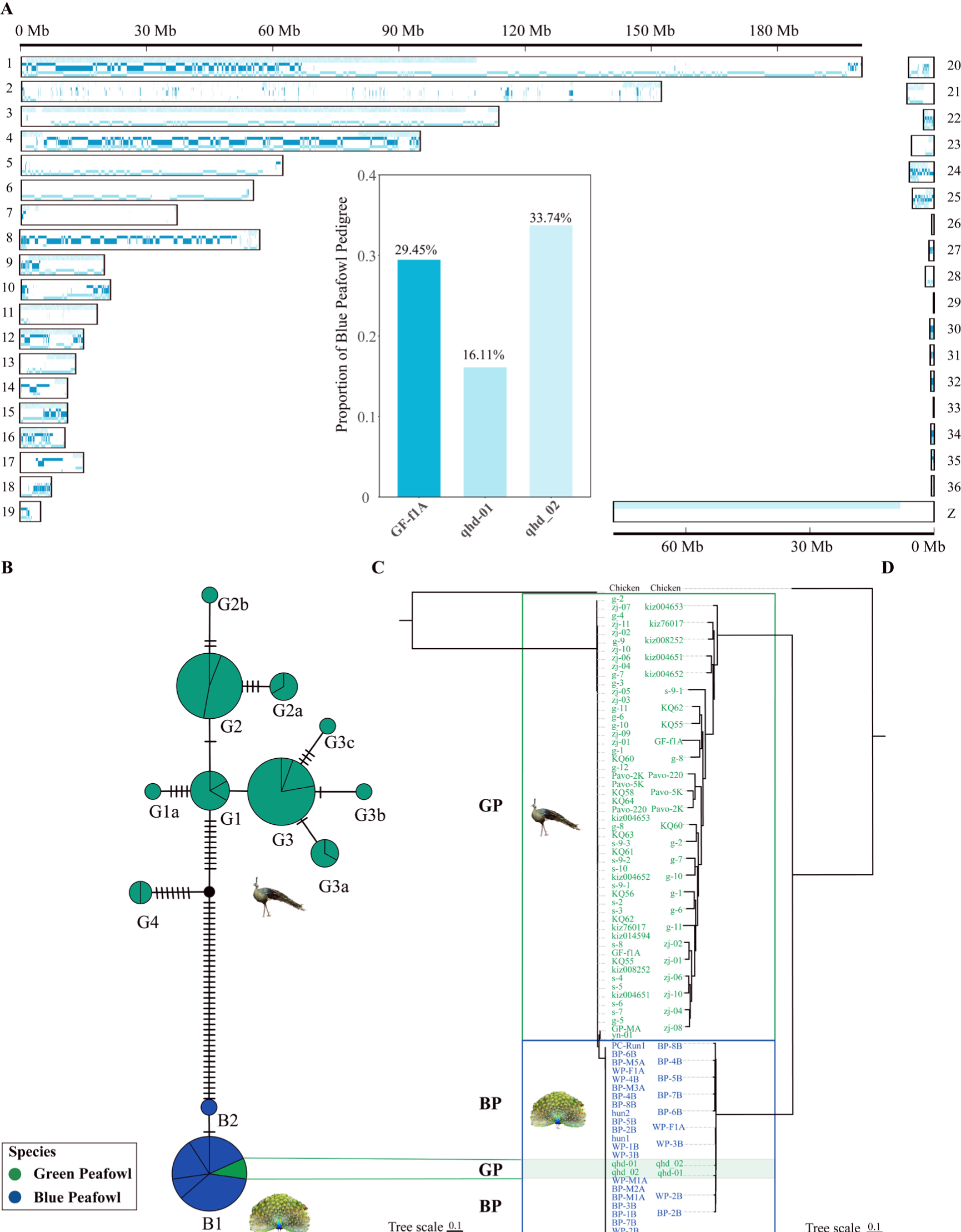
Lineage distribution of hybrid green peafowl individuals in autosomes, W chromosome and mitochondrial DNA. (A) Genome-wide distribution of haplotypes fragments between blue peafowl (blue) and green peafowl (green) in hybrid green peafowl individuals. Each row represents a diploid individual with two haplotypes stacked on top of one another. (B) The inset shows the proportion of autosomal blue peafowl ancestry in hybrid green peafowl individuals by using rfmix. (B) Median-joining haplotype network from the 76 peafowl individual mitochondrial sequence. (C) Mitochondrial phylogenetic tree of 76 peafowl individuals constructed using the ML method. (D) W chromosome phylogenetic tree of 38 female peafowls constructed using the ML method. BP represents the blue peafowl and GP represents the green peafowl.

Based on species and geographic location, we performed genetic diversity analysis inferred nucleotide diversity, the linkage disequilibrium (LD) parameters (r2) and long runs of homozygosity (ROH) for populations with sample sizes >2 (including GYUEN, GZJ, GYN, GTL, GQHD, WPB, BPW, BPB). Among the green peafowl, the GQHD population had the highest genetic diversity, while the GYUEN population had the lowest genetic diversity. BPB populations have the highest genetic diversity and WPB populations the lowest in blue peafowl. We also found that the nucleotide diversity of the green peafowl (π=0.00147241) was lower than that of the blue peafowl (π=0.00197888) (Supplementary Fig. S16). The LD decay rate of the blue peafowl was slower than green peafowl, the Yunnan green peafowl showed the fastest LD decay and the smallest r2, indicating that they had higher diversity in wild environment (Supplementary Fig. S17). The length of the long homozygous segment can reflect the degree of inbreeding of the population. Compared with the peafowl, the ROH length of all peafowl populations is short and the number is small. These results suggests that the degree of inbreeding of the green peafowl is much lower than that of the blue peafowl, which may be due to the long-term wild environment of the green peafowl (Supplementary Fig. S18).

### 3.7 Maternal phylogenetic analyses of peafowl

Both W-chromosome and mitochondrial DNA (mtDNA) haplotypes represent strong foci in the investigations of Aves. Here, we used complete mitogenomes to construct a haplotype network and rooted haplotype trees with chicken as outgroup (Fig. 3C and 3D). A total of 12 mitochondrial haplogroups (G1, G1a, G2, G2a, G2b, G3, G3a, G3b, G3c, G4 for green peafowl, and B1, B2 for blue peafowl) emerged. We found the G4 haplotypes in green peafowl that was older. We also used 745 SNPs in the W-chromosome to construct a phylogenetic tree (Fig. 3E). The evolutionary tree can clearly divide the blue peafowl and green peafowl into two branches.

There are two green peafowl individuals that caught our attention. Mitochondrial phylogenetic tree of all peafowl individuals shown that two green peafowl individuals (qhd-01, qhd_02) were in the blue peafowl clade. This is contrary to the results of autochromatic NJ trees. Combining the results of autosomal Admixture and PCA, we speculate that the two green peafowl individuals may be the individuals of the recent cross between blue peafowl and green peafowl. The phylogenetic tree of the W chromosome is another strong evidence of studying the maternal genetic relationship of peafowls. We found that the two green peafowl individuals are located on the blue peafowl branch on the W chromosome ML tree of 33 peafowl individuals. This further confirms our speculation. Although the maternal inheritance of the GF-f1A individual was in the green peafowl, we also assumed that it is also a hybrid individual because the autosomes show a similar ancestry distribution to the other two individuals.

### 3.8 Hybrid green peafowl individuals in the Zoo

In the wild, the habitats of the blue peafowl and the green peafowl do not overlap^3,4^ (Figure 2A). The number of wild green peafowls is extremely rare. It is reported that the total number of green peafowls is less than 500^14^. Although blue peafowls are not reproductively isolated from green peafowls, no hybrid green peafowls have been reported in the wild. Through phylogenetic tree analysis of euchromatic, mitochondrial and W chromosomes sequence, we discovered three putative green peafowl individuals hybridizing blue peafowls and green peafowls. Next, we will further verifiy this hybridization event and analyze the proportion of blue peafowl blood in these samples. Migration events between green peafowls and blue peafowls populations were estimated using TreeMix and constructing ML phylogenetic trees. When the migration event was set to an optimal value of 3 (M=3), gene flow from blue peafowl and green peafowl occurred (Fig. 2E). To investigate the relationship of the three green peafowl individuals to blue peafowl, we selected core groups of green peafowl respectively based on the genetic structure analysis using ADMIXTURE and a phylogenetic analysis using f3 statistics (Supplementary Table S12 and S13). Then, we performed D statistics and f3 statistics for all individuals based on SNP frequency differences. In D statistical analysis, we set (P1, P2, P3, O) as (green peafowl individuals, putative hybrid individuals, blue peafowl individuals, chicken). The D statistics showed signifcant introgression events (Z > 3, *P* < 0.001) from all putative hybrid individuals. The f3 statistics and D statistics also confirmed that the three green peafowl individuals shared the most derived polymorphisms with green peafowl. These results corroborated the PCA and ADMIXTURE results that demonstrated the existence of hybridization between blue peafowl and green peafowl.

Using the core populations of the two species as a reference, we showed that hybrid green peafowl individuals share allele SNPs and haplotype-region with the blue peafowl (Fig. 4A and Supplementary Fig. S19). Two individuals (qhd-01, qhd_02) with similar shared allele profiles and another individual (GF-f1A) who differs from them, possibly due to differences in genetic background due to geographic location. We also used the same sample to calculate the proportion of ancestry of blue peafowl among the three hybrid individuals (Fig. 4A and Supplementary Fig. S20). Among them, the proportion of ancestry of blue peafowl in qhd_02 individual accounted for the highest 33.74%, which indicated that this individual was a hybrid individual of one or two generations of blue peafowl to green peafowl. Although the mitochondrial DNA of GF-f1A individual is in the green peafowl clade, the proportion of ancestry of blue peafowl in its genome reached 29.45%. Phylogenetic relationships inferred from ML trees constructed using mitochondrial SNPs and W chromosome SNPs (Fig. 4B, 4C and 4D) further supported these findings, indicating distinct clusters of green peafowl individuals and blue peafowl, whereas two green peafowl individuals (qhd-01, qhd_02) is located in the blue peafowl branch.

**Fig. 4.**
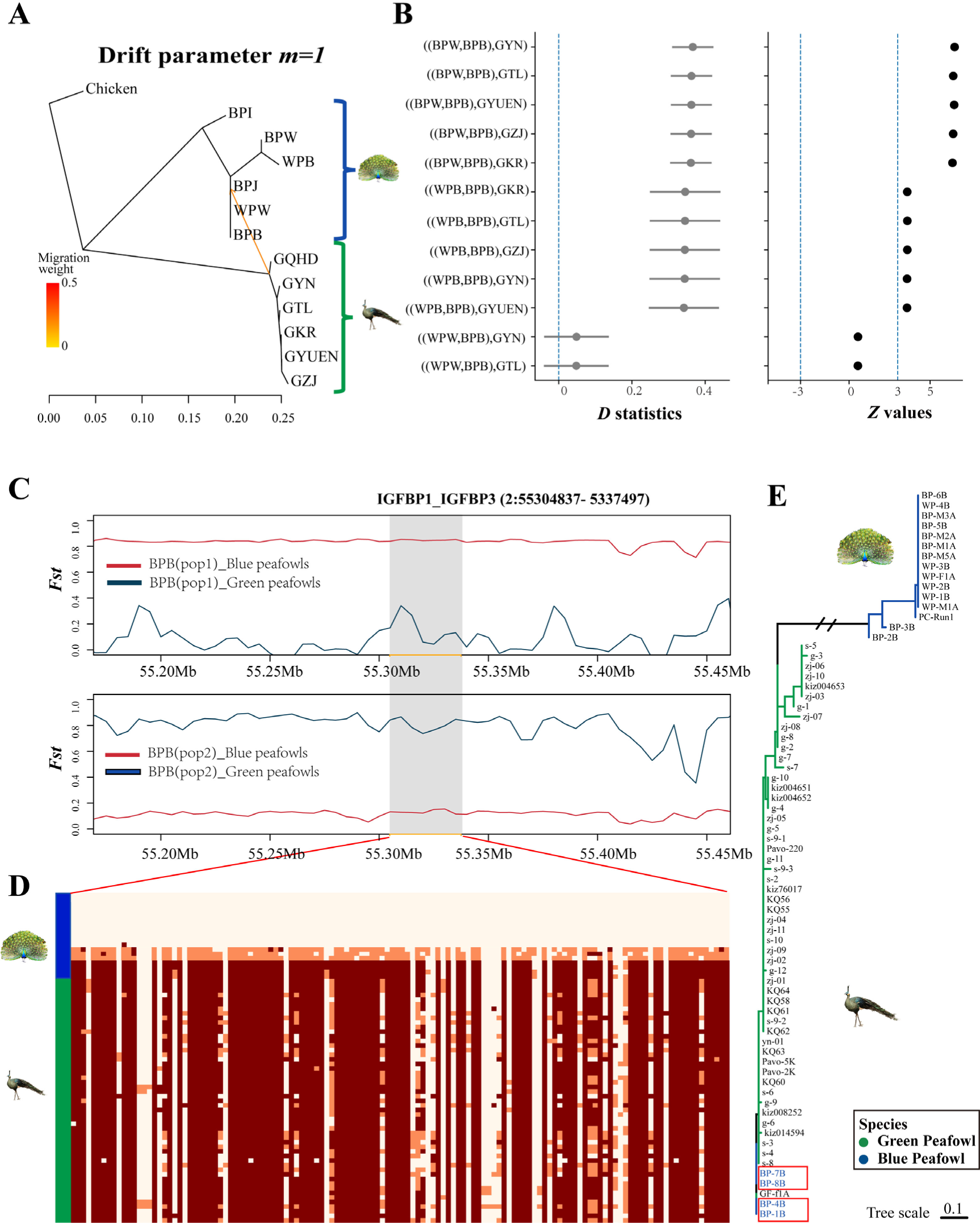
Introgression analysis of peafowl population. (A) Genetic migration inferred using TreeMix (M=1). (B) D-statistics for introgression between BPB blue peafowl population and green peafowl population. (C) *Fst* between BPB blue peafowl individuals and the green peafowl populations. The BPB blue peafowl samples were divided into two groups based on *Fst* values, BPB (pop1) includes BP-1B, BP-4B, BP-7B and BP-8B; BPB (pop2) includes BP-2B, BP-3B, BP-5B and BP-6B. The blue line represents *Fst* between BPB and green peafowl, and red line represents *Fst* between BPB and blue peafowl. The gray region indicates the location of the IGFBP1 and IGFBP3 gene regions. (D) Introgression plot of the blue peafowl contacted based on shared allele SNPs in IGFBP1 and IGFBP3 gene regions across chromosome2 (from 55304837 bp to 55337497 bp). (E) Phylogenetic tree of 77 peafowl individuals constructed by 109 SNPs in IGFBP1 and IGFBP3 gene regions using ML method. The blue peafowl individuals in the red box have introgression in this region.

### 3.9 Historical introgression regions influence Blue Peafowl body size and immunity

Treemix analysis of all individuals not only showed the gene flow from blue peafowl to green peafowl, but the gene flow from green peafowl to Beijing blue peafowl (Fig. 2E). Therefore, blue peafowls and green peafowls may have historical infiltration events. We removed the individuals identified as hybrids, and the calculated treemix showed the gene flow from green peafowl to Bejing blue peafowl occurred when the migration event was set to 1(M=1) (Fig. 4A). To further analyze introgression events, we performed f3 statistics and D statistic based on SNP frequency difference. Both of the two methods showed significant introgression events (Z > 3) (Fig. 4B).

The introgression region was further defined by identifying 267 genes in the top 5% D-value window positions and presuming these to be candidates for introgression (Supplementary Fig. S21 and Table S14). The GO and KEGG annotation showed that the introgression genes were enriched for significantly introgression genes mainly included cytoplasm (GO:0005737), cellular senescence KEGG PATHWAY (gga04218) (Supplementary Fig. S22). Among these genes, IGFBP1 and IGPBP3 which is a member of the Insulin-like growth factor-binding protein (IGFBP) family, were highlighted because of associating with body size^128,129^.

The degree and direction of introgression in the BPB blue peafowl was further detected by calculating the *Fst* between BPB blue peafowl individuals and other blue peafowl, BPB blue peafowl individuals and green peafowl individuals, respectively, based on the results of D statistic analysis (Fig. 4B). We obtained a region located on chromesome 2 (from 55304837 bp to 55337497 bp), which contained the candidate gene IGFBP1 and IGPBP3. In the region, the BPB pop1 (BP-1B, BP-4B, BP-7B, BP-8B) and green peafowl were poorly differentiated (*Fst* = 0), BPB pop2 (BP-2B, BP-3B, BP-5B, BP-6B) and blue peafowl were highly differentiated (*Fst* = 0.89) (Fig. 4C). An ML tree was constructed using IGFBP1 and IGPBP3 gene sequences (Fig. 4E). In the ML tree, BP-1B, BP-4B, BP-8B, BP-7B clustered with green peafowl branch and other blue peafowl formed separate branches. Haplotypes in this region also showed similar results (Fig. 4D and 4E).

In addition, we retrieved several immune response related genes (IL6, IL12B, IL25, NFATC1, DROSHA and CUL1) in introgressed regions that may help blue peafowl fight disease. Among all immune genes, IL-6 gene is involved in multiple immune pathways and is a chicken heat shock protein^130,131^. Studies have found that IL-6 is involved in the innate immunity of poultry diseases such as Newcastle disease (ND)^132^.

### 3.11 Nonsense mutation in EDNRB2 gene creates white plumage individuals in blue peafowls

As early as 1868, Darwin reported the white peafowls and pied peafowl (blue white-flight plumage color), the mutant individuals of the blue peafowl’s plumage color, which shows that the white peafowl appeared at least 150 years ago^133^ (Fig. 5E and 5F). Under artificial breeding, white peafowls have reached a state of self-sustaining population. As a striking ornamental bird, research on the genetic mechanism of plumage color in white peafowls has never ended. Studies have reported that the recessive allele that controls the white feathers of peafowls is located on an autosomal chromosome, and its inheritance conforms to Mendel’s law of segregation^33^. However, the molecular mechanism of white peafowl plumage color has not been elucidated due to technical limitations^32^.

**Fig. 5.**
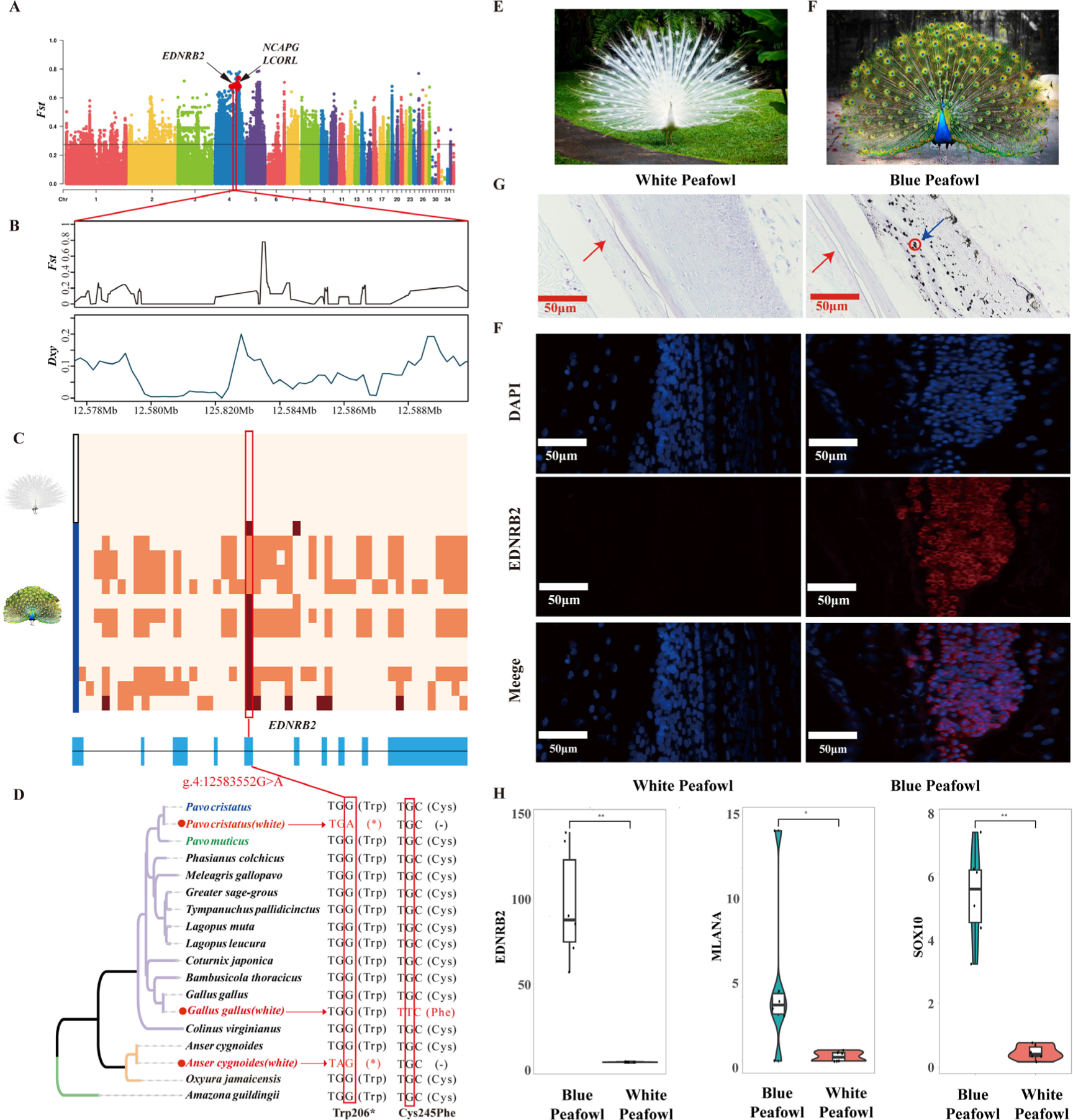
White peafowl white plumage phenotype characteristics and molecular mechanism. A. Whole genome scan with *Fst*. The red arrow indicates the genome position of the strongest selective signal for the white plumage phenotype. B. *Fst* and Dxy corresponding to the white plumage selective sweep on chromosome 4 which encompasses the EDNRB2 gene. C. Plot of the haplotype structure of SNPs around the EDNRB2 gene in all white peafowls and blue peafowls. Nucleotides with brown box represent blue peafowl homozygote genotypes, and with orange box represent heterozygote genotypes. The plot also showed the genetic structure of EDNRB2, and the blue box is the CDS region. The position of the red line indication is the nonsense mutation of the w white plumage phenotype (g.4:12583552G>A), which is located in the fifth CDS region. D. The convergence evolution of EDNRB2 codons in different white plumage species comparison. The first convergent site is Trp206*(Stop codon). The white peafowl has a nonsense mutation at this position. White geese have a 14-BP insertion before the position, so that the codon is turned to be a stop codon in this position. The second convergent site is Cys245Phe, which only occurs on the white plumage chickens. E. A white peafowl with white plumages and black eyes. F. A Blue peafowl with black and blue plumages. G. Micrographs of plumage sections from white and blue peafowls stained with the Masson-Fontana. The red arrow is the plumage tube wall. Blue arrows are melanosome and melanocytes. F. (F) FISH was performed to observed the cellular location of EDNRB2 in plumage tube. Nuclei were stained with DAPI. Scale bar = 50 µm. G. EDNRB2 and melanocyte maker genes (SOX10 and MLANA) are expressed in white peafowl and blue peafowl plumages, respectively. The genetic expression is measured by transcripts per million (TPM). Data were indicated as mean ±SEM (n = 3), ns *P* ≥ 0.05, * *P* < 0.05, * * *P* < 0.01.

In this study, we compared 11 blue peafowl samples and 5 white peafowl samples from different geographic regions to identify the genomic region responsible for the emergence of mutated white peafowl individuals. We used *Fst* method to search for the causal mutations. The most significant region was located on the 12–13 Mb interval of chromosome 4 (Fig. 5A, 5B and Supplementary Table. S15). The analysis based on Absolute genetic divergence (Dxy) and Tajima’s D supported a strong selective signal in this region. Through genotypic analysis of 159 mutations (SNP, Indel and CNV) in this interval, we found that the only polymorphism completely associated with peafowl white plumages was a nonsense mutation site located in the EDNRB2 coding region (g.4:12583552 G>A) (Fig. 5C and Supplementary Fig. S23). EDNRB2 is an important key gene in the formation of bird feather color. EDNRB2 is one of the receptors for EDNs, which is strong mitogens for melanoblasts. Kawasaki-Nishihara et al. suggested that EDN3–EDNRB2 signaling is required for normal melanoblast migration in Xenopus embryos on the basis of in vivo experiments^134^. It does not participate in the melanin synthesis pathway, but regulates the differentiation, proliferation, and migration of melanocytes^135,136^. Therefore, we speculated that this incorrect genetic information leads to RNA degradation, known as NMD (nonsense-mediated mRNA decay)^137^. The degradation of EDNRB2 mRNA inhibits the differentiation, proliferation and migration of peafowl melanocytes, resulting in the white peafowl individuals with white plumages and black eyes.

To confirm this assumption, we analyzed the genotypes of 11 white peafowls, 29 blue peafowls, and one pied peafowl. This SNP showed homozygous or heterozygous genotypes (G/G or G/A) in all blue peafowls and heterozygous genotypes (G/A) in pied peafowl, while all white peafowls were homozygous for the mutant (A/A) (Supplementary Table S16). These results indicate that the white plumage phenotype emerged as a result of the nonsense mutation in the EDNRB2 gene and the white allele (A) is shown to be a incompletely dominant. RNA-seq data of three blue peafowls and three white peafowls plumage found that there was no expression of the EDNRB2 gene in the white peafowl plumage, while the gene was highly expressed in the blue peafowl feathers. FISH (Fluorescent in situ hybridization) experiments on this tissue showed the same results (Fig. 5F). In addition, we also found that the melanocyte marker genes (SOX10 and MLANA) are not expressed in white peafowl plumage^138^, but are highly expressed in blue peafowl plumage (Fig. 5H and Supplementary Fig. S24). The formation and deposition of melanin mainly occurs on the amyloid fibres of melanosomes. Masson-Fontana staining found no melanin and melanocytes in white peafowl plumage tissue (Fig. 5G). The above results indicate that melanocytes do not exist in white peafowl feathers. This verified our conjecture that the nonsense mutation of the EDNRB2 gene caused the NMD reaction in white peafowls, and the mRNA of the EDNRB2 gene was degraded during translation, thus preventing melanocytes from being transported into plumage tissue.

### 3.12 Convergent selection for the White feather in geese, chicken and peafowls

The white plumage trait in birds represents a common phenotypic convergence, particularly observable in poultry breeds^139,140^. In these species, various colorful birds have been domesticated, leading to the emergence of white plumage individuals due to the domestication syndrome^141^. However, the molecular mechanisms underlying this white plumage trait differ across species. For instance, in waterfowl, white plumage in duck results from alternative splicing of the MITF gene^142,143^, whereas in geese, it is caused by mutations in the EDNRB2 gene^35^. In chickens, dominant white plumage is governed by the PMEL 17 gene^140^, while Tyrosine-Independent white plumage is controlled by the EDNRB2 gene^34^. Notably, both white plumage chickens and geese regulated by the EDNRB2 gene exhibit predominantly white plumage across their bodies but possess black eyes.

In this study, in order to explore the underlying molecular regulatory mechanism of this phenotypic convergence, we compared the EDNRB2 gene sequences in 15 species of birds, including white plumage chickens and white plumage geese. In the case of Minohiki chickens, a nonsynonymous mutation (G>T) associated with the white feather phenotype was identified in the coding region (CDS) of the EDNRB2 gene^34^. This mutation leads to a functional defect in EDNRB2’s ability to bind to EDN ligands, which in turn interferes with melanocyte differentiation, proliferation, and migration (Fig. 5D). In the Gang geese, a 14-base pair insertion caused a frameshift mutation in the gene’s coding region, resulting in a premature stop codon (TAG)^35^. This insertion also triggered the NMD (nonsense-mediated decay) mechanism, leading to the absence of detectable mRNA in white geese individuals.

Although these findings highlight the involvement of the same causative gene (EDNRB2) is involved, the mutations responsible for white feather coloration in the three species occurred independently after the species diverged. Notably, the mRNA expression of EDNRB2 in geese is also lost due to nonsense mutations caused by frameshift mutations, thus indicating that the molecular mechanisms of white feathers in peafowls and geese are very similar.

## 4. Discussion

In this study, we report the first chromosome-level de novo genome assembly for the blue peafowl with the 95 Mb Scaffold N50 of the assembly which was substantially higher than those obtained in previous studies and green peafowl genome^8,32,78^. We obtained more pairs of macrochromosomes than in karyotype study of the blue peafowl, which only had 8 pairs of macrochromosomes^113^. The genome synteny and collinearity analysis with chicken and other birds and the evaluation of various algorithms prove that the current blue peafowl genome assembly is of high quality, with consistency, accuracy and completeness. Although we achieved partial chromosome-level assembly, due to the GC-richness, specific repeats, high microchromosome mutation rate in birds^144^, and lack of cytogenetic support, the peafowl microchromosomes and W chromosome require a combination of different data sources including optical mapping^145^ and linked reads^146^ to improve the assembly quality. In general, this chromosome-level genome of blue peafowl strongly supports the subsequent comparative genomic and population structure analysis.

The karyotypes of birds are relatively conserved, especially in the pheasant family^147,148^. This pattern was confirmed in our analysis of the conserved synteny between turkey and chicken. We only detected chromosome fusions on chromosomes 6 and 8. However, comparisons for micro-chromosomes are less accurate due to the more incomplete assembly of these other Galliform species.

Regarding the relationship between peafowl and chicken and turkey, there are two views: (1) Peafowl is closer to turkey than chicken^32,149^ (2) Peafowl is closer to chicken than turkey^9,78,150^. To resolve previous inconsistencies concerning peafowl phylogenetic relationships, a new phylogenomic tree was reconstructed from a total of one-to-one orthologs based on a combination of the concatenation method and the coalescent method. We also obtained the above two results. In the autosomal phylogenetic relationship, peafowls are closer to turkeys, while mitochondrial sequences show that peafowls are closer to chickens. We found that the appearance of peafowls (∼37 million years ago) coincided with the rapid speciation of pheasants based on multiple species fossil time correction or site spectrum. Rapid speciation have occurred many times during the evolution of birds, and these events have resulted in a large number of incomplete lineage sorting (ILS) among bird species. Combined with the timing of the peafowl’s emergence (∼37 million years ago) and a number of phylogenetic inconsistencies across the genome, we believe that ILS is also responsible for the huge controversy over the phylogenetic relationships of peafowls with chickens and turkeys.

Our results indicated that blue peafowl and wild green peafowl have different population histories. The wild green peafowl experienced rapid population expansion followed by a dramatic population decline during the LGP which consistent with the results of Feng et al^18^. And the most other threatened species experienced a similar history of group dynamics^151–155^. The effective group size of blue peafowl is higher than that of blue peafowl before LIG. Moreover, the effective group size of the blue peafowl gradually became smaller after the LIG, which is very similar to the historical dynamics of domestic chickens^156^. The different group histories of the two species may be an important factor in the current divergent fate of the two species. Although the wild green peafowl experienced a significant decrease of genetic diversity in the last 50 years^18^, the genetic diversity of wild green peafowl populations is still significantly higher than that of captive blue peafowl.

Habitat loss and fragmentation has long been considered the primary cause for biodiversity loss and wild animal extinction worldwide^157^. Over the past three decades, the endangered wild green peafowl as the only peafowl native to China, has experienced sharp population declines and faced the problem of habitat fragmentation^14,15^. In order to respond to the Convention on Biological Diversity and save the endangered wild green peafowl^158^, the Chinese government has issued a number of protection policies, including banning poaching and captive breeding^21,159^. The wild green peafowl can hybridize with blue peafowl and produce fertile off-spring that can backcross with ancestral species^160^. However, since the habitats of the wild green peafowl and wild blue peafowl do not overlap, and the number of green peafowl is extremely small, no hybrid green peafowls have been reported in the wild^3,14^. Preventing blue peafowls from interbreeding with green peafowls is critical to the conservation of endangered green peafowls. Some researchers in Southeast Asia suggested that the breeding of blue peafowl near the distribution area (including the potential distribution area) of the wild green peafowl should be prohibited^161^. The blue peafowl and the green peafowl no longer have geographical isolation and hybridization may occur in Zoo. Through autosomal and mitochondrial analysis of peafowl populations and validation of multiple methods, we found three green peafowl individuals crossed with blue peafowl. The blood ratio of the blue peafowl among the three green peafowl individuals is less than 50%. Obviously, these three peafowls have undergone backcrossing after hybridization. Moreover, consistent with the description of Du et al., hybrid green peafowls were indistinguishable from purebred green peafowls in appearance^160^. Hybridization is a conservation threat for endangered species, such as red wolf Canis rufus hybridizing with coyote Canis latrans, bison Bison bison with domestic cattle (Bos spp.) or banded pig (Sus scrofa vitattus) with the endangered Java warty pig (Sus verrucosus)^25–28^. Captive breeding for reintroduction is potentially an important measure for localized population restoration^160^. However, it is critical to prevent hybrid green peafowl individuals from entering the wild. If there are hybrid individuals among the released green peafowl, purebred green peafowl individuals in the wild are likely to cause pollution and eventually lead to extinction. Therefore, we appeal to prevent the hybridization of green peafowls and blue peafowls when we call for artificial breeding of green peafowls. At the same time, we suggest that purebred identification must be carried out before releasing green peafowls to prevent genetic contamination.

In addition to recent hybridization, we also found historical introgression events between blue peafowl and green peafowl, with blue peafowl having gene flow from green peafowl. Interbreeding is a widespread event among birds^162^, and the habitats of blue peafowl and green peafowl overlap historically, so our study confirms their past interbreeding. Our analyses identifed multiple regions of putative green peafowl derived ancestry in blue peafowl, which contained 267 candidate protein-coding genes. The sequence of IGFBP1 and IGFBP3 had the strong signals among these genes. Both of the genes which conserved in birds are member of the insulin-like growth factor binding protein (IGFBP) family and encodes a protein with an IGFBP domain and a thyroglobulin type-I domain^129^. The IGFBP family mediates IGF effects by enhancing or dampening IGF signaling. This occurs by either increasing IGF-receptor affinity, physically sequestering it to prevent receptor binding, or extending IGF’s half-life in circulation^128^. Additionally, many IGFBPs can act independently to induce cellular activity^163^. A large number of studies have found that IGFBP3 affects growth traits in common domestic animals, such as pig^164^, cattle^165^ and sheep^166^. Moreover, the IGFBPs system is highly conserved in chicken, and is involved in the regulation of egg production, growth, and carcass traits. IGFBP3 participates in myogenic cell proliferation and myoblast differentiation, and the SNPs in the IGFBP3 promoter region were significantly associated with body weight, breast muscle weight, and leg muscle weight^167^. In addition, the genes (IGF2BP3, TGBR1, ISPD, MEOX2, GLI3 and MC4R) related to body size in blue peafowl were also found to have introgression areas from green peafowl. The size of the peafowl is larger in the pheasant family, and the size of the green peafowl is even larger than that of the blue peafowl^168^. These body size-related genes may be an important reason for the current larger size of the blue peafowl, but whether it is a factor affecting the viability of the blue peafowl needs further research. The immune-related genes found in introgressed regions are likely the result of adaptive introgression which possibly improved the viability of the blue peafowl, consistent with studies in common domesticated animals such as chickens and cattle^169,170^.

Molecular genetic studies in a variety of organisms highlight the repeatability of phenotypic change by changes in the same genes^171,172^. Among vertebrates, this trend is especially pronounced in studies of pigmentation diversity^173,174^. One dramatic example of this repeatability is the dilution of the coloration typical of the Dun phenotype displays very similar microscopic and macroscopic features in the Perissodactyla horses and donkeys^175^. TBX3 is responsible for the Dun pattern of pigmentation in both species. The causal mutation of the non-Dun phenotype in donkeys is a 1 bp deletion with a probable regulatory effect. Similarly, in horses, the non-Dun phenotype is explained by two deletions with regulatory effects^176^. In another study, Lopes et al. provided evidence that the yellow and red plumages pigmentation of canaries and finches is caused by C(4)-oxygenation of carotenoids by the cytochrome P450 enzyme CYP2J19^177^. This chemical modification increases the length of the conjugated part of the carotenoid molecule, causing a red shift in its absorbance spectrum^178^. These and other examples also highlight an emerging pattern: Not only are the same genes involved in convergent evolution among populations and species, but so are specific molecular regions^179^. Here, we showed that the EDNRB2 gene was convergently selected in blue peafowls, chickens and geese. The three types of white-feathered birds have different mutation types in the same gene. The reason for the white feathers in chickens is that the protein molecular structure of the EDNRB2 gene has changed^34^. Although the mutation sites in peafowls and geese are different, they ultimately cause premature termination of codons which triggering the NMD molecular mechanism of organisms and preventing melanocytes from being transported to plumages^35^. Our results demonstrate that not only the same gene regions are involved in convergent evolution between populations and species, but also specific molecular response mechanisms resulting from mutations in this region are involved. Although this particular case cannot be generalized to other phenotypes, it emphasizes the need to precisely clarify the role of convergent evolution in the fixation of Mendelian phenotypes.

## 5. Conclusions

In conclusion, we achieved a high-resolution, chromosome-level genome assembly for the blue peafowl, which has significantly contributed to resolving the controversy surrounding the phylogenetic position of peafowl within the Phasianidae family. Our findings indicate that the blue peafowl and the green peafowl diverged approximately 12 to 13 million years ago, and their relationship is closer to the turkey rather than the chicken. Furthermore, we also demonstrate different demographic trajectories in the two species: the population history of the wild green peafowl is closer to that of endangered animals, whereas the population history of the blue peafowl is closer to that of domestic birds. In addition, we demonstrated not only historical introgression but also recent hybridization between these green peafowl and blue peafowl. The historical hybrid introgression has been found to impact body size and immunity in blue peafowls. Notably, we found three individuals of green peafowl crossed with blue peafowl in zoos based on genome-wide data, which confirmed that the two species can interbreed and produce fertile offspring. Therefore, in order to better protect the endangered wild green peafowl, we call for preventing the occurrence of hybridization between green peafowl and blue green peafowl as soon as possible. Moreover, we suggest that purebred identification must be carried out before releasing green peafowl to prevent hybrid green peafowl from entering the wild. We also elucidate the tyrosine-independent molecular mechanisms that control white peafowl plumage color. We speculate that the emergence of different white plumage colors is the result of multiple independent evolutions. Our research provides both theoretical and empirical support for the conservation of the endangered green peafowl, as well as the sustainable management and breeding of the blue peafowl population.

## Data availability statement

The datasets presented in this study can be found in online repositories. The genome assembly and corresponding sequencing data were deposited at CNGB Sequence Archive (CNSA) of China National GeneBank DataBase (CNGBdb) with accession number CNP0004612. The whole sequencing data and green peafowl Hi-C data collected were from the BioProject accession number CNP0002498, PRJNA665082, PRJNA340135 and PRJNA644939. RNA sequencing data collected were from the BioProject accession number PRJNA661158 and PRJNA271731.A full list of accession IDs is available in the Supplementary Table S8.

## Additional Files

### Supplementary Figure

Supplementary Fig. S1. Pipeline of the draft genome assembly and genome annotation of blue peafowl (WP-1).

Supplementary Fig. S2. Estimation of the genome size of the blue peafowl by K-mer analysis. Supplementary Fig. S3. Hi-C-based chromosome-level assemblies of blue peafowl genome and green peafowl.

Supplementary Fig. S4. BUSCO assesses the completeness of published genomes and this assembled genome.

Supplementary Fig. S5. Genome synteny and collinearity among the blue peafowl and Turkey. Supplementary Fig. S6. Mitochondrial genome map of blue peafowl. Genes that belong to different functional groups are color-coded.

Supplementary Fig. S7. Coalescent species tree inferred from single-copy orthologous gene trees by ASTRAL.

Supplementary Fig. S8. ML phylogenetic tree of 15 birds inferred using single-copy orthologous genes by using iqtree.

Supplementary Fig. S9. ML phylogenetic tree and molecular clock dating analysis of 11 birds based on mitochondria genomes.

Supplementary Fig. S10. Population size history inference of blue peafowl (BP) and green peafowl (GP).

Supplementary Fig. S11. Group history dynamic simulation of blue peafowl and green peafowl by using DaDi.

Supplementary Fig. S12. KEGG pathway enrichment analysis of the expansion gene family of peafowl.

Supplementary Fig. S13. KEGG pathway enrichment analysis of PSG and RGs of peafowl.

Supplementary Fig. S14. Variation annotation information of all peafowl individual.

Supplementary Fig. S15. Group Structure of blue peafowl and green Peafowl. Supplementary Fig. S16. Nucleotide diversity of blue peafowl and green peafowl.

Supplementary Fig. S17. Linkage disequilibrium in peafowl groups with individuals greater than two.

Supplementary Fig. S18. Runs of homozygosity (ROH) in peafowl groups with individuals greater than two.

Supplementary Fig. S19. Each chromosome distribution of shared alleles SNPs between blue peafowl (blue) and green peafowl (green) in hybrid green peafowl individuals.

Supplementary Fig. S20. Comparison of hybrid individual green peafowl and purebred green peafowl.

Supplementary Fig. S21. Manhattan plot of BPB blue peafowl group introgression with a window of 2500 SNPs and a step size of 500 SNPs (P1, P2, P3, O).

Supplementary Fig. S22. Enrichment analysis of green peafowl introgression regions in BPB blue peafowl group.

Supplementary Fig. S23. Caused mutation of the EDNRB2 gene locus (chr4: g.12583552 G>A) for the white plumage trait in white peafowls.

Supplementary Fig. S24. Bar plot of differential mRNA expression (log2-transformed fold change) and (Transcripts per million (TPM)) of pigmentation-related genes expressed in the plumage of white peafowls (n = 6) versus blue peafowls (n = 6).

### Supplementary Tables

Supplementary Table S1. Statistics of genome assembly data of blue peafowl (WP-1).

Supplementary Table S2. Summary of de novo genome assembly of blue peafowl.

Supplementary Table S3. Assembly assessment of completeness using BUSCOs.

Supplementary Table S4. Statistics of repeats in our assembled genome.

Supplementary Table S5. Statistics of non-coding RNAs in the assembly of peafowl.

Supplementary Table S6. Four model paraments simulated group history dynamic of blue peafowl and green peafowl by using DaDi.

Supplementary Table S7. PSG (positive selection gene) of peafowl by using the branch-site models.

Supplementary Table S8. PSG (positive selection gene) and RGs (rapidly evolving genes) of green peafowl by using the branch-site models and branch models.

Supplementary Table S9. PSG (positive selection gene) and RGs (rapidly evolving genes) of blue peafowl by using the branch-site models and branch models.

Supplementary Table S10. All samples information.

Supplementary Table S11. Sequencing depth and coverage of all samples.

Supplementary Table S12. f3-statictis of peafowl groups.

Supplementary Table S13. “core” group individuals of green peafowl and blue peafowl by using f3 statistics and ADMIXTURE.

Supplementary Table S14. D statistics of introgression regions (BPW, BPB, GYN, chicken).

Supplementary Table S15. *Fst* value of blue peafowl and white peafowl.

Supplementary Table S16.All variations within 10kb before and after the SNP (chr4: g.12583552 G>A).

Supplementary Table S17. Genotype distribution of the short deletion at chromosome 4 (chr4: g.12583552 G>A) in Blue and White peafowls.

## Acknowledgments

We would like to thank Professor Judith Mank, Department of Zoology, University of British Columbia, for her valuable comments and suggestions on this paper. Thanks to the Beijing Zoo for providing the peafowl blood samples and photos. Thanks for the supporting by High-performance Computing Platform of China Agricultural University.

## Author contributions

G.W. ‘contributed’ Data curation, Formal analysis, Writing and editing; Z.N. and R.X. ‘contributed’ Y,” Investigation. L.B., X.Z., L.Z., Y.L., Z.J., J.W., X.Z., Y.Z., X.C. and Y.L. ‘contributed’ Collect samples; X.R. ‘contributed’ Data curation; A.C. and W.D. ‘contributed’ Methodology; L.Q. ‘contributed’ Project administration, supervision, Validation and Writing—review.

## Ethics approval and consent to participate

This study conformed to protocols approved by the Laboratory Animal Welfare and Animal experimentation Ethics Review Committee of China Agricultural University (No. XK622).

## References

1. Liu, S. et al., A high-quality assembly reveals genomic characteristics, phylogenetic status, and causal genes for leucism plumage of Indian peafowl. GIGASCIENCE 11 (2022).

2. Gadagkar, R., Is the peacock merely beautiful or also honest? CURR SCI INDIA 1012 (2003).

3. Kushwaha, S. & Kumar, A., A review on Indian peafowl (Pavo cristatus) Linnaeus, 1758. J Wildl Res 4 42 (2016).

4. Desai, A., Cry, the peacock. (Orient paperbacks, 1983).

5. Fontana, D.,, The secret language of symbols: A visual key to symbols and their meanings. (Chronicle Books, 2003).

6. Hernowo, J. B., Mardiastuti, A., Alikodra, H. S. & Kusmana, C., Behavior ecology of the javan green peafowl (Pavo muticus muticus Linnaeus 1758) in Baluran and Alas Purwo national park, East Java. HAYATI Journal of Biosciences 18 164 (2011).

7. Jaiswal, S. K. et al., Genome Sequence of Peacock Reveals the Peculiar Case of a Glittering Bird. FRONT GENET 9 392 (2018).

8. Zhang, X. et al., Chromosome-Level Genome Assembly of the Green Peafowl (Pavo muticus). GENOME BIOL EVOL 14 (2022).

9. Dhar, R. et al., De novo assembly of the Indian blue peacock (Pavo cristatus) genome using Oxford Nanopore technology and Illumina sequencing. GIGASCIENCE 8 (2019).

10. Jaiswal, S. K. et al., Genome Sequence of Peacock Reveals the Peculiar Case of a Glittering Bird. FRONT GENET 9 392 (2018).

11. Harrison, P. W. et al., Sexual selection drives evolution and rapid turnover of male gene expression. P NATL ACAD SCI USA 112 4393 (2015).

12. Ramesh, K. & McGowan, P., On the current status of Indian Peafowl Pavo cristatus (Aves: Galliformes: Phasianidae): keeping the common species common. Journal of Threatened Taxa 1 106 (2009).

13. McGowan, P. et al., Handbook of the birds of the world alive. (2019).

14. Tang, W., Wang, X., Yan, M., Zeng, G. & Liang, J., China’s dams threaten green peafowl. SCIENCE 364 943 (2019).

15. Kong, D. et al., Status and distribution changes of the endangered Green Peafowl (Pavo muticus) in China over the past three decades (1990s‒2017). Avian Research 9 1 (2018).

16. McGowan, P. et al., A review of the status of the Green Peafowl Pavo muticus and recommendations for future action. BIRD CONSERV INT 8 331 (1998).

17. Pang, B., Birds in Compendium of Materia Medica. Chinese Journal of Zoology 2 35 (1976).

18. Dong, F. et al., Population genomic, climatic and anthropogenic evidence suggest the role of human forces in endangerment of green peafowl (Pavo muticus). P ROY SOC B-BIOL SCI 288 20210073 (2021).

19. Arenas, M., Ray, N., Currat, M. & Excoffier, L., Consequences of range contractions and range shifts on molecular diversity. MOL BIOL EVOL 29 207 (2012).

20. Zhou, T. C., Sha, T., Irwin, D. M. & Zhang, Y. P., Complete mitochondrial genome of the Indian peafowl (Pavo cristatus), with phylogenetic analysis in phasianidae. Mitochondrial DNA 26 912 (2015).

21. Gu, B. & Wang, F., A review on the ecology and conservation biology of green peafowl (Pavo muticus). Biodiversity Science 29 1554 (2021).

22. Lacy, R. C., Importance of genetic variation to the viability of mammalian populations. J MAMMAL 78 320 (1997).

23. Nei, M., Maruyama, T. & Chakraborty, R., THE BOTTLENECK EFFECT AND GENETIC VARIABILITY IN POPULATIONS. EVOLUTION 29 1 (1975).

24. Rhymer, J. M. & Simberloff, D., Extinction by hybridization and introgression. Annual review of ecology and systematics 27 83 (1996).

25. Adavoudi, R. & Pilot, M., Consequences of hybridization in mammals: A systematic review. GENES-BASEL 13 50 (2022).

26. Adams, J. R., Kelly, B. T. & Waits, L. P., Using faecal DNA sampling and GIS to monitor hybridization between red wolves (Canis rufus) and coyotes (Canis latrans). MOL ECOL 12 2175 (2003).

27. Freese, C. H. et al., Second chance for the plains bison. BIOL CONSERV 136 175 (2007).

28. Drygala, F., Rode-Margono, J., Semiadi, G., Wirdateti & Frantz, A. C., Evidence of hybridisation between the common Indonesian banded pig (Sus scrofa vitattus) and the endangered Java warty pig (Sus verrucosus). CONSERV GENET 21 1073 (2020).

29. Harrison, R. G. & Larson, E. L., Hybridization, introgression, and the nature of species boundaries. J HERED 105 Suppl 1 795 (2014).

30. Poelstra, J. W. et al., The genomic landscape underlying phenotypic integrity in the face of gene flow in crows. SCIENCE 344 1410 (2014).

31. Racimo, F., Sankararaman, S., Nielsen, R. & Huerta-Sanchez, E., Evidence for archaic adaptive introgression in humans. NAT REV GENET 16 359 (2015).

32. Liu, S. et al., A high-quality assembly reveals genomic characteristics, phylogenetic status, and causal genes for leucism plumage of Indian peafowl. GIGASCIENCE 11 (2022).

33. Somes R. G. Jr & Burger, R. E., Inheritance of the white and pied plumage color patterns in the Indian peafowl (Pavo cristatus). J HERED 84 57 (1993).

34. Kinoshita, K. et al., Endothelin receptor B2 (EDNRB2) is responsible for the tyrosinase-independent recessive white (mo(w)) and mottled (mo) plumage phenotypes in the chicken. PLOS ONE 9 e86361 (2014).

35. Xi, Y. et al., A 14-bp insertion in endothelin receptor B-like (EDNRB2) is associated with white plumage in Chinese geese. BMC GENOMICS 21 162 (2020).

36. Marcais, G. & Kingsford, C., A fast, lock-free approach for efficient parallel counting of occurrences of k-mers. BIOINFORMATICS 27 764 (2011).

37. Ranallo-Benavidez, T. R., Jaron, K. S. & Schatz, M. C., GenomeScope 2.0 and Smudgeplot for reference-free profiling of polyploid genomes. NAT COMMUN 11 1432 (2020).

38. Cheng, H., Concepcion, G. T., Feng, X., Zhang, H. & Li, H., Haplotype-resolved de novo assembly using phased assembly graphs with hifiasm. NAT METHODS 18 170 (2021).

39. Durand, N. C. et al., Juicer Provides a One-Click System for Analyzing Loop-Resolution Hi-C Experiments. CELL SYST 3 95 (2016).

40. Zhou, C., McCarthy, S. A. & Durbin, R., YaHS: yet another Hi-C scaffolding tool. BIOINFORMATICS 39 (2023).

41. Hassani, Z. S., Salehi-Abargouei, A., Mirzaei, M., Nadjarzadeh, A. & Hosseinzadeh, M., The association between dietary approaches to stop hypertension diet and mediterranean diet with metabolic syndrome in a large sample of Iranian adults: YaHS and TAMYZ Studies. FOOD SCI NUTR 9 3932 (2021).

42. Rhie, A., Walenz, B. P., Koren, S. & Phillippy, A. M., Merqury: reference-free quality, completeness, and phasing assessment for genome assemblies. GENOME BIOL 21 245 (2020).

43. Dierckxsens, N., Mardulyn, P. & Smits, G., NOVOPlasty: de novo assembly of organelle genomes from whole genome data. NUCLEIC ACIDS RES 45 e18 (2017).

44. Flynn, J. M. et al., RepeatModeler2 for automated genomic discovery of transposable element families. P NATL ACAD SCI USA 117 9451 (2020).

45. Benson, G., Tandem repeats finder: a program to analyze DNA sequences. NUCLEIC ACIDS RES 27 573 (1999).

46. Kirov, I., Gilyok, M., Knyazev, A. & Fesenko, I., Pilot satellitome analysis of the model plant, Physcomitrellapatens, revealed a transcribed and high-copy IGS related tandem repeat. COMP CYTOGENET 12 493 (2018).

47. Xu, Z. & Wang, H., LTR_FINDER: an efficient tool for the prediction of full-length LTR retrotransposons. NUCLEIC ACIDS RES 35 W265 (2007).

48. Gremme, G., Brendel, V., Sparks, M. E. & Kurtz, S., Engineering a software tool for gene structure prediction in higher organisms. INFORM SOFTWARE TECH 47 965 (2005).

49. Kim, D., Paggi, J. M., Park, C., Bennett, C. & Salzberg, S. L., Graph-based genome alignment and genotyping with HISAT2 and HISAT-genotype. NAT BIOTECHNOL 37 907 (2019).

50. Shumate, A., Wong, B., Pertea, G. & Pertea, M., Improved transcriptome assembly using a hybrid of long and short reads with StringTie. PLOS COMPUT BIOL 18 e1009730 (2022).

51. Bruna, T., Hoff, K. J., Lomsadze, A., Stanke, M. & Borodovsky, M., BRAKER2: automatic eukaryotic genome annotation with GeneMark-EP+ and AUGUSTUS supported by a protein database. NAR GENOM BIOINFORM 3 lqaa108 (2021).

52. Stanke, M., Diekhans, M., Baertsch, R. & Haussler, D., Using native and syntenically mapped cDNA alignments to improve de novo gene finding. BIOINFORMATICS 24 637 (2008).

53. Bruna, T., Lomsadze, A. & Borodovsky, M., GeneMark-EP+: eukaryotic gene prediction with self-training in the space of genes and proteins. NAR GENOM BIOINFORM 2 lqaa026 (2020).

54. Haas, B. J. et al., Automated eukaryotic gene structure annotation using EVidenceModeler and the Program to Assemble Spliced Alignments. GENOME BIOL 9 R7 (2008).

55. Rhind, N. et al., Comparative functional genomics of the fission yeasts. SCIENCE 332 930 (2011).

56. Cantalapiedra, C. P., Hernandez-Plaza, A., Letunic, I., Bork, P. & Huerta-Cepas, J., eggNOG-mapper v2: Functional Annotation, Orthology Assignments, and Domain Prediction at the Metagenomic Scale. MOL BIOL EVOL 38 5825 (2021).

57. Huerta-Cepas, J. et al., eggNOG 5.0: a hierarchical, functionally and phylogenetically annotated orthology resource based on 5090 organisms and 2502 viruses. NUCLEIC ACIDS RES 47 D309 (2019).

58. Bairoch, A. & Apweiler, R., The SWISS-PROT protein sequence database and its supplement TrEMBL in 2000. NUCLEIC ACIDS RES 28 45 (2000).

59. O’Leary, N. A. et al., Reference sequence (RefSeq) database at NCBI: current status, taxonomic expansion, and functional annotation. NUCLEIC ACIDS RES 44 D733 (2016).

60. El-Gebali, S. et al., The Pfam protein families database in 2019. NUCLEIC ACIDS RES 47 D427 (2019).

61. Kanehisa, M. & Goto, S., KEGG: kyoto encyclopedia of genes and genomes. NUCLEIC ACIDS RES 28 27 (2000).

62. Lowe, T. M. & Eddy, S. R., tRNAscan-SE: a program for improved detection of transfer RNA genes in genomic sequence. NUCLEIC ACIDS RES 25 955 (1997).

63. Nawrocki, E. P. & Eddy, S. R., Infernal 1.1: 100-fold faster RNA homology searches. BIOINFORMATICS 29 2933 (2013).

64. Griffiths-Jones, S. et al., Rfam: annotating non-coding RNAs in complete genomes. NUCLEIC ACIDS RES 33 D121 (2005).

65. Jung, J., Kim, J. I., Jeong, Y. S. & Yi, G., AGORA: organellar genome annotation from the amino acid and nucleotide references. BIOINFORMATICS 34 2661 (2018).

66. Riva, G. & Mauri, M., MuMMER: How Robotics Can Reboot Social Interaction and Customer Engagement in Shops and Malls. CYBERPSYCH BEH SOC N 24 210 (2021).

67. Wang, Y. et al., MCScanX: a toolkit for detection and evolutionary analysis of gene synteny and collinearity. NUCLEIC ACIDS RES 40 e49 (2012).

68. He, W. et al., NGenomeSyn: an easy-to-use and flexible tool for publication-ready visualization of syntenic relationships across multiple genomes. BIOINFORMATICS 39 (2023).

69. Emms, D. M. & Kelly, S., OrthoFinder: phylogenetic orthology inference for comparative genomics. GENOME BIOL 20 238 (2019).

70. Suyama, M., Torrents, D. & Bork, P., PAL2NAL: robust conversion of protein sequence alignments into the corresponding codon alignments. NUCLEIC ACIDS RES 34 W609 (2006).

71. Stamatakis, A., RAxML version 8: a tool for phylogenetic analysis and post-analysis of large phylogenies. BIOINFORMATICS 30 1312 (2014).

72. Kozlov, A. M., Aberer, A. J. & Stamatakis, A., ExaML version 3: a tool for phylogenomic analyses on supercomputers. BIOINFORMATICS 31 2577 (2015).

73. Zhang, C., Rabiee, M., Sayyari, E. & Mirarab, S., ASTRAL-III: polynomial time species tree reconstruction from partially resolved gene trees. BMC BIOINFORMATICS 19 153 (2018).

74. Kumar, S. et al., TimeTree 5: An Expanded Resource for Species Divergence Times. MOL BIOL EVOL 39 (2022).

75. Yang, Z., PAML 4: phylogenetic analysis by maximum likelihood. MOL BIOL EVOL 24 1586 (2007).

76. Armstrong, J. et al., Progressive Cactus is a multiple-genome aligner for the thousand-genome era. NATURE 587 246 (2020).

77. Wright, A. E. et al., Variation in promiscuity and sexual selection drives avian rate of Faster-Z evolution. MOL ECOL 24 1218 (2015).

78. Jaiswal, S. K. et al., Genome Sequence of Peacock Reveals the Peculiar Case of a Glittering Bird. FRONT GENET 9 392 (2018).

79. Li, H. & Durbin, R., Inference of human population history from individual whole-genome sequences. NATURE 475 493 (2011).

80. Nadachowska-Brzyska, K., Li, C., Smeds, L., Zhang, G. & Ellegren, H., Temporal Dynamics of Avian Populations during Pleistocene Revealed by Whole-Genome Sequences. CURR BIOL 25 1375 (2015).

81. Terhorst, J., Kamm, J. A. & Song, Y. S., Robust and scalable inference of population history from hundreds of unphased whole genomes. NAT GENET 49 303 (2017).

82. You, M. et al., Variation among 532 genomes unveils the origin and evolutionary history of a global insect herbivore. NAT COMMUN 11 2321 (2020).

83. Gutenkunst, R. N., Hernandez, R. D., Williamson, S. H. & Bustamante, C. D., Inferring the joint demographic history of multiple populations from multidimensional SNP frequency data. PLOS GENET 5 e1000695 (2009).

84. Huang, X. et al., Inferring Genome-Wide Correlations of Mutation Fitness Effects between Populations. MOL BIOL EVOL 38 4588 (2021).

85. Wen, J. et al., Origins, timing and introgression of domestic geese revealed by whole genome data. J ANIM SCI BIOTECHNO 14 26 (2023).

86. Coffman, A. J., Hsieh, P. H., Gravel, S. & Gutenkunst, R. N., Computationally Efficient Composite Likelihood Statistics for Demographic Inference. MOL BIOL EVOL 33 591 (2016).

87. Mendes, F. K., Vanderpool, D., Fulton, B. & Hahn, M. W., CAFE 5 models variation in evolutionary rates among gene families. BIOINFORMATICS 36 5516 (2021).

88. Bu, D. et al., KOBAS-i: intelligent prioritization and exploratory visualization of biological functions for gene enrichment analysis. NUCLEIC ACIDS RES 49 W317 (2021).

89. Chen, S., Zhou, Y., Chen, Y. & Gu, J., fastp: an ultra-fast all-in-one FASTQ preprocessor. BIOINFORMATICS 34 i884 (2018).

90. Li, H. & Durbin, R., Fast and accurate long-read alignment with Burrows-Wheeler transform. BIOINFORMATICS 26 589 (2010).

91. Danecek, P. et al., Twelve years of SAMtools and BCFtools. GIGASCIENCE 10 (2021).

92. Okonechnikov, K., Conesa, A. & Garcia-Alcalde, F., Qualimap 2: advanced multi-sample quality control for high-throughput sequencing data. BIOINFORMATICS 32 292 (2016).

93. McKenna, A. et al., The Genome Analysis Toolkit: a MapReduce framework for analyzing next-generation DNA sequencing data. GENOME RES 20 1297 (2010).

94. Cingolani, P. et al., Using Drosophila melanogaster as a Model for Genotoxic Chemical Mutational Studies with a New Program, SnpSift. FRONT GENET 3 35 (2012).

95. Wang, X. et al., CNVcaller: highly efficient and widely applicable software for detecting copy number variations in large populations. GIGASCIENCE 6 1 (2017).

96. Purcell, S. et al., PLINK: a tool set for whole-genome association and population-based linkage analyses. AM J HUM GENET 81 559 (2007).

97. Alexander, D. H. & Lange, K., Enhancements to the ADMIXTURE algorithm for individual ancestry estimation. BMC BIOINFORMATICS 12 246 (2011).

98. Tamura, K., Stecher, G. & Kumar, S., MEGA11: Molecular Evolutionary Genetics Analysis Version 11. MOL BIOL EVOL 38 3022 (2021).

99. Pickrell, J. K. & Pritchard, J. K., Inference of population splits and mixtures from genome-wide allele frequency data. PLOS GENET 8 e1002967 (2012).

100. Danecek, P. et al., The variant call format and VCFtools. BIOINFORMATICS 27 2156 (2011).

101. Zhang, C., Dong, S. S., Xu, J. Y., He, W. M. & Yang, T. L., PopLDdecay: a fast and effective tool for linkage disequilibrium decay analysis based on variant call format files. BIOINFORMATICS 35 1786 (2019).

102. Curik, I., Ferenčaković, M. & Sölkner, J., Inbreeding and runs of homozygosity: A possible solution to an old problem. LIVEST SCI 166 26 (2014).

103. Darriba, D., Taboada, G. L., Doallo, R. & Posada, D., jModelTest 2: more models, new heuristics and parallel computing. NAT METHODS 9 772 (2012).

104. Rozas, J. et al., DnaSP 6: DNA Sequence Polymorphism Analysis of Large Data Sets. MOL BIOL EVOL 34 3299 (2017).

105. Leigh, J. W. & Bryant, D., <scp>popart<scp>: full-feature software for haplotype network construction. METHODS ECOL EVOL 6 1110 (2015).

106. Dias-Alves, T., Mairal, J. & Blum, M., Loter: A Software Package to Infer Local Ancestry for a Wide Range of Species. MOL BIOL EVOL 35 2318 (2018).

107. Pickrell, J. K. & Pritchard, J. K., Inference of population splits and mixtures from genome-wide allele frequency data. PLOS GENET 8 e1002967 (2012).

108. Patterson, N. et al., Ancient admixture in human history. GENETICS 192 1065 (2012).

109. Malinsky, M., Matschiner, M. & Svardal, H., Dsuite - Fast D-statistics and related admixture evidence from VCF files. MOL ECOL RESOUR 21 584 (2021).

110. Moran, R. L. et al., Hybridization underlies localized trait evolution in cavefish. ISCIENCE 25 103778 (2022).

111. Pertea, G. & Pertea, M., GFF Utilities: GffRead and GffCompare. F1000Res 9 (2020).

112. Love, M. I., Huber, W. & Anders, S., Moderated estimation of fold change and dispersion for RNA-seq data with DESeq2. GENOME BIOL 15 550 (2014).

113. Saski, M., Ikechi, T. & Makino, S., A feather pulp culture technique for avian chromosomes, with notes on the chromosomes of the peafowl and the ostrich. Experientia 24 1292 (1968).

114. Manni, M., Berkeley, M. R., Seppey, M., Simao, F. A. & Zdobnov, E. M., BUSCO Update: Novel and Streamlined Workflows along with Broader and Deeper Phylogenetic Coverage for Scoring of Eukaryotic, Prokaryotic, and Viral Genomes. MOL BIOL EVOL 38 4647 (2021).

115. Zhang, G. et al., Comparative genomics reveals insights into avian genome evolution and adaptation. SCIENCE 346 1311 (2014).

116. Kapusta, A., Suh, A. & Feschotte, C., Dynamics of genome size evolution in birds and mammals. P NATL ACAD SCI USA 114 E1460 (2017).

117. Ellegren, H., Evolutionary stasis: the stable chromosomes of birds. TRENDS ECOL EVOL 25 283 (2010).

118. Kretschmer, R., Ferguson-Smith, M. A. & de Oliveira, E., Karyotype Evolution in Birds: From Conventional Staining to Chromosome Painting. GENES-BASEL 9 (2018).

119. Kawahara-Miki, R. et al., Next-generation sequencing reveals genomic features in the Japanese quail. GENOMICS 101 345 (2013).

120. He, C. et al., Chromosome level assembly reveals a unique immune gene organization and signatures of evolution in the common pheasant. MOL ECOL RESOUR 21 897 (2021).

121. Suh, A., Smeds, L. & Ellegren, H., The Dynamics of Incomplete Lineage Sorting across the Ancient Adaptive Radiation of Neoavian Birds. PLOS BIOL 13 e1002224 (2015).

122. Jarvis, E. D. et al., Whole-genome analyses resolve early branches in the tree of life of modern birds. SCIENCE 346 1320 (2014).

123. Hyun, S., Body size regulation and insulin-like growth factor signaling. CELL MOL LIFE SCI 70 2351 (2013).

124. Maridas, D. E. et al., IGFBP-4 regulates adult skeletal growth in a sex-specific manner. J ENDOCRINOL 233 131 (2017).

125. McPherron, A. C. & Lee, S. J., Double muscling in cattle due to mutations in the myostatin gene. P NATL ACAD SCI USA 94 12457 (1997).

126. Ekblom, R., French, L., Slate, J. & Burke, T., Evolutionary analysis and expression profiling of zebra finch immune genes. GENOME BIOL EVOL 2 781 (2010).

127. Ramakrishnan, B. et al., A Structural and Mathematical Modeling Analysis of the Likelihood of Antibody-Dependent Enhancement in Influenza. TRENDS MICROBIOL 24 933 (2016).

128. Gui, Y. & Murphy, L. J., Insulin-like growth factor (IGF)-binding protein-3 (IGFBP-3) binds to fibronectin (FN): demonstration of IGF-I/IGFBP-3/fn ternary complexes in human plasma. The Journal of Clinical Endocrinology & Metabolism 86 2104 (2001).

129. Hwa, V., Oh, Y. & Rosenfeld, R. G., The insulin-like growth factor-binding protein (IGFBP) superfamily. ENDOCR REV 20 761 (1999).

130. Prakasam, R. et al., Chicken IL-6 is a heat-shock gene. FEBS LETT 587 3541 (2013).

131. Kaiser, P., Rothwell, L., Goodchild, M. & Bumstead, N., The chicken proinflammatory cytokines interleukin-1β and interleukin-6: differences in gene structure and genetic location compared with their mammalian orthologues. ANIM GENET 35 169 (2004).

132. Zhang, D., Ding, Z. & Xu, X., Pathologic Mechanisms of the Newcastle Disease Virus. VIRUSES-BASEL 15 864 (2023).

133. Darwin, C., The variation of animals and plants under domestication. (J. Murray, 1868).

134. Kawasaki-Nishihara, A., Nishihara, D., Nakamura, H. & Yamamoto, H., ET3/Ednrb2 signaling is critically involved in regulating melanophore migration in Xenopus. DEV DYNAM 240 1454 (2011).

135. Lecoin, L. et al., Cloning and characterization of a novel endothelin receptor subtype in the avian class. P NATL ACAD SCI USA 95 3024 (1998).

136. Lahav, R., Ziller, C., Dupin, E. & Le Douarin, N. M., Endothelin 3 promotes neural crest cell proliferation and mediates a vast increase in melanocyte number in culture. P NATL ACAD SCI USA 93 3892 (1996).

137. Kurosaki, T., Popp, M. W. & Maquat, L. E., Quality and quantity control of gene expression by nonsense-mediated mRNA decay. NAT REV MOL CELL BIO 20 406 (2019).

138. Hu, C. et al., CellMarker 2.0: an updated database of manually curated cell markers in human/mouse and web tools based on scRNA-seq data. NUCLEIC ACIDS RES 51 D870 (2023).

139. Zhang, Z. et al., Whole-genome resequencing reveals signatures of selection and timing of duck domestication. GIGASCIENCE 7 (2018).

140. Kerje, S. et al., The Dominant white, Dun and Smoky color variants in chicken are associated with insertion/deletion polymorphisms in the PMEL17 gene. GENETICS 168 1507 (2004).

141. Rubio, A. O. & Summers, K., Neural crest cell genes and the domestication syndrome: A comparative analysis of selection. PLOS ONE 17 e0263830 (2022).

142. Zhou, Z. et al., An intercross population study reveals genes associated with body size and plumage color in ducks. NAT COMMUN 9 2648 (2018).

143. Zhang, Z. et al., Whole-genome resequencing reveals signatures of selection and timing of duck domestication. GIGASCIENCE 7 giy027 (2018).

144. Kawagoshi, T. et al., Molecular structures of centromeric heterochromatin and karyotypic evolution in the Siamese crocodile (Crocodylus siamensis) (Crocodylidae, Crocodylia). CHROMOSOME RES 16 1119 (2008).

145. Lam, E. T. et al., Genome mapping on nanochannel arrays for structural variation analysis and sequence assembly. NAT BIOTECHNOL 30 771 (2012).

146. Zheng, G. X. et al., Haplotyping germline and cancer genomes with high-throughput linked-read sequencing. NAT BIOTECHNOL 34 303 (2016).

147. Zhang, G. et al., Comparative genomics reveals insights into avian genome evolution and adaptation. SCIENCE 346 1311 (2014).

148. Barros, C. P. et al., A new haplotype-resolved turkey genome to enable turkey genetics and genomics research. bioRxiv 2022 (2022).

149. Kaiser, V. B., van Tuinen, M. & Ellegren, H., Insertion events of CR1 retrotransposable elements elucidate the phylogenetic branching order in galliform birds. MOL BIOL EVOL 24 338 (2007).

150. Wang, N., Kimball, R. T., Braun, E. L., Liang, B. & Zhang, Z., Ancestral range reconstruction of Galliformes: the effects of topology and taxon sampling. J BIOGEOGR 44 122 (2017).

151. Nadachowska-Brzyska, K., Li, C., Smeds, L., Zhang, G. & Ellegren, H., Temporal dynamics of avian populations during Pleistocene revealed by whole-genome sequences. CURR BIOL 25 1375 (2015).

152. Hung, C. et al., Drastic population fluctuations explain the rapid extinction of the passenger pigeon. Proceedings of the National Academy of Sciences 111 10636 (2014).

153. Jaiswal, S. K. et al., Genome sequence of peacock reveals the peculiar case of a glittering bird. FRONT GENET 9 392 (2018).

154. Prüfer, K. et al., The complete genome sequence of a Neanderthal from the Altai Mountains. NATURE 505 43 (2014).

155. Zhao, Y. et al., Resequencing 545 ginkgo genomes across the world reveals the evolutionary history of the living fossil. NAT COMMUN 10 4201 (2019).

156. Shi, S. et al., Whole genome analyses reveal novel genes associated with chicken adaptation to tropical and frigid environments. J ADV RES 47 13 (2023).

157. Wilson, M. C. et al., (Springer, 2016), Vol. 31, pp. 219.

158. Bai, Y. et al., New ecological redline policy (ERP) to secure ecosystem services in China. LAND USE POLICY 55 348 (2016).

159. Wu, F. et al., Ongoing green peafowl protection in China. ZOOL RES 40 580 (2019).

160. Du, H. Y. et al., Identification of hybrid green peafowl using mitochondrial and nuclear markers. CONSERV GENET RESOUR 12 669 (2020).

161. Goes, F., The status and distribution of green peafowl Pavo muticus in Cambodia. Cambodian Journal of Natural History 2009 7 (2009).

162. Vellend, M. et al., Effects of exotic species on evolutionary diversification. TRENDS ECOL EVOL 22 481 (2007).

163. Yamada, P. M. & Lee, K., Perspectives in mammalian IGFBP-3 biology: local vs. systemic action. AM J PHYSIOL-CELL PH 296 C954 (2009).

164. Cong, R. et al., Maternal high-protein diet modulates hepatic growth axis in weaning piglets by reprogramming the IGFBP-3 gene. EUR J NUTR 59 2497 (2020).

165. Schlee, P. et al., Growth hormone and insulin-like growth factor I concentrations in bulls of various growth hormone genotypes. THEOR APPL GENET 88 497 (1994).

166. Shen, M. et al., A novel polymorphism of IGFBP-3 gene and its relationship with several wool traits in Chinese Merino sheep. Yi Chuan= Hereditas 30 1182 (2008).

167. Guo, Y. et al., Evolutionary analysis and functional characterization reveal the role of the insulin-like growth factor system in a diversified selection of chickens (Gallus gallus). Poultry Science 102 102411 (2023).

168. Talha, M. M. H. et al., Morphometric, productive and reproductive traits of Indian peafowl (Pavo cristatus) in Bangladesh. International Journal of Development Research 8 19039 (2018).

169. Lawal, R. A. et al., The wild species genome ancestry of domestic chickens. BMC BIOL 18 1 (2020).

170. Chen, N. et al., Whole-genome resequencing reveals world-wide ancestry and adaptive introgression events of domesticated cattle in East Asia. NAT COMMUN 9 2337 (2018).

171. Gompel, N. & Prud’Homme, B., The causes of repeated genetic evolution. DEV BIOL 332 36 (2009).

172. Christin, P. A., Weinreich, D. M. & Besnard, G., Causes and evolutionary significance of genetic convergence. TRENDS GENET 26 400 (2010).

173. Manceau, M., Domingues, V. S., Linnen, C. R., Rosenblum, E. B. & Hoekstra, H. E., Convergence in pigmentation at multiple levels: mutations, genes and function. PHILOS T R SOC B 365 2439 (2010).

174. Guernsey, M. W. et al., A Val85Met mutation in melanocortin-1 receptor is associated with reductions in eumelanic pigmentation and cell surface expression in domestic rock pigeons (Columba livia). PLOS ONE 8 e74475 (2013).

175. Wang, C. et al., Donkey genomes provide new insights into domestication and selection for coat color. NAT COMMUN 11 6014 (2020).

176. Imsland, F. et al., Regulatory mutations in TBX3 disrupt asymmetric hair pigmentation that underlies Dun camouflage color in horses. NAT GENET 48 152 (2016).

177. Lopes, R. J. et al., Genetic Basis for Red Coloration in Birds. CURR BIOL 26 1427 (2016).

178. Mundy, N. I. et al., Red Carotenoid Coloration in the Zebra Finch Is Controlled by a Cytochrome P450 Gene Cluster. CURR BIOL 26 1435 (2016).

179. Vickrey, A. I., Domyan, E. T., Horvath, M. P. & Shapiro, M. D., Convergent Evolution of Head Crests in Two Domesticated Columbids Is Associated with Different Missense Mutations in EphB2. MOL BIOL EVOL 32 2657 (2015).

