## Supplementary Fig. S1-24 and Supplementary Table. S1-5, 17 for "Genomic evidence for hybridization and introgression between blue peafowl and green peafowl and selection for white plumage"

### Additional Files

This document contains Supplementary Fig. S1-24 and Supplementary Table. S1-5, 17. Please refer to the Supplementary table document for Supplementary Table. S6-16.

[Supplementary Fig. S23.	2](#_Toc143526300)4

[Supplementary Fig. S24.	2](#_Toc143526300)5

[Supplementary Table S1.	2](#_Toc143526301)6

[Supplementary Table S2.	2](#_Toc143526302)7

[Supplementary Table S3.	2](#_Toc143526303)8

[Supplementary Table S4.	2](#_Toc143526304)9

[Supplementary Table S5.](#_Toc143526305)	30

[Supplementary Table S17.](#_Toc143526305)	31

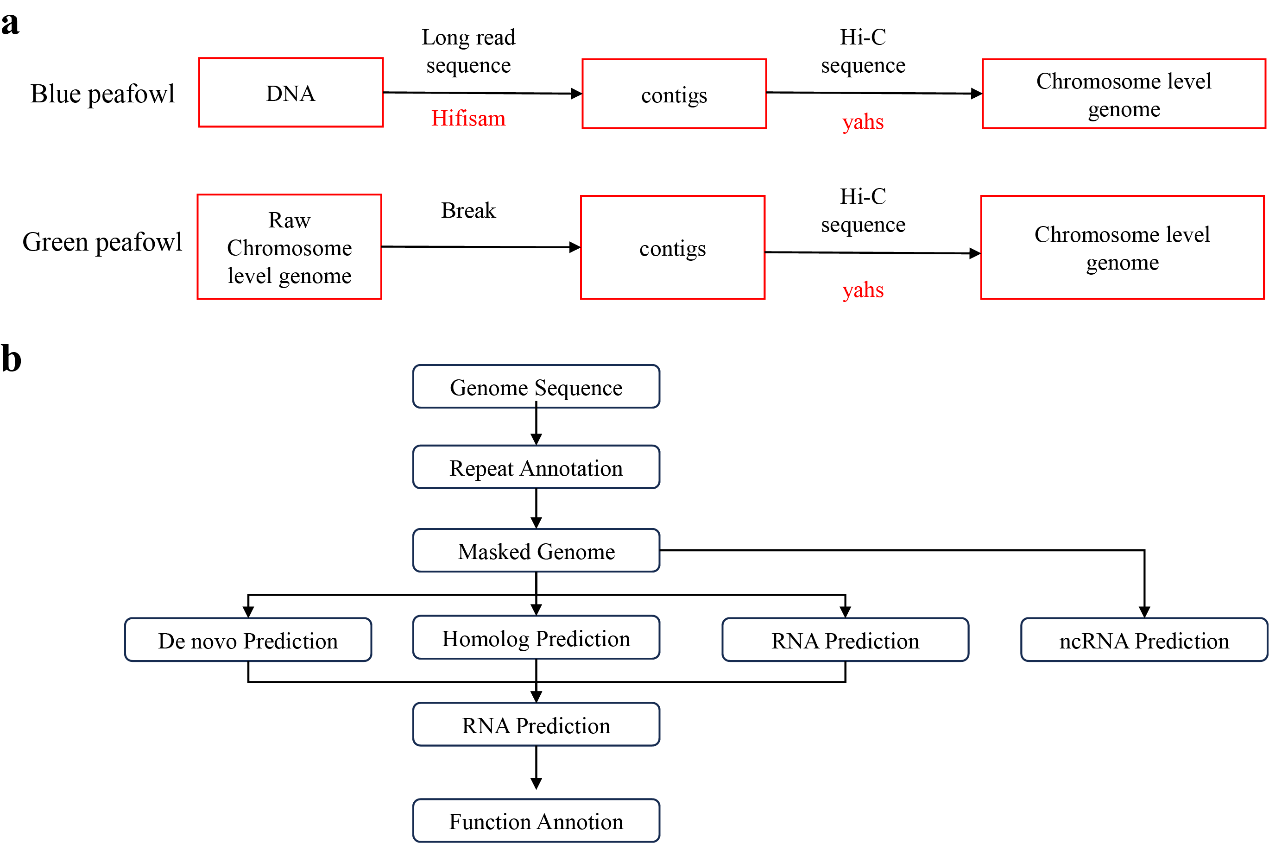

Supplementary Fig. S1. Pipeline of the draft genome assembly and genome annotation of blue peafowl (WP-1). (a) Workflow of the genome assembly, software used is marked in red font; (b) Workflow of the genome annotation.

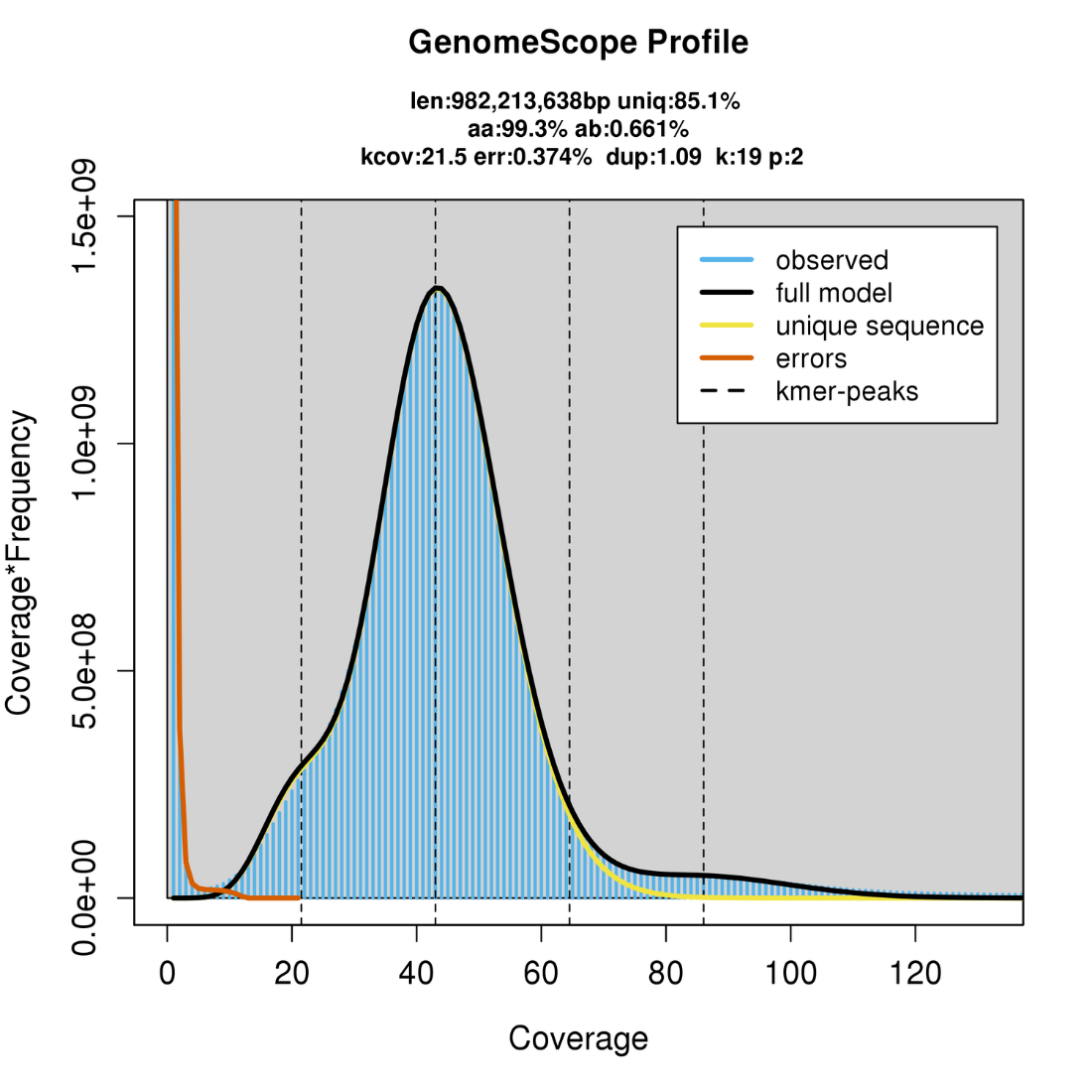

Supplementary Fig. S2. Estimation of the genome size of the blue peafowl by K-mer analysis. As can be seen from the plot, when diploid is used as the fitting standard, the K-mer depth corresponding to the first peak is 21.5, and the calculated length of a single genome is about 982.21 Mb. According to the K-mer distribution, the estimated repetitive sequence content is about 14.85%, and the estimated heterozygosity is about 0.66%.

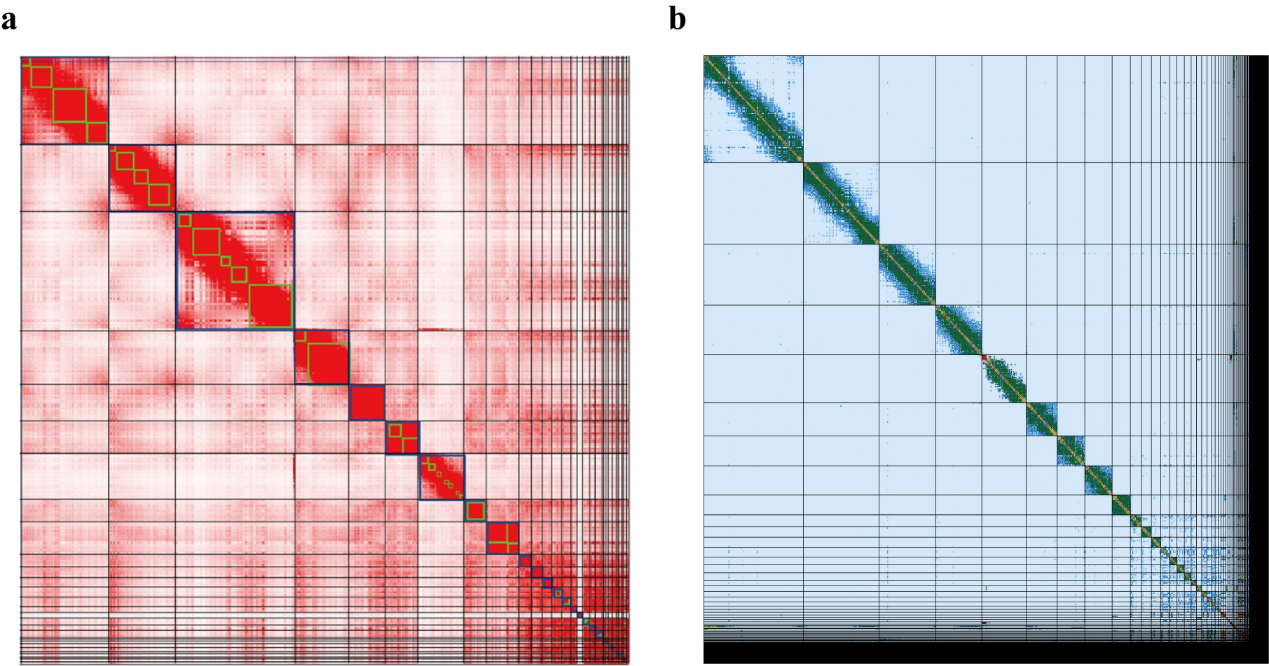

Supplementary Fig. S3. Hi-C-based chromosome-level assemblies of blue peafowl genome and green peafowl. (a) The Hi-C read pairs were mapped to final blue peafowl genome (WP-1) assembly using the yahs pipeline, followed by visualization in Juicebox. (b) The Hi-C read pairs were mapped to final green peafowl genome assembly using the yahs pipeline.

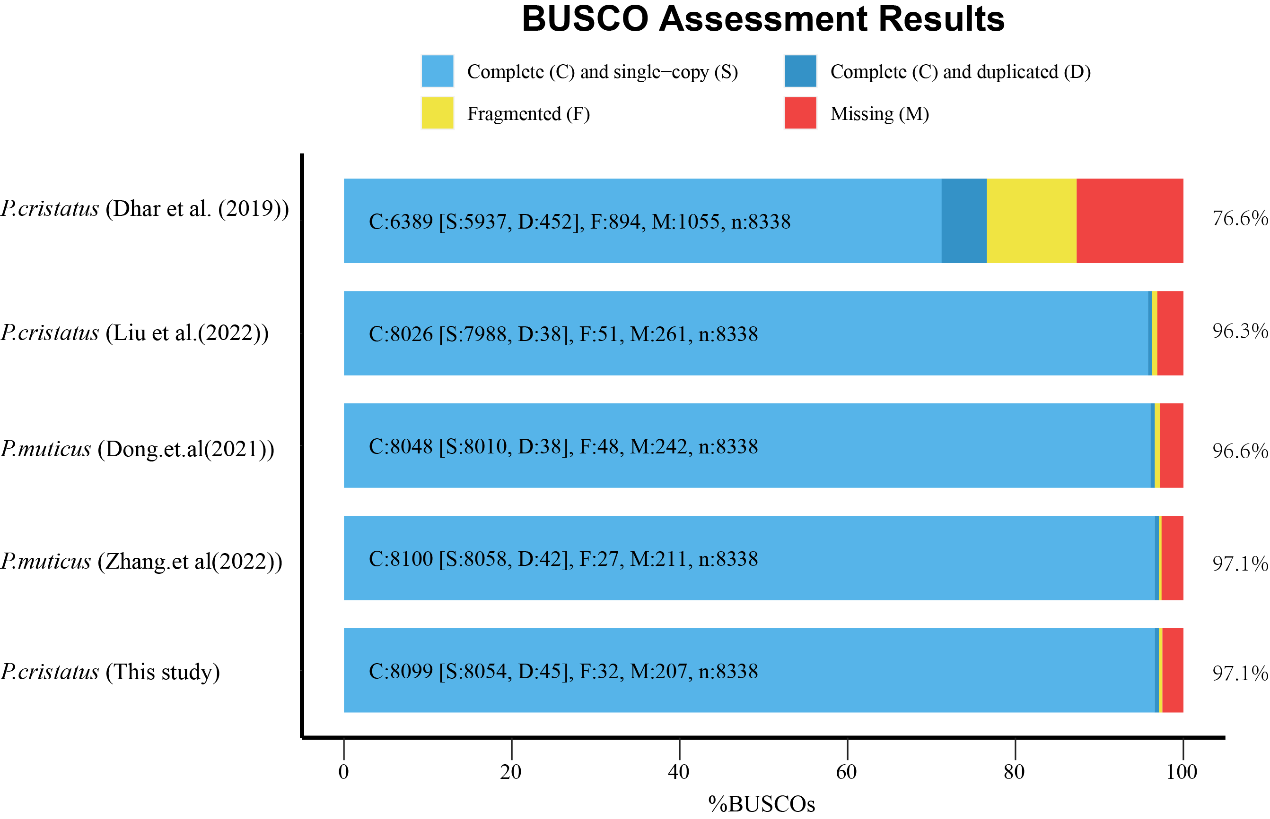

Supplementary Fig. S4. BUSCO assesses the completeness of published genomes and this assembled genome.

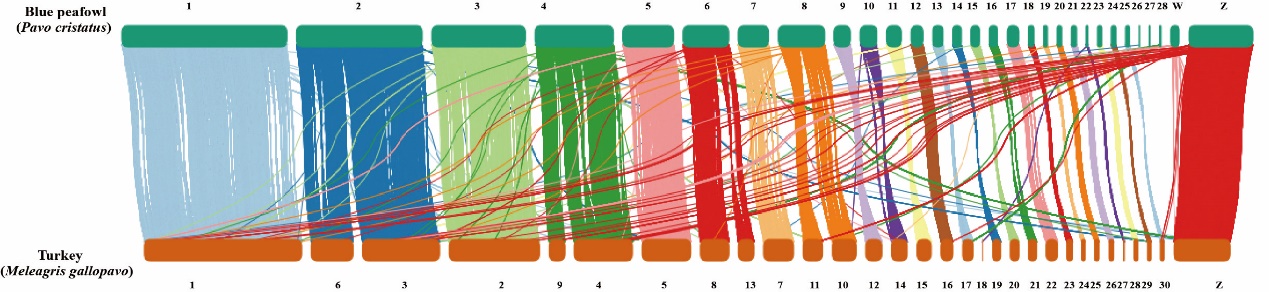

Supplementary Fig. S5. Genome synteny and collinearity among the blue peafowl and Turkey.

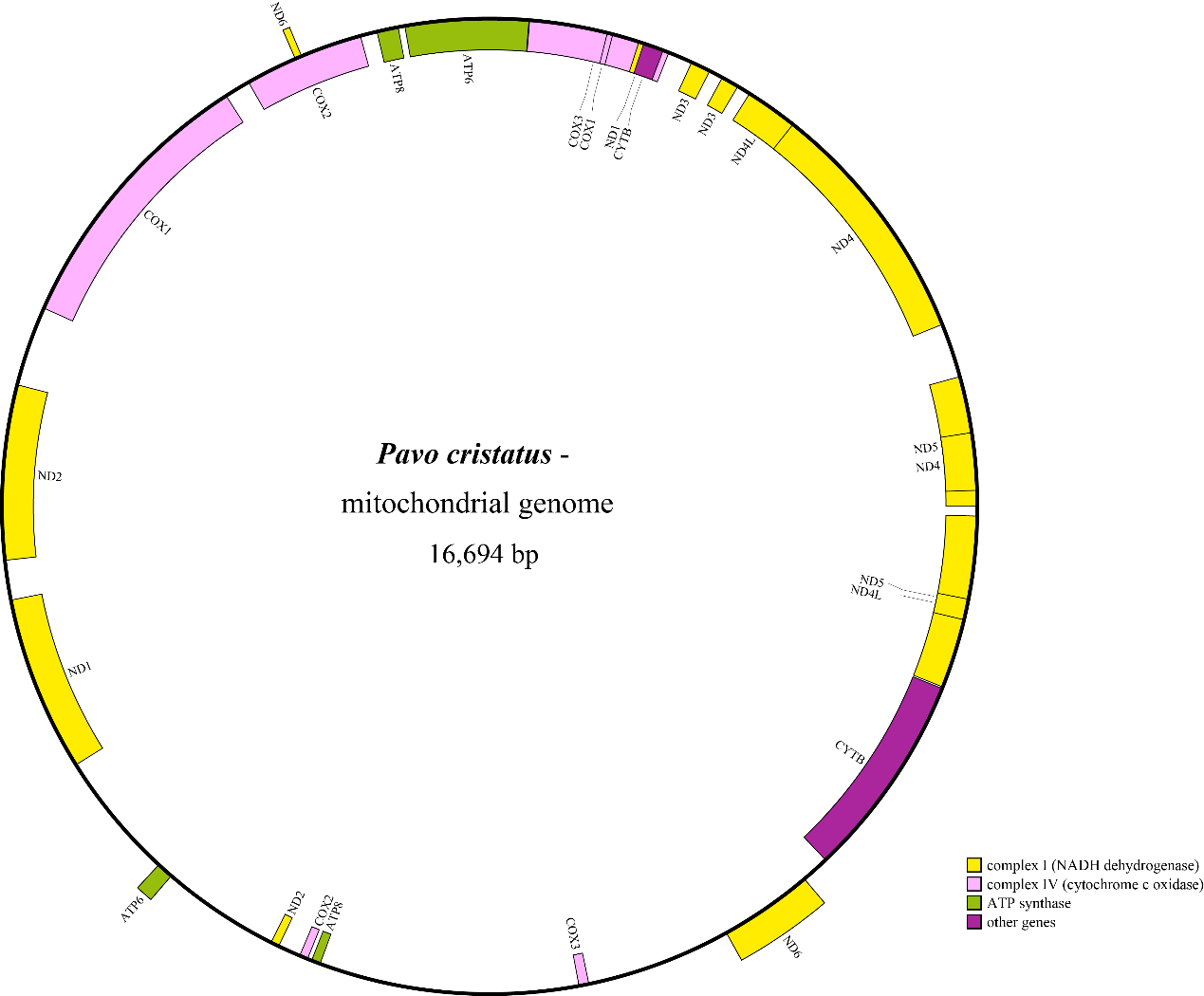

Supplementary Fig. S6. Mitochondrial genome map of blue peafowl. Genes that belong to different functional groups are color-coded.

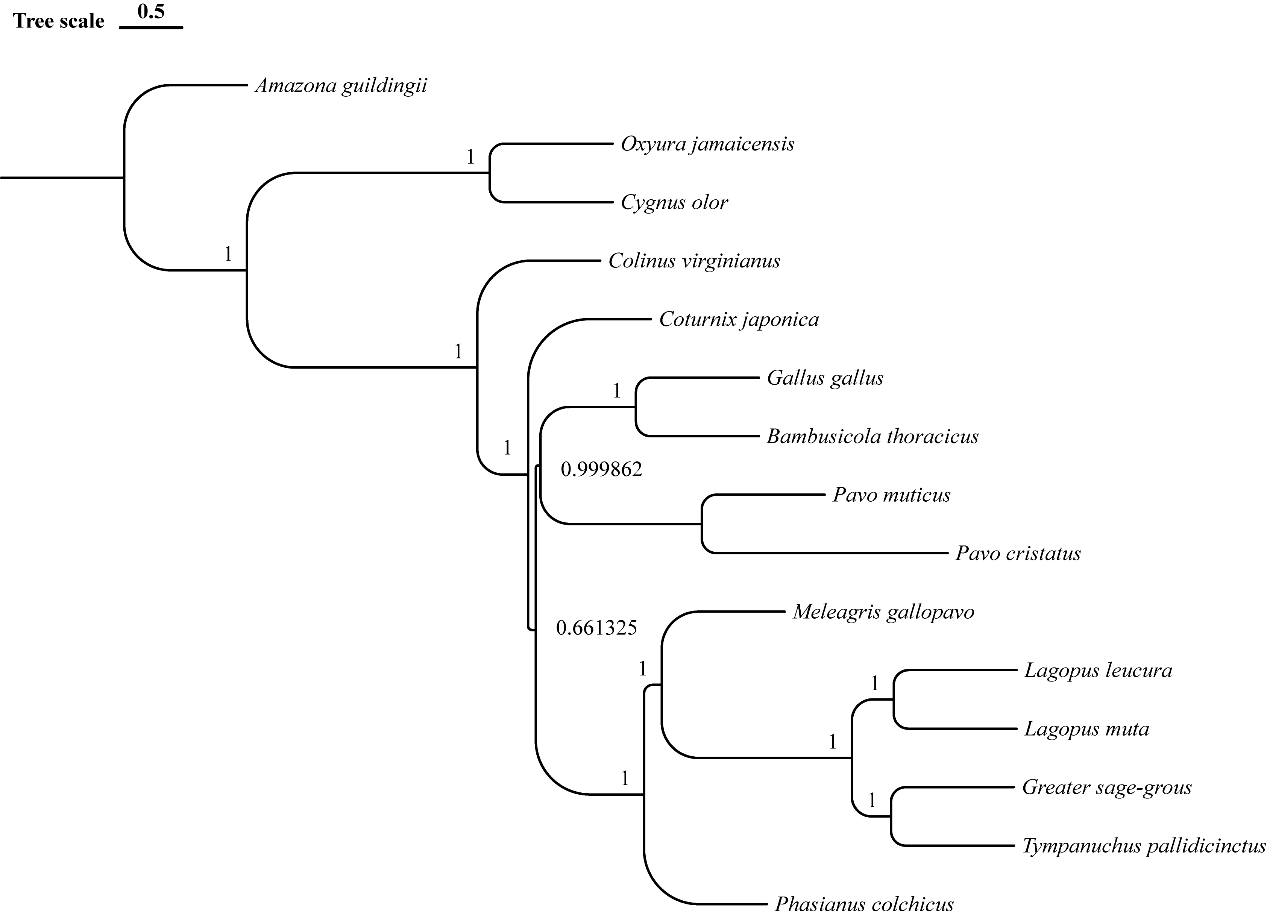

Supplementary Fig. S7. Coalescent species tree inferred from single-copy orthologous gene trees by ASTRAL. A total of 2,323 single-copy orthologous genes trees were used to infer the species tree using ASTRALIII. The local posterior probability is labeled on each node.

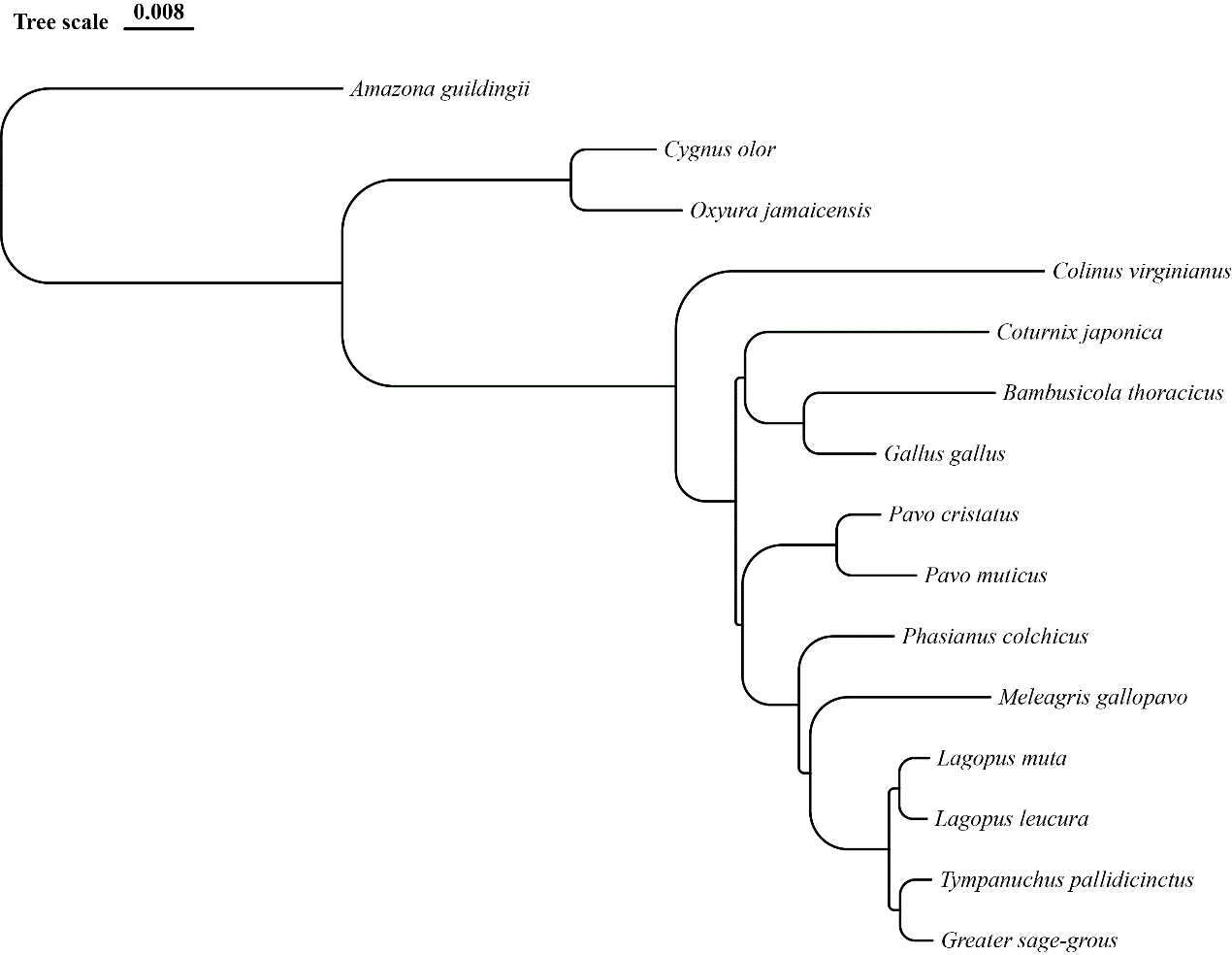

Supplementary Fig. S8. ML phylogenetic tree of 15 birds inferred using single-copy orthologous genes by using iqtree.

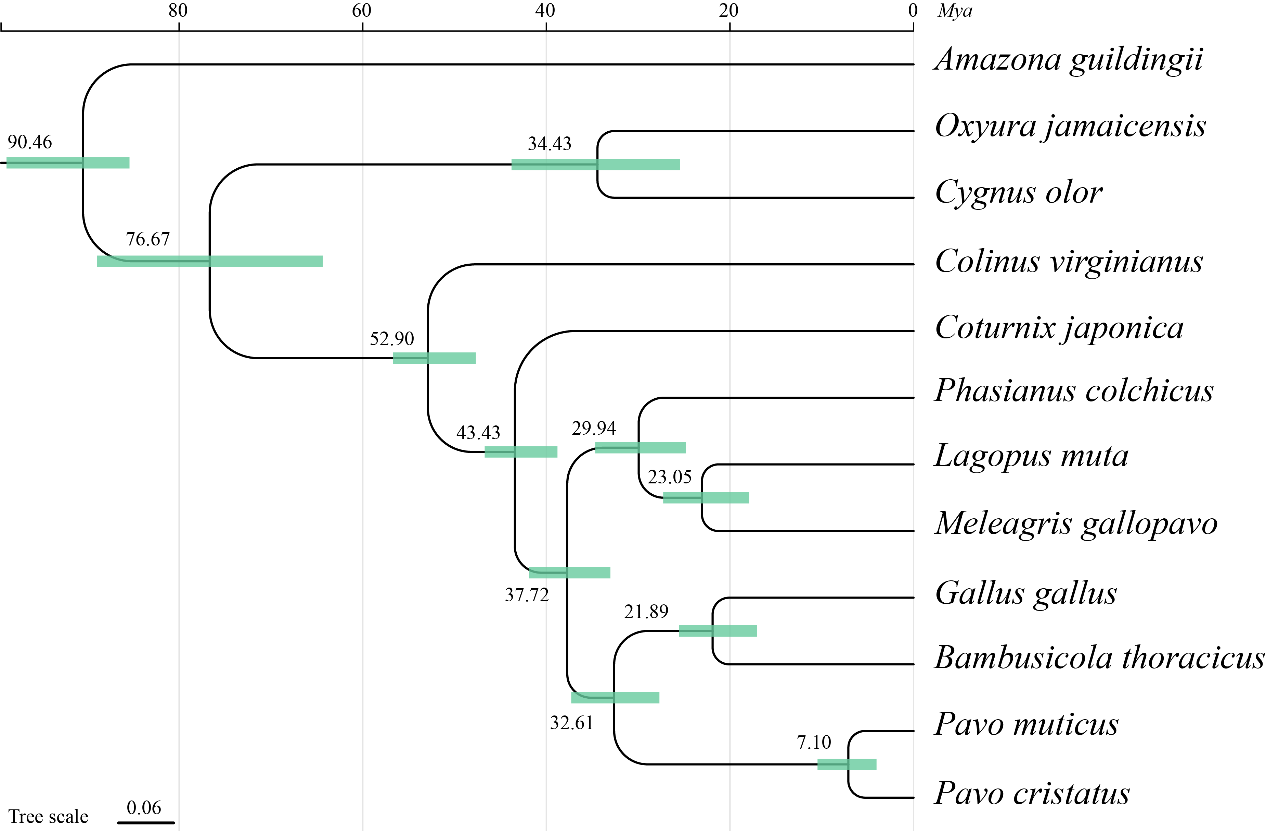

Supplementary Fig. S9. ML phylogenetic tree and molecular clock dating analysis of 11 birds based on mitochondria genomes. Mitochondrial genomes of 11 birds were used to infer a Maximum Likelihood (ML) phylogenetic tree by RAxML and MCMCtree.

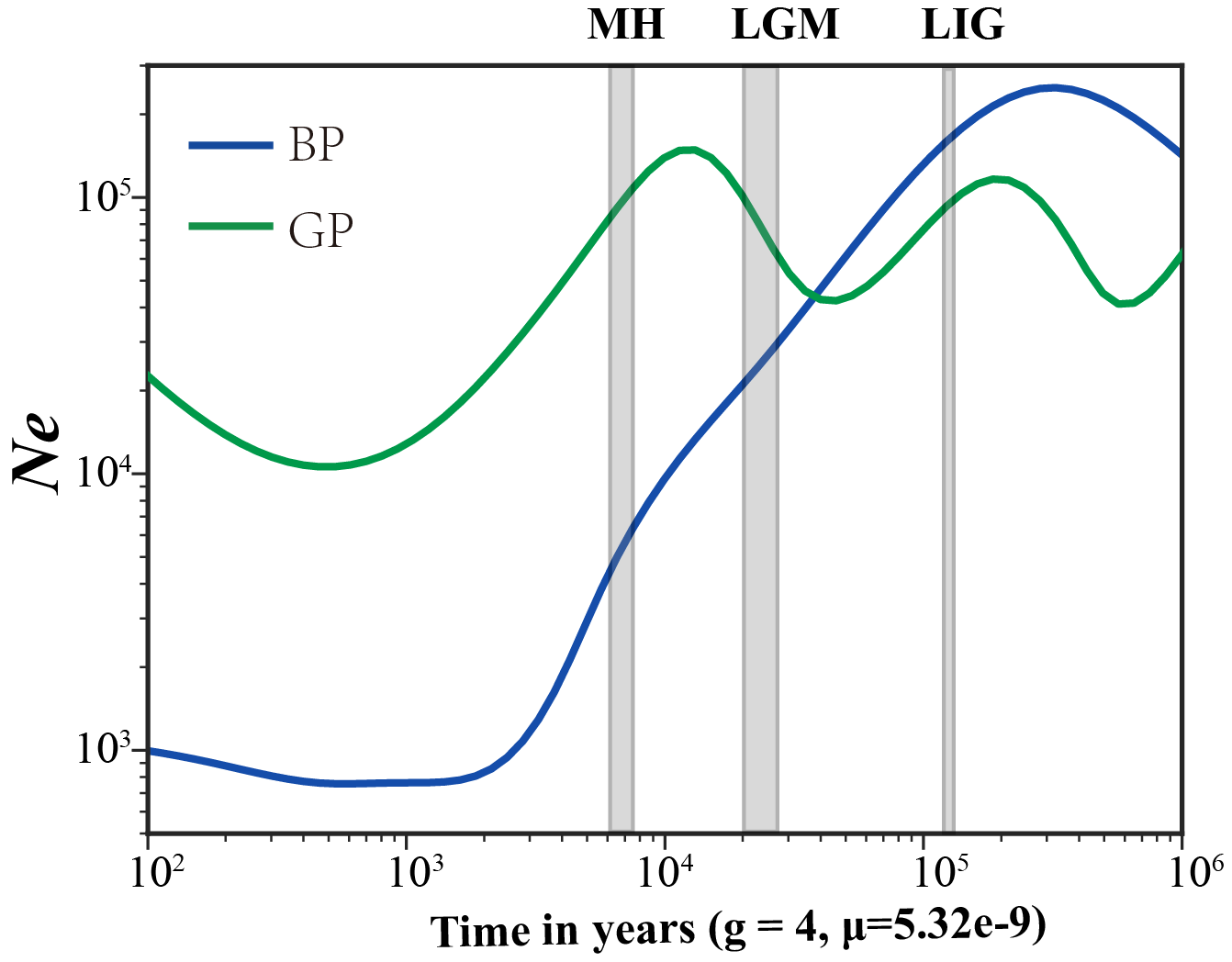

Supplementary Fig. S10. Population size history inference of blue peafowl (BP) and green peafowl (GP). The last interglacial period (LIG, approx. 132–112 Ka), the last glacial maximum (LGM, approx. 26–19 Ka) and the mid-Holocene (MH, 7–5 thousand years ago (Ka)).

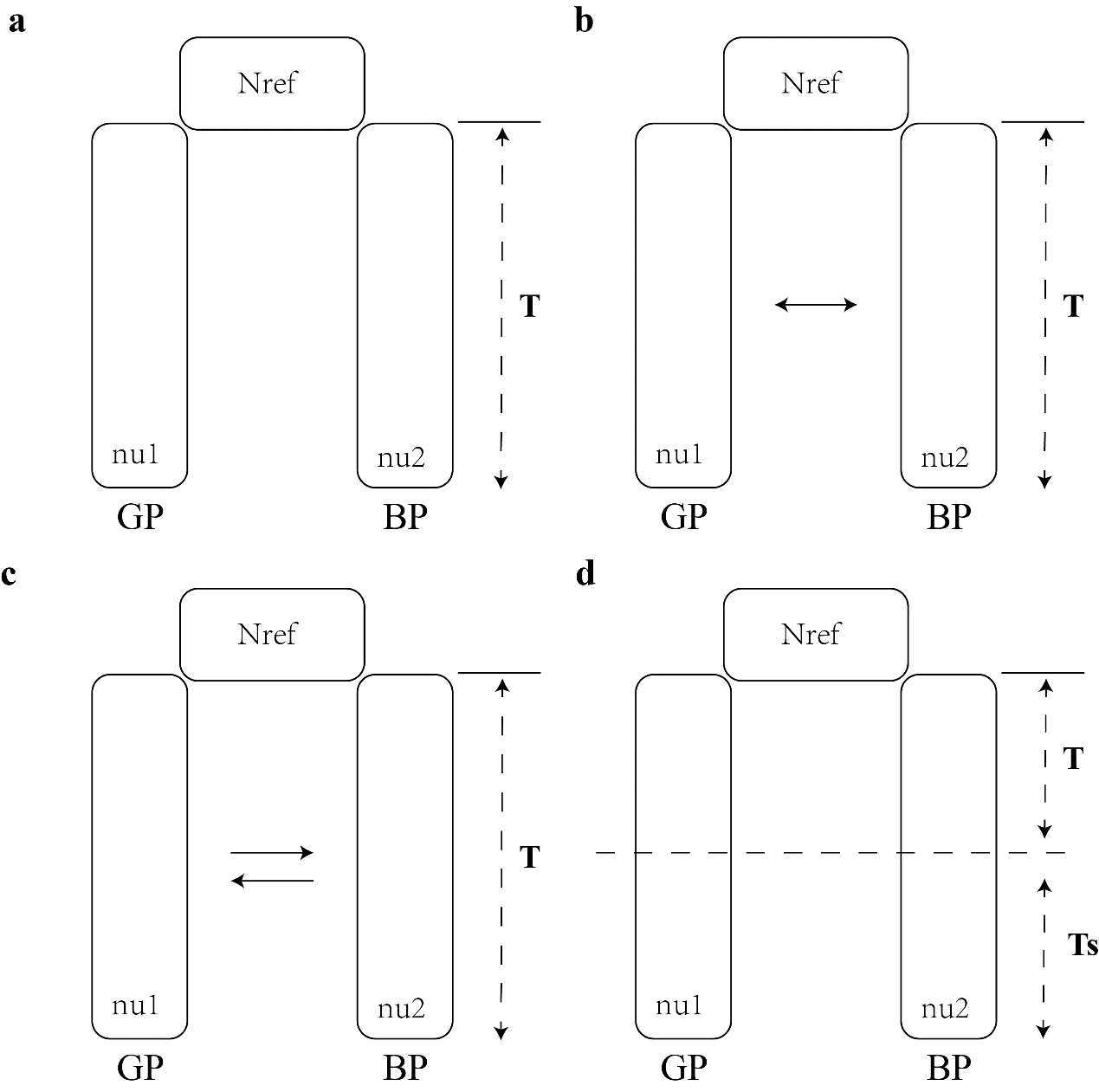

Supplementary Fig. S11. Group history dynamic simulation of blue peafowl and green peafowl by using DaDi. (a) Model 1 (sym_mig): Instantanous size change followed by exponential growth with no population split; (b) Model 2 (bottlegrowth_2d): Instantanous size change followed by exponential growth then split with migration; (c) Model 3 (bottlegrowth_split_mig): Split into two populations of specifed size, with migration; (d) Model 4 (split_asym_mig): Split into two populations of specifed size, with asymetric migration. The model 4 is the best model.

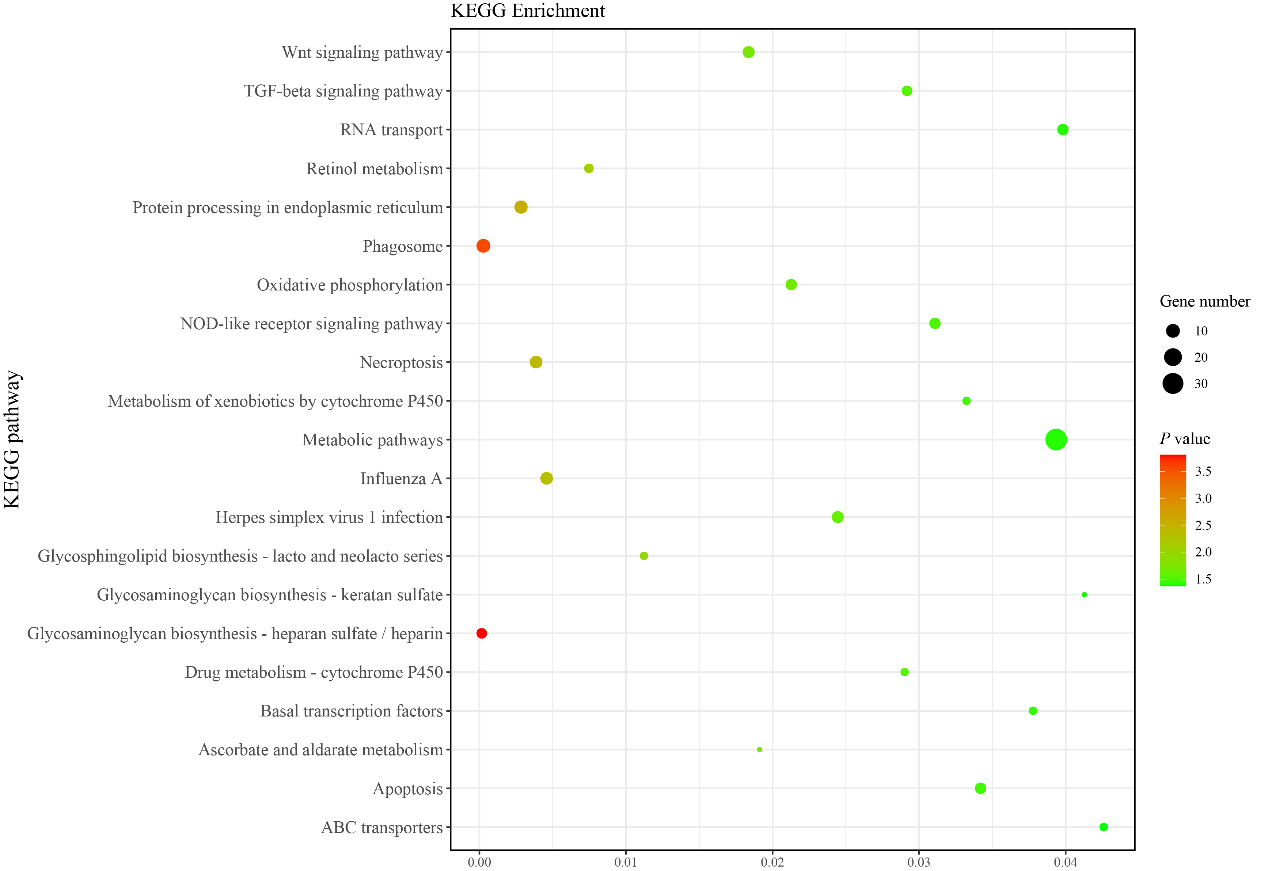

Supplementary Fig. S12. KEGG pathway enrichment analysis of the expansion gene family of peafowl. KEGG pathways with *P*-value < 0.05 were shown.

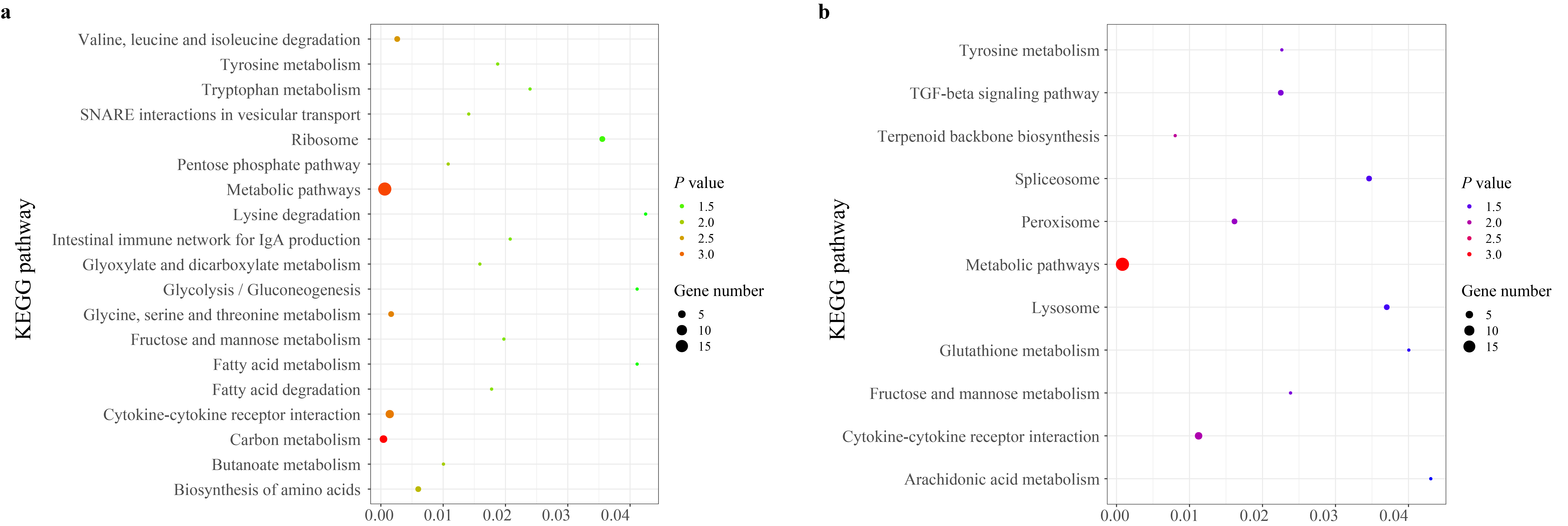

Supplementary Fig. S13. KEGG pathway enrichment analysis of PSG and RGs of peafowl. KEGG pathways with P-value < 0.05 were shown. (a) KEGG pathway enrichment analysis of blue peafowl. (a) KEGG pathway enrichment analysis of green peafowl.

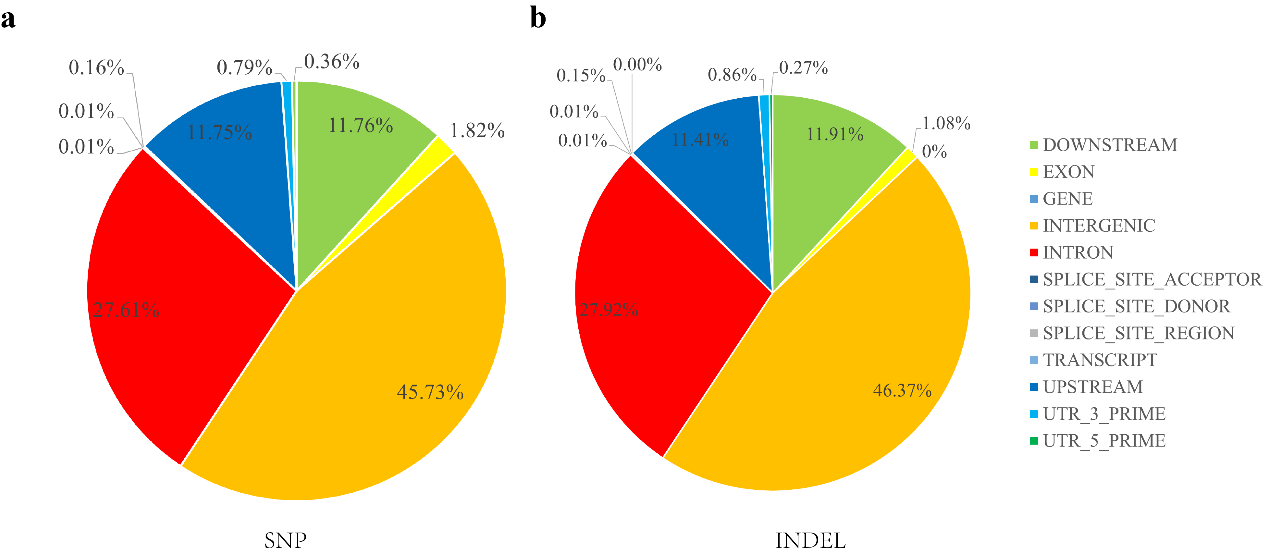

Supplementary Fig. S14. Variation annotation information of all peafowl individual. (a) SNPs annotation information. (b) Indels annotation information.

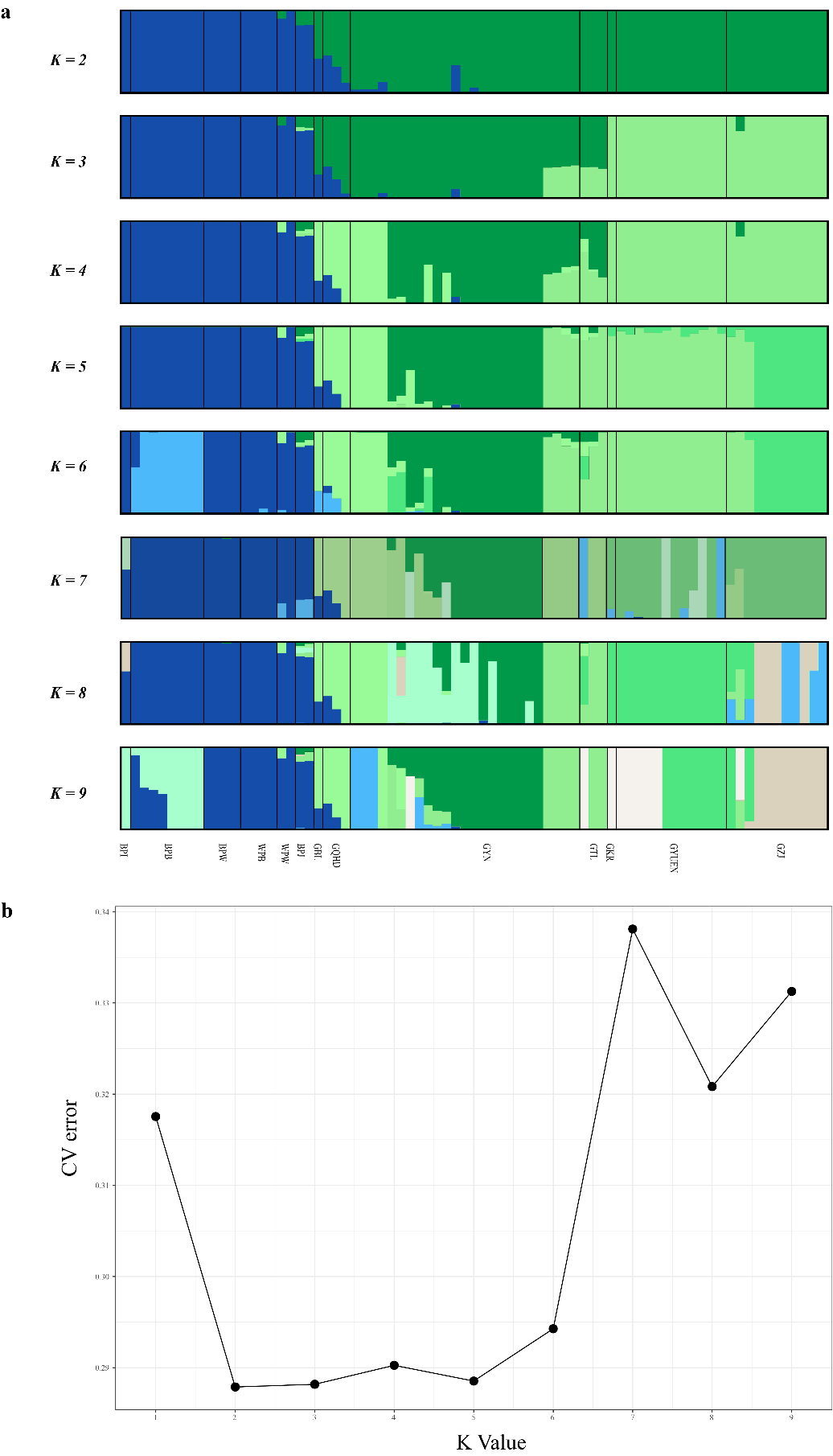

Supplementary Fig. S15. Group Structure of blue peafowl and green Peafowl. (a) ADMIXTURE clustering of core accessions from K=2 to 9. (b) Cross-validation error plot for the unsupervised ADMIXTURE analysis.

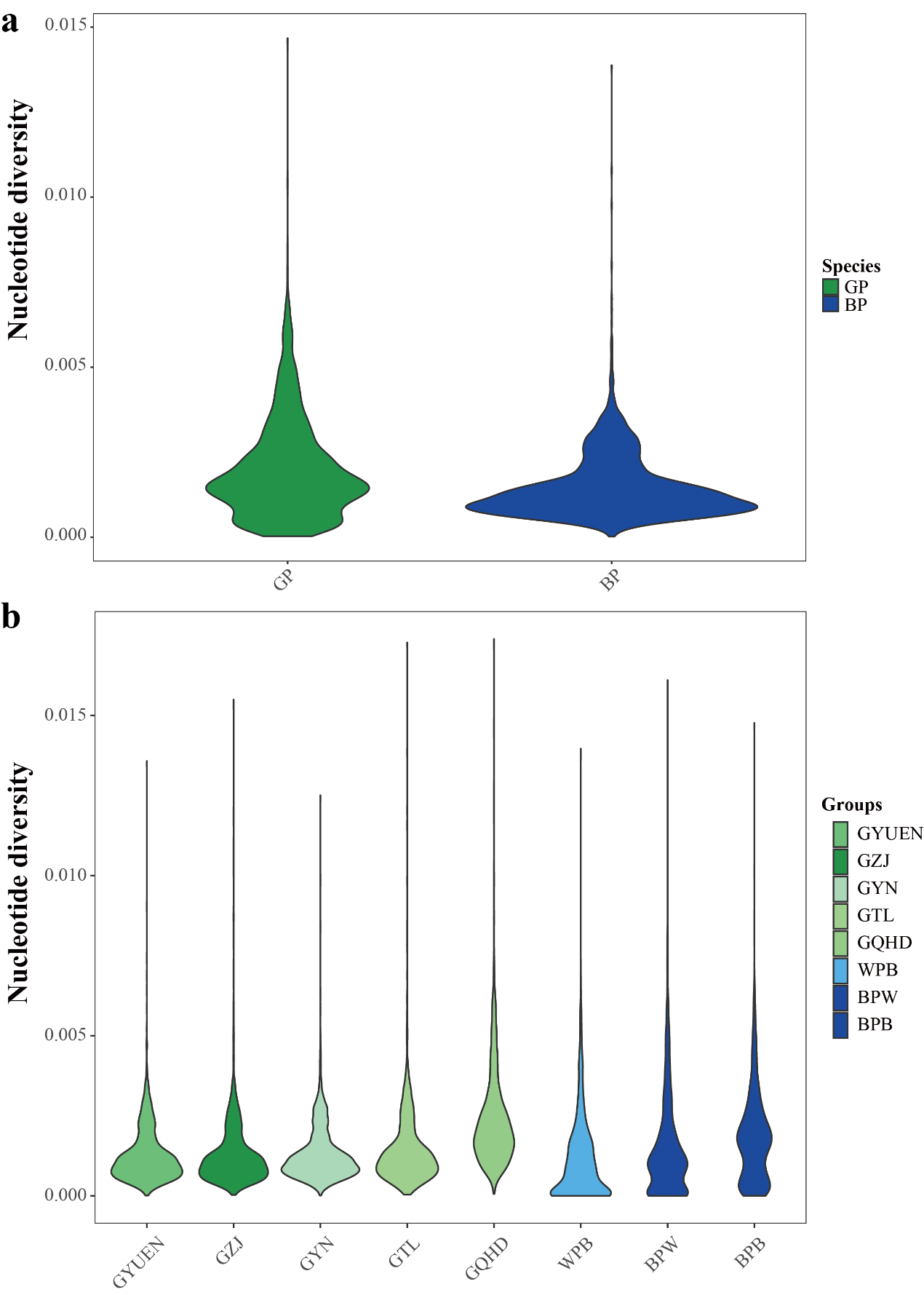

Supplementary Fig. S16. Nucleotide diversity of blue peafowl and green peafowl. (a) GP means green peafowl and BP means green. (b) Nucleotide diversity in peafowl groups with individuals greater than two.

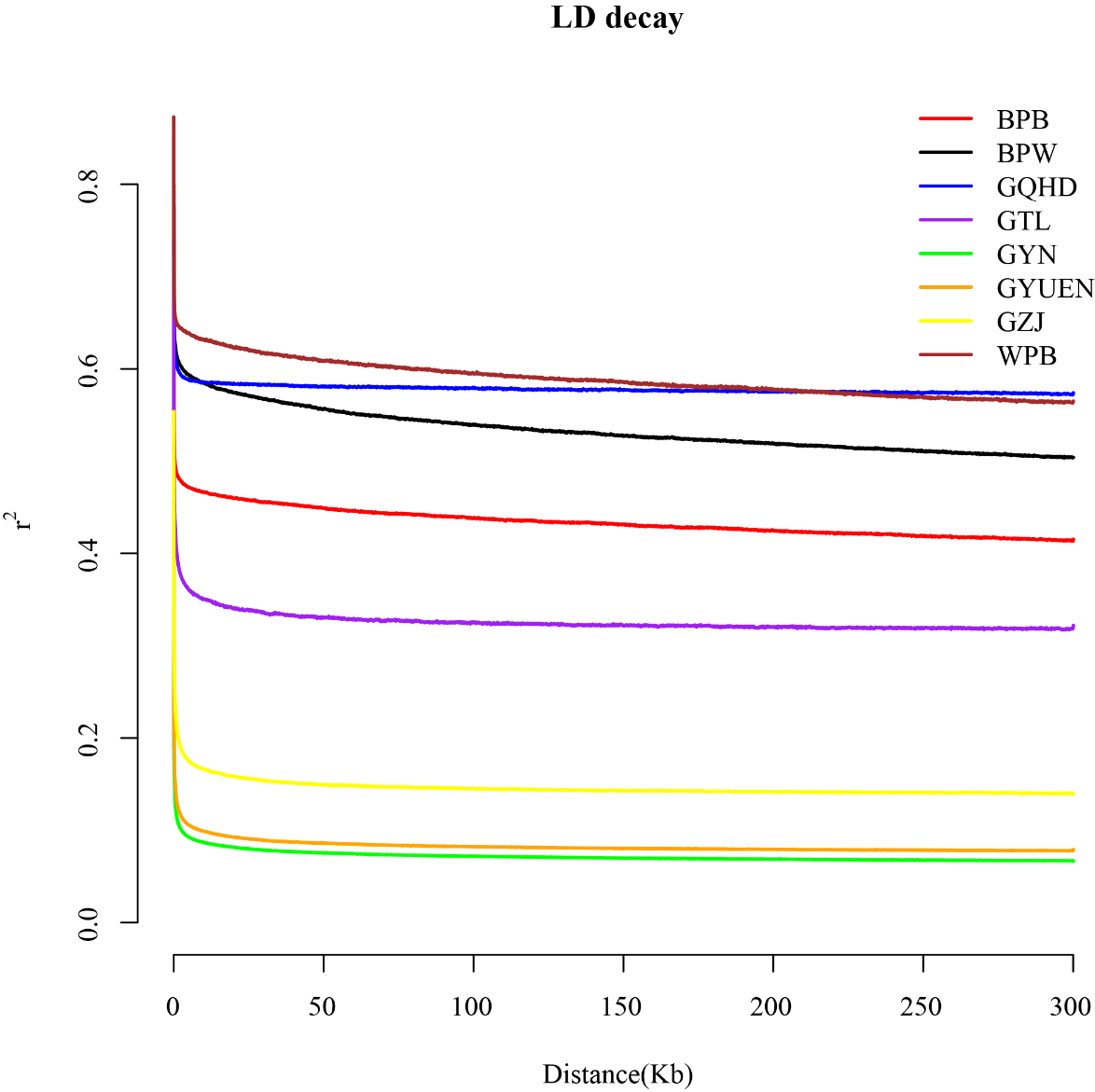

Supplementary Fig. S17. Linkage disequilibrium in peafowl groups with individuals greater than two.

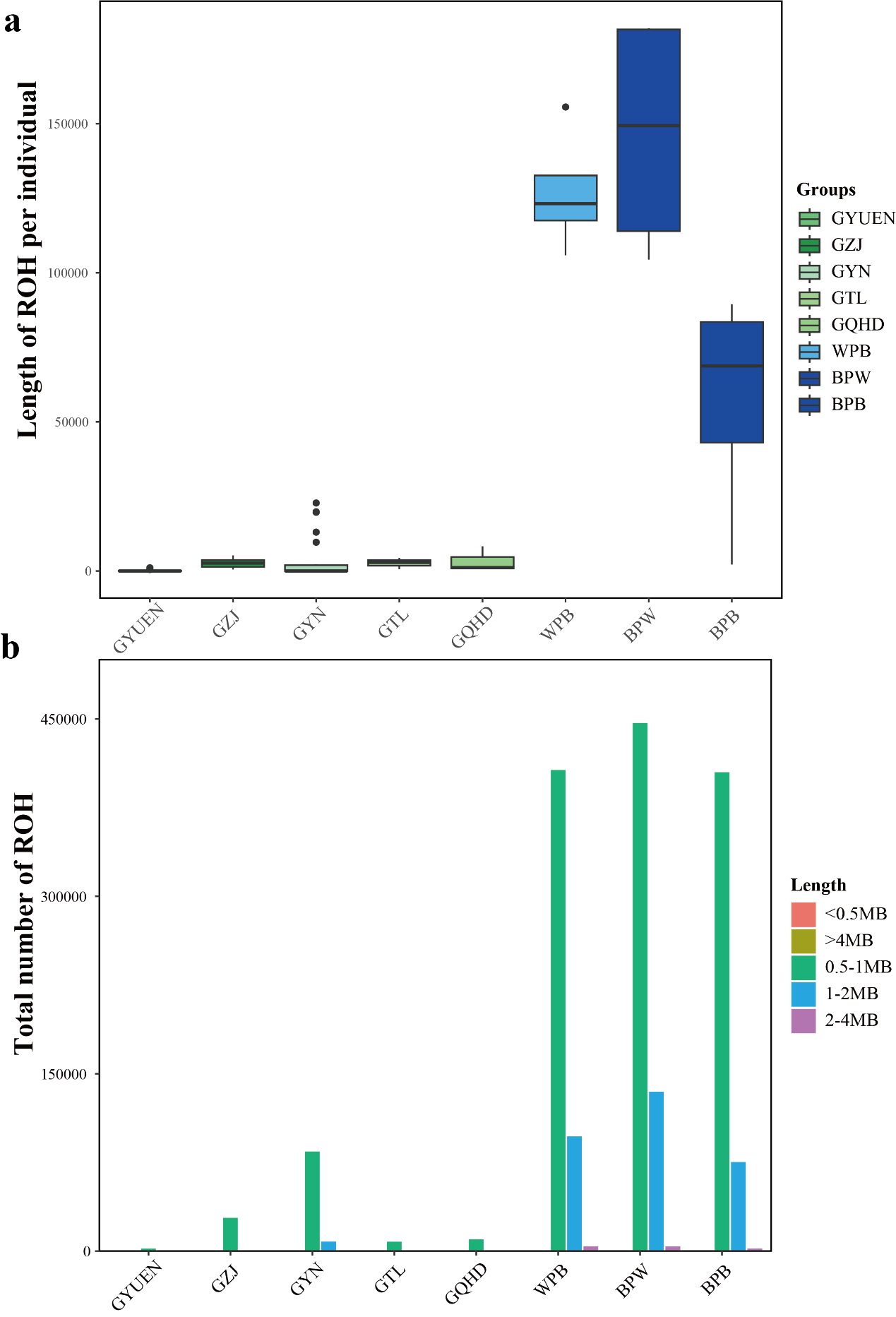

Supplementary Fig. S18. Runs of homozygosity (ROH) in peafowl groups with individuals greater than two. (a) The distribution of total number of ROH. (b) Box plot of ROH length in each peafowl group.

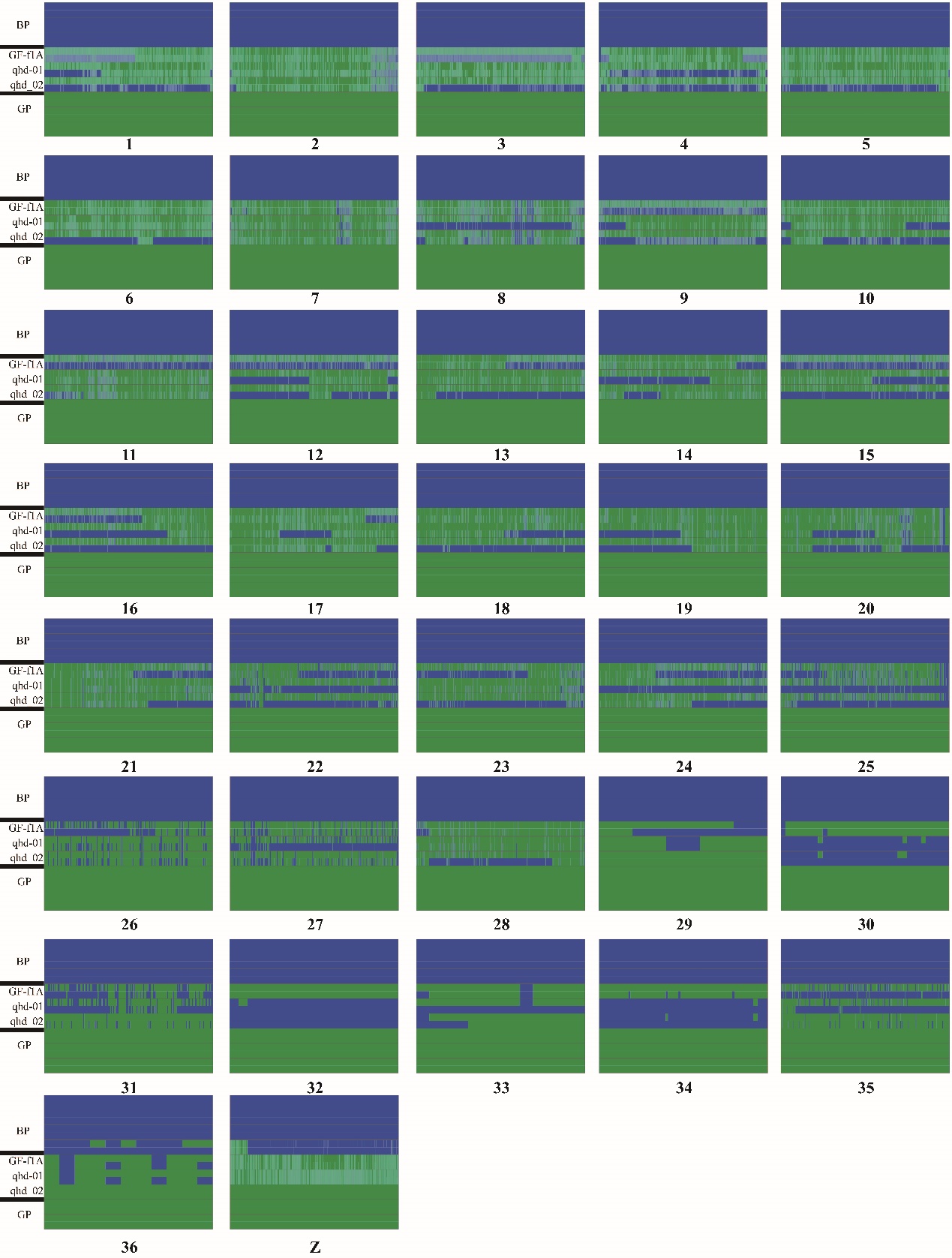

Supplementary Fig. S19. Each chromosome distribution of shared alleles SNPs between blue peafowl (blue) and green peafowl (green) in hybrid green peafowl individuals. Each row represents a diploid individual with two haplotypes stacked on top of one another, and three blue peafowl individuals and three green peafowl individuals were selected as controls.

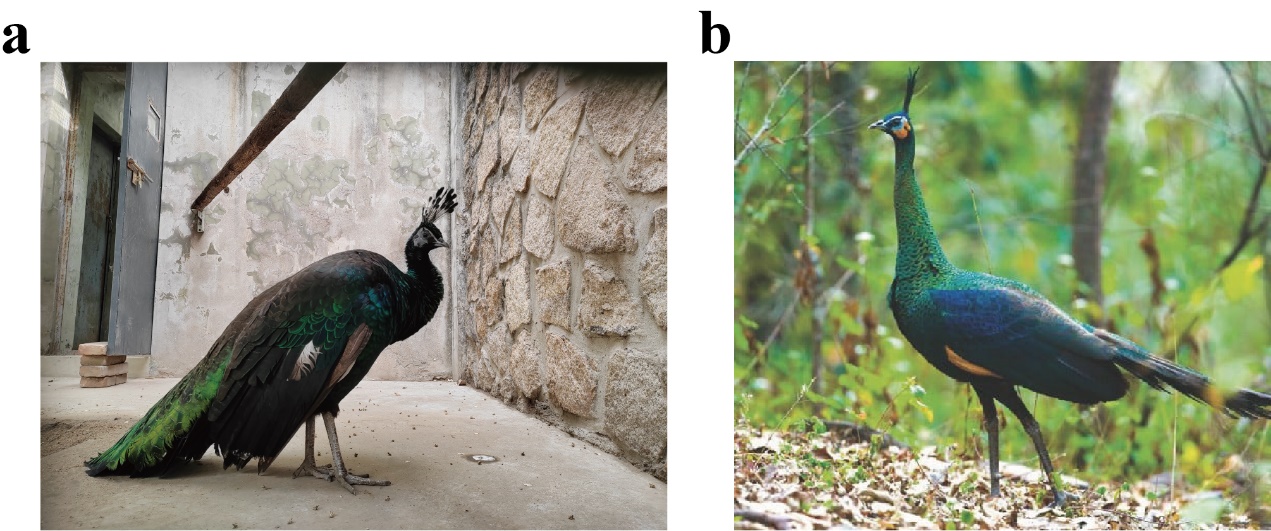

Supplementary Fig. S20. Comparison of hybrid individual green peafowl and purebred green peafowl. (a) The green peafowl sample shown in the picture is GF-f1A, which is a hybrid green peafowl. (b) The green peafowl sample shown in the picture is BP-5B, which is a purebred green peafowl. Two green peafowl individuals are very similar in appearance.

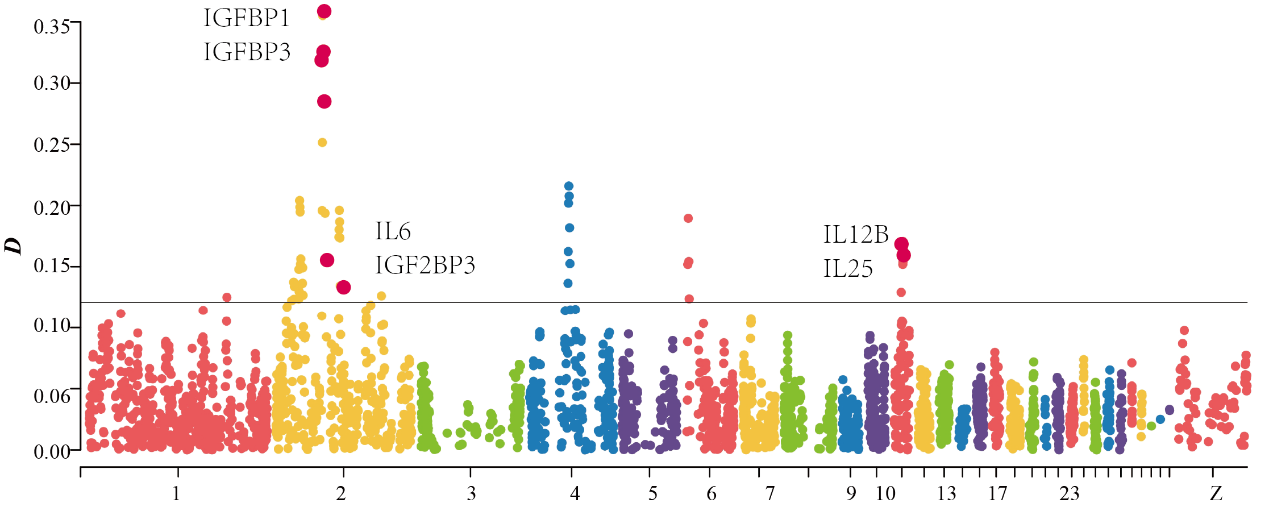

Supplementary Fig. S21. Manhattan plot of BPB blue peafowl group introgression with a window of 2500 SNPs and a step size of 500 SNPs (P1, P2, P3, O). P1 is BPW blue peafowl group, P2 is BPB blue peafowl group, P3 is GYN green peafowl group and O is chicken.

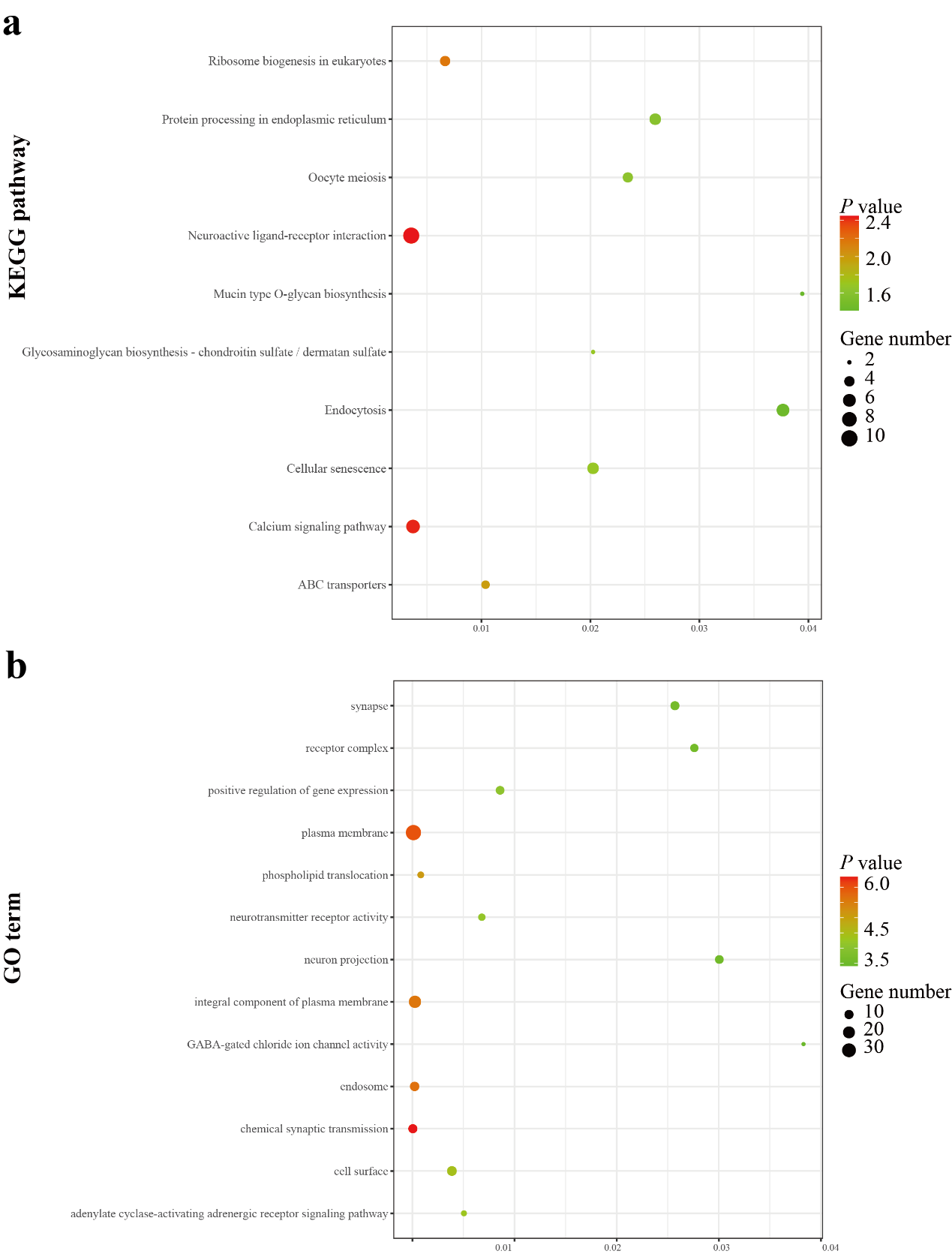

Supplementary Fig. S22. Enrichment analysis of green peafowl introgression regions in BPB blue peafowl group. (a) KEGG Pathway with *P* value < 0.05. (b) Go terms with *P* value < 0.05.

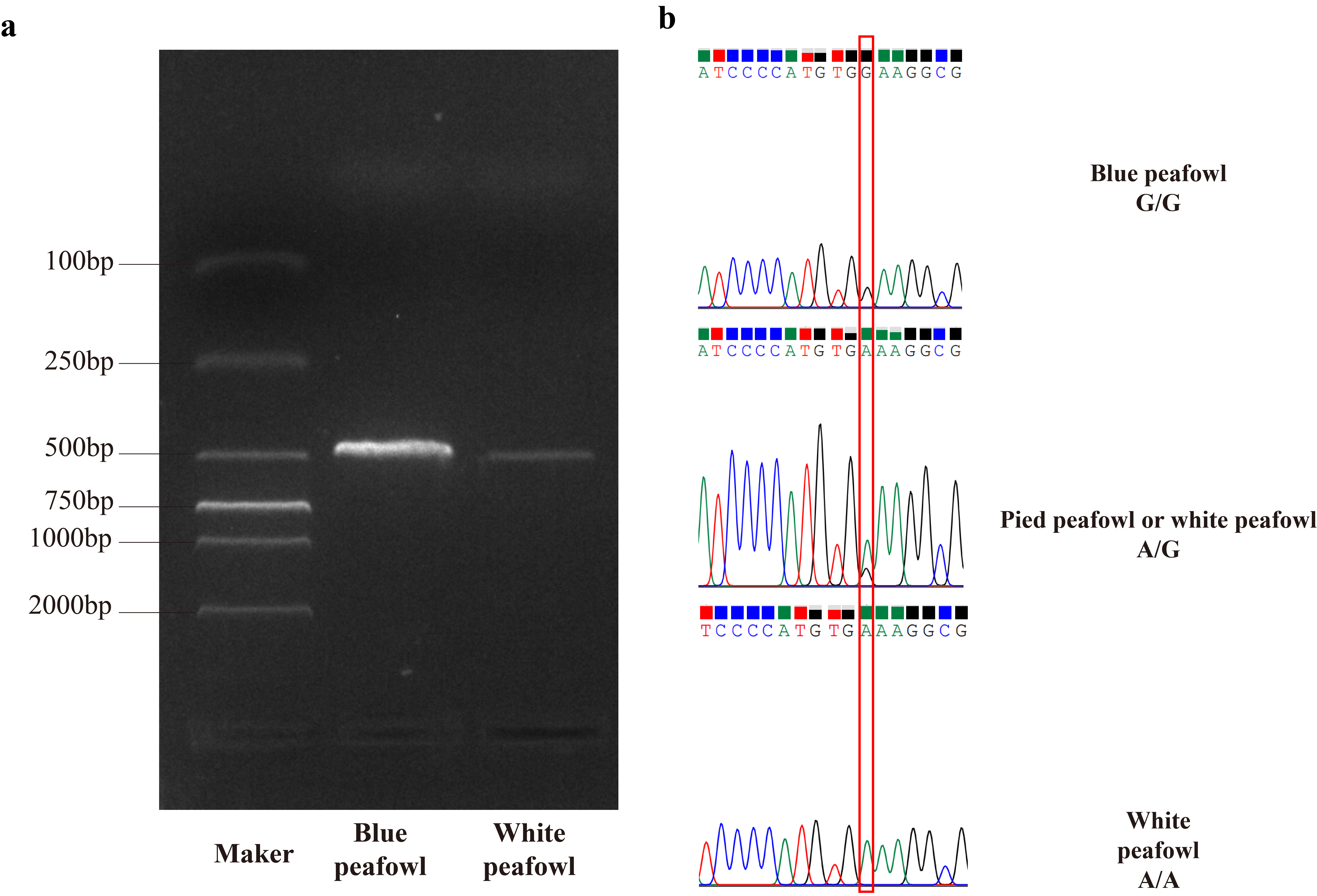

Supplementary Fig. S23. Caused mutation of the EDNRB2 gene locus (chr4: g.12583552 G>A) for the white plumage trait in white peafowls. (a) EDNRB2 gene fragment PCR amplified using primers (forward primer 5′- TGAAGAAGTGTAAGTCCCGCTG-3′ and reverse primer 5′-AGGTCTCGGTCCCAGTAGTT-3′). (b) Sanger sequencing displays the bases of mutation sites.

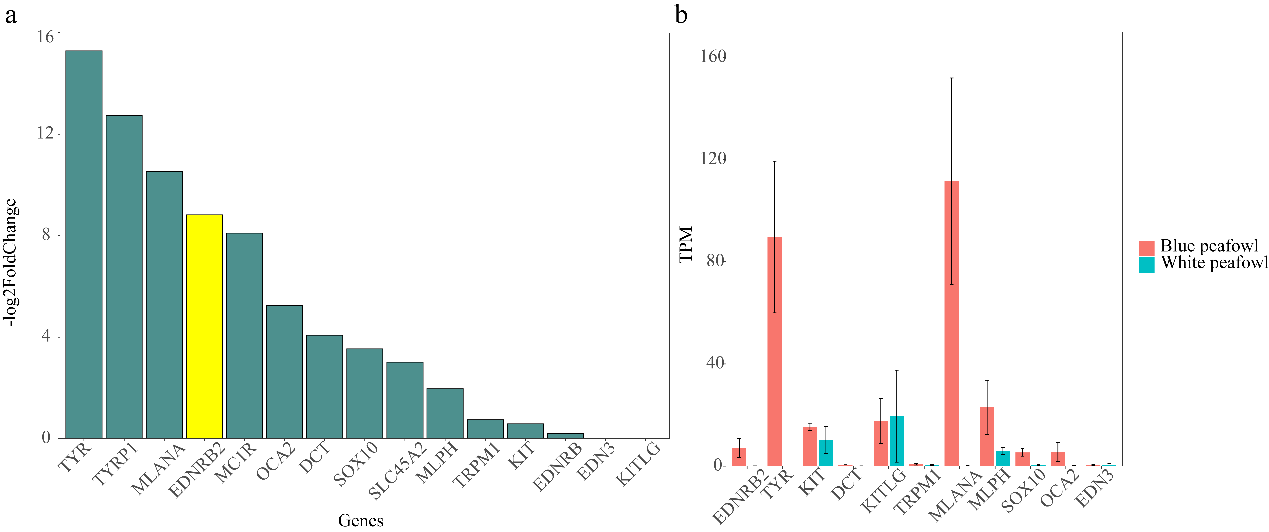

Supplementary Fig. S24. Bar plot of differential mRNA expression (log2-transformed fold change) and (Transcripts per million (TPM)) of pigmentation-related genes expressed in the plumage of white peafowls (n = 6) versus blue peafowls (n = 6).

Supplementary Table S1. Statistics of genome assembly data of blue peafowl (WP-1).

| Pair-end libraries | NCBI accession number | Reads Number | Total length of reads | Sequence coverage (×) | Source |
| --- | --- | --- | --- | --- | --- |
| PacBio HiFi（CCS） | - | 281,697 | 4,085,715,549bp | 29 | This study |
| Hi-C | - | 101,151,513,778 | 11,025,776,600 bp | 101 | This study |
| RNA-seq | SRR12578362 | 46,298,061 | 6.9G | - | Download |
|  | SRR12578363 | 51,306,082 | 7.7G | - | Download |
|  | SRR12578364 | 46,298,061 | 6.9G | - | Download |
|  | SRR12578365 | 48,173,671 | 7.2G | - | Download |
|  | SRR12578366 | 56,208,409 | 8.4G | - | Download |
|  | SRR12578367 | 47,936,094 | 7.2G | - | Download |
|  | SRR12578368 | 49,018,929 | 7.4G | - | Download |
|  | SRR12578369 | 46,368,286 | 7G | - | Download |
|  | SRR1797848 | 28,064,99 | 5.6G | - | Download |
|  | SRR1797849 | 19,796,449 | 4G | - | Download |
|  | SRR1797850 | 21,703,534 | 4.3G | - | Download |
|  | SRR1797859 | 35,627,018 | 7.1G | - | Download |
|  | SRR1797860 | 19,892,444 | 4G | - | Download |
|  | SRR1797861 | 23,556,090 | 4.7G | - | Download |
|  | SRR1797862 | 21,473,714 | 4.3G | - | Download |
|  | SRR1797863 | 23,354,828 | 4.7G | - | Download |
|  | SRR1797864 | 20,276,891 | 4.1G | - | Download |
|  | SRR1797865 | 22,855,972 | 4.6G | - | Download |
|  | SRR1797866 | 26,024,982 | 5.2G | - | Download |
|  | SRR1797868 | 18,628,156 | 3.7G | - | Download |
|  | SRR1797869 | 34,328,478 | 6.9G | - | Download |
|  | SRR1797870 | 21,140,602 | 4.2G | - | Download |
|  | SRR1797873 | 21,409,956 | 4.3G | - | Download |
|  | SRR1797874 | 21,907,777 | 4.4G | - | Download |
|  | SRR1797876 | 25,381,484 | 5.1G | - | Download |
|  | SRR1797877 | 19,984,406 | 4G | - | Download |
|  | SRR1797880 | 22,035,503 | 4.4G | - | Download |
|  | SRR1797882 | 15,830,797 | 3.2G | - | Download |

Supplementary Table S2. Summary of de novo genome assembly of blue peafowl

| Items | Value |
| --- | --- |
| Number of scaffolds | 38 |
| Number of contigs | 389 |
| Length of the longest scaffold | 19,996,252 |
| Genome scaffolds size | 1,041,782,769 |
| Genome contig size | 1,134,053,066 |
| Total of N | 182000 |
| Rate of GC | 0.4253 |
| Scaffold N50 | 94,977,930 |
| Contig N50 | 30,610,815 |
| Scaffold N90 | 12,832,000 |
| Contig N90 | 3,121,499 |
| Number of sequences >= 1kb | 38 |
| Number of sequences >= 2kb | 38 |
| Number of sequences >= 3kb | 38 |
| Number of protein-coding genes | 37,401 |
| Number of protein-coding mRNAs | 37,401 |
| Number of exons | 302,149 |
| Number of coding regions | 282,849 |

Supplementary Table S3. Assembly assessment of completeness using BUSCOs.

| BUSCO categories | Gene number | Percentage (%) |
| --- | --- | --- |
| Complete BUSCOs | 8,099 | 97.10% |
| Complete and Single-copy BUSCOs | 8,054 | 96.60% |
| Complete and duplicated BUSCOs | 45 | 0.50% |
| Fragmented BUSCOs | 32 | 0.40% |
| Missing BUSCOs | 207 | 2.50% |

Supplementary Table S4. Statistics of repeats in our assembled genome.

| Type | Length (bp) | % of genome |
| --- | --- | --- |
| SINEs | 519129 | 0.05 |
| Penelope | 122535 | 0.01 |
| LINEs | 77,192,339 | 7.41 |
| LTR | 31,100,097 | 2.99 |
| DNA transposons | 8,770,541 | 0.84 |
| Total | 139,353,537 | 13.38 |

Supplementary Table S5. Statistics of non-coding RNAs in the assembly of peafowl.

| Type | | Copy number | Average length(bp) | Total length(bp) | % of genome |
| --- | --- | --- | --- | --- | --- |
| miRNA | | 354 | 102.23 | 36,190 | 0.003457 |
| tRNA | | 308 | 76.22 | 23,477 | 0.002243 |
| rRNA | rRNA | 151 | 151.53 | 22,881 | 0.002186 |
|  | 18S | 23 | 235.96 | 5,427 | 0.000518 |
|  | 28S | 105 | 142.03 | 14,913 | 0.001425 |
|  | 5.8S | 0 | 0 | 0 | 0 |
|  | 5S | 23 | 110.48 | 2,541 | 0.000243 |
| snRNA | snRNA | 334 | 128.77 | 43,010 | 0.004109 |
|  | CD-box | 131 | 101.06 | 13,239 | 0.001265 |
|  | HACA-box | 81 | 144.16 | 11,677 | 0.001116 |
|  | splicing | 101 | 142.1 | 14,352 | 0.001371 |

Supplementary Table S17. Genotype distribution of the short deletion at chromosome 4 (chr4: g.12583552 G>A) in Blue and White peafowls. The term “Photo” means that a photographic record of the feather color was taken; “record” means feather color was not photographed but annotated. “Genotype” means the genotype of the SNP in the EDNRB2 gene, which is associated with feather color in peafowl.

| Sample | Photo/Record | Genotype | Feather color phenotype |
| --- | --- | --- | --- |
| WP-N-1 | Photo | A/A | White |
| WP-N-2 | Photo | A/A | White |
| WP-N-3 | Photo | A/A | White |
| WP-N-5 | Photo | A/A | White |
| WP-N-6 | Photo | A/A | White |
| WP-N-7 | Photo | A/A | White |
| WP-1B | Photo | A/A | White |
| WP-2B | Photo | A/A | White |
| WP-3B | Photo | A/A | White |
| WP-4B | Photo | A/A | White |
| WP-M1A | Photo | A/A | White |
| BP-N-1 | Photo | G/G | Blue |
| BP-N-2 | Photo | G/G | Blue |
| BP-N-3 | Photo | G/G | Blue |
| BP-N-4 | Photo | G/G | Blue |
| BP-N-5 | Photo | G/G | Blue |
| BP-N-6 | Photo | G/G | Blue |
| BP-N-7 | Photo | G/G | Blue |
| BP-N-8 | Photo | G/G | Blue |
| BP-N-9 | Photo | G/G | Blue |
| BP-N-10 | Photo | G/G | Blue |
| BP-N-11 | Photo | G/G | Blue |
| BP-N-13 | Photo | G/G | Blue |
| BP-N-15 | Photo | G/G | Blue |
| BP-N-16 | Photo | G/G | Blue |
| BP-N-18 | Photo | G/G | Blue |
| BP-N-19 | Photo | G/G | Blue |
| BP-1B | Photo | G/G | Blue |
| BP-2B | Record | A/G | Blue |
| BP-3B | Record | A/G | Blue |
| BP-4B | Record | A/G | Blue |
| BP-5B | Record | A/G | Blue |
| BP-6B | Photo | G/G | Blue |
| BP-7B | Photo | G/G | Blue |
| BP-8B | Photo | G/G | Blue |
| BP-M1A | Record | G/G | Blue |
| BP-M2A | Record | G/G | Blue |
| BP-M3A | Record | G/G | Blue |
| BP-M5A | Record | G/G | Blue |
| PC-Run1 | Record | G/G | Blue |
| BP-N-20 | Photo | A/G | Blue and White |
